# Pervasive Transcriptome Interactions of Protein-Targeted Drugs

**DOI:** 10.1101/2022.07.18.500496

**Authors:** Linglan Fang, Willem A. Velema, Yujeong Lee, Xiao Lu, Michael G. Mohsen, Anna M. Kietrys, Eric T. Kool

## Abstract

The off-target toxicity of drugs targeted to proteins imparts substantial health and economic costs. Proteome interaction studies can reveal off-target effects with unintended proteins; however, little attention has been paid to intracellular RNAs as potential off targets that may contribute to toxicity. To begin to assess this, we developed a reactivity-based RNA profiling (RBRP) methodology, and applied it to uncover transcriptome interactions of a set of FDA-approved small-molecule drugs *in vivo*. We show that these protein-targeted drugs pervasively interact with the human transcriptome and can exert unintended biological effects on RNA function. In addition, we show that many off-target interactions occur at RNA loci associated with protein binding and structural changes, allowing us to generate hypotheses to infer the biological consequences of RNA off-target binding. The results suggest that rigorous characterization of drugs’ transcriptome interactions may help assess target specificity and potentially avoid toxicity and clinical failures.

---

Dose-limiting toxicity is a routinely encountered problem in therapeutic drug development that is often discovered in clinical trials^1^. This is both detrimental to positive human health outcomes and costly in capital and human effort. Even for approved drugs, dose-limiting toxicity results in adverse outcomes and post-marketing drug withdrawal^2^. These drug toxicities are commonly attributed to off-target binding to unintended cellular proteins^3^. Given RNA’s broad bioregulatory roles in human physiology and the structural resemblance of protein-targeted drugs with known RNA-binding molecules^4,5^ (**Fig. 1a-c**), we hypothesize that many protein-targeted FDA-approved small-molecule drugs can interact with the human transcriptome (and RNA-protein interfaces) *in vivo* and may confer serious toxicity in patients as a result.

**Fig. 1.**
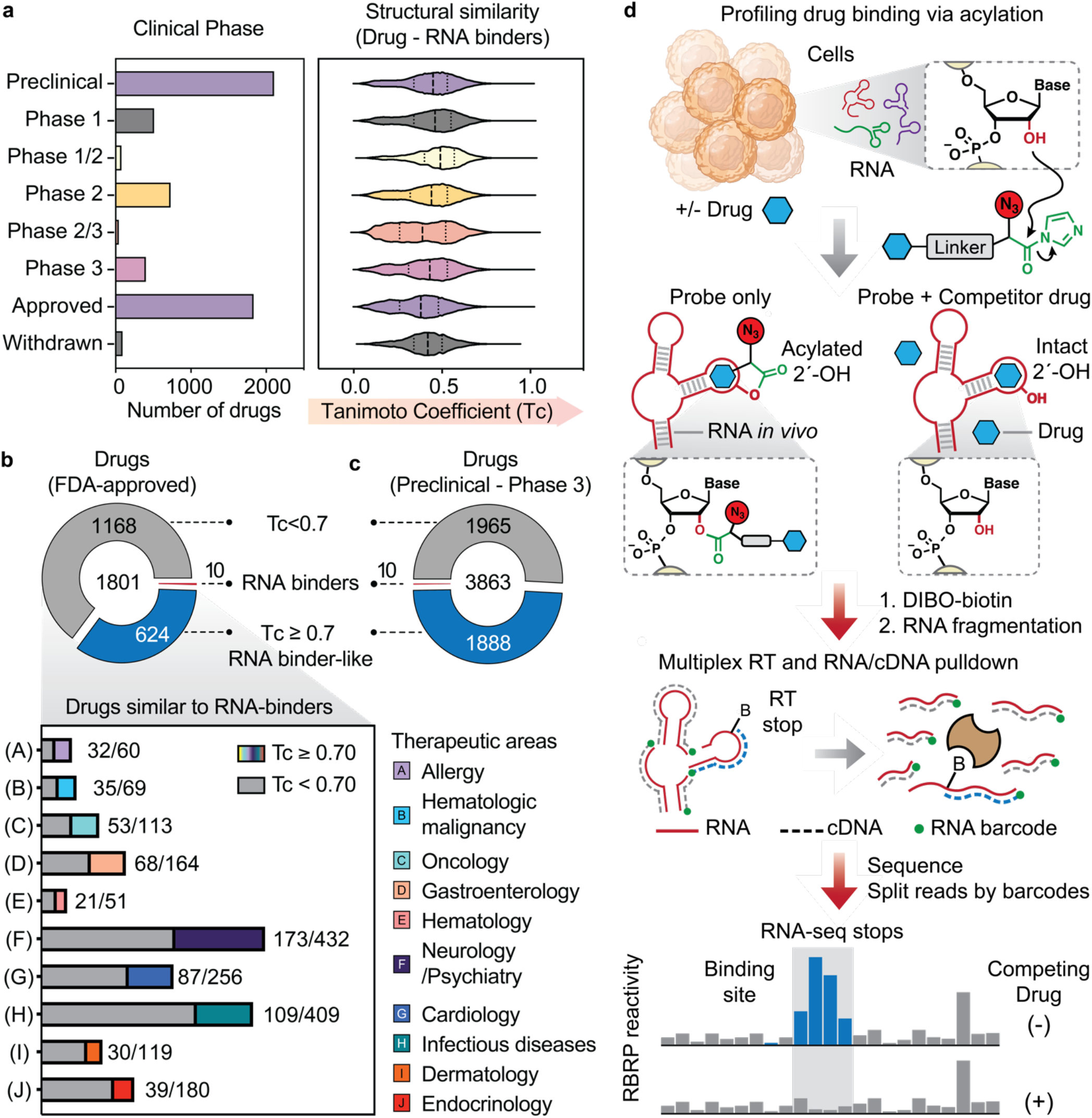
RBRP decodes transcriptome-interaction of protein-targeted small-molecule drugs *in vivo*. **a**, Structure key-based 2D Tanimoto coefficient (Tc) characterizes pairwise structural similarity between 131 reported small-molecule RNA binders^5^ and 6043 drugs at various clinical phases. Number of small-molecule drugs at each clinical phase^24^ (left panel). **b**, Fractions of FDA-approved drugs that structurally resemble at least one known RNA binder in the top 10 therapeutic areas. **c**, Fractions of drugs from preclinical to phase 3 stages that structurally resemble at least one known RNA binder as measured by Tanimoto coefficient. See **Supplementary file S1** for data. **d**, Schemes for comparative Reactivity-Based RNA Profiling (RBRP) to identify transcriptome-interaction of smallmolecule drugs. Acylating analog of drug binds rapidly at specific RNA sites, promoting local acylation, which is analyzed by reverse transcription (RT) stops via multiplexed RNA sequencing (RNA-Seq). Pulldown of acylated RNAs greatly enhances signal/noise, and competition with unmodified drugs reveals specific sites. Binding sites exhibit statistically significant differences in RBRP reactivities.

Studies provide support for this notion; for example, Disney *et al*. showed that the investigational drug Dovitinib engages pre-miR-21 in triple-negative breast cancer (TNBC) cells, which guided the development of re-purposed Dovitinib-RIBOTAC to target pre-miR-21^6^. A parallel microarray-based library-versus-library screening *in vitro* indicated that kinase inhibitors can bind RNA and topoisomerase inhibitors can engage RNAs such as pre-miR-21^7^. A computation-based method (Inforna) also suggested that protein-targeted topoisomerase inhibitors can bind this same pre-miRNA^7^. These prior studies have led to the postulation that failed drug toxicity may be due in part to off-target RNA interactions^8^. Our analysis suggests that nearly 35% of all small-molecule FDA-approved drugs chemically resemble at least one known RNA-binding molecule (Tanimoto coefficient^9^ ≥0.7) (**Fig. 1a-b** and **Supplementary file S1**), and at least ten approved drugs are already documented as RNA binders *in vitro* (**Supplementary file S2**)^4,5^. Such chemical similarity with RNA-binding molecules also extends to drugs from preclinical studies and all phases in clinical trials (**Fig. 1c**).

Investigation of small molecule-RNA interactions has been carried out recently via photocrosslinking (diazirine)^6,10–12^, alkylation (chlorambucil)^13,14^, in-line probing^15^, and SHAPE^16^, and these have emerged as useful tools to identify RNA-ligand interactions *in vitro* and recently in cultured cells^17^. Although elegant, application of these methods *in vivo* is potentially limited by transcriptional effects during prolonged probe treatment^18^, cellular damage by ultraviolet irradiation^19^, and nucleobase biases^20–22^. To address these issues and potentially complement prior methods that identify RNA-ligand interactions, we developed cell-permeable RNA 2’-hydroxyl (2’-OH) acylating analogs of drugs that permit *in vivo* analysis of RNA-drug contacts in a sequence-independent manner. This reactivity-based RNA profiling (RBRP) methodology allows us to decode transcriptome interactions of drugs via deep sequencing and bioinformatics (**Fig. 1d**).

In this study, we employed RBRP to map *in vivo* transcriptome interactions of representative protein-targeted drugs in human cells. We provide evidence that RNA off-targets exist for FDA-approved protein-targeted drugs in cells. These transcriptome interactions are largely determined by the chemical structure of small-molecule drugs. We further demonstrate that drug engagement with RNA off-targets can exhibit target-associated cellular outcomes; for example, drug engagement with an RNA G-quadruplex in YBX1 5’ UTR inhibits the translation of genes encoded downstream of the UTR. Further, drug engagement with snoRNA sequences modulates downstream 2’-OH methylation of ribosomal RNAs. Moreover, RBRP enables mapping of the binding site across ~16,000 RNAs containing a poly(A) tail with nucleotide-level precision, and reveals intricate interplay of RNA-drug binding, RNA-binding proteins (RBP), and RNA structural accessibility and dynamics.

## Results and discussion

### RBRP decodes transcriptome interactions of protein-targeted small-molecule drugs *in vivo*

Our profiling methodology involves the use of cell-permeable RNA acylation probes (employing acylimidazole-substituted linkers to react at RNA 2’-OH groups) to evaluate and quantify the tendency of drugs to bind cellular RNAs (**Fig. 1d**). The binding of an acylimidazole conjugate of a drug to a structured RNA or protein-RNA interface should lead to enrichment of acylated 2’-OH near the drug binding sites. We identify this binding-promoted acylation by modifying *in vivo* RNA mapping protocols^23^, analyzing messenger RNAs (mRNAs) and noncoding RNAs (ncRNAs) by poly(A) pulldown and deep sequencing the resulting libraries at high depth (>11 million reads per replicate). Specifically, RNA drug-binding sites are enriched with acylated 2’-OH groups, which cause reverse transcriptase (RT) to stop. We designed, optimized, and validated a workflow to enrich these stops over random RNA breaks, random site acylation by acylimidazole warheads, nonspecific binding events by biotin-mediated pulldown,^23^ and potential transcriptional changes upon treatment with the unmodified drug (**Extended Data Fig. 1**). This comparative workflow allows us to locate and quantify proximal binding sequences within the desired cellular RNA population; only sites that exhibit competition with the unmodified drug are scored as authentic drug-binding sites.

### RBRP reveals the transcriptome interactions of Levofloxacin (Lev)

We prototyped *in vivo* RBRP experiments with an acylimidazole conjugate of the small-molecule drug Levofloxacin (Lev), containing an azido “click” handle, in human embryonic kidney cells HEK293 (**Fig. 2a-c**). Levofloxacin (Lev) is a member of widely prescribed fluoroquinolone antibiotics known to cause neuropathy, fatigue, and depression in patients^25^. Its structure suggests possible affinity for folded RNAs, given its fused aromatic rings and positive charge, as well as its close structural similarity with known RNA binders (**Fig. 2a**, right panel). To test this possibility, we treated HEK293 cells with the acylating analog of Lev (Lev-AI, **Fig. 2c**) at a clinically relevant concentration (50 μM)^26^ for 30 minutes in the absence or presence of competing unmodified Lev in excess. We also performed RNA-seq experiments with HEK293 cells that were treated only with the competing drug or vehicle control (DMSO) (**Extended Data Fig. 2a-b**). After confirming that treatment with excess competing drug did not significantly alter the expression level of most cellular transcripts, we performed RBRP for acylating probe-treated polyadenylated transcripts. Deep sequencing results from RBRP showed very strong correlation of transcript expression value (RPKM) between two biological replicates (Pearson correlation *r* = 1.00) (**Extended Data Fig. 3a-c**). The concordance of RT-stop frequencies also remained high for most transcripts with read depth higher than the optimized cutoff value (200) (**Extended Data Fig. 3d-f**). Thus, we performed bioinformatics analysis at the regions with read depth high enough to provide strong concordance among replicates. Specifically, we used a modified icSHAPE pipeline^23,27^ (read depth= 200 as threshold) to quantify the yield of 2’-OH acylation at each nucleotide transcriptome-wide and generated 0.8 billion measurements *in vivo* (**Extended Data Fig. 1**). To remove random site acylation by the acylimidazole warheads and non-specific binding events by biotin-mediated pulldown, we subtracted the yield of 2’-OH acylation by the linker alone (Linker-AI) from that by Lev-AI. We also embedded RNA-seq experiments to account for potentially changed transcript abundance. Finally, RBRP reactivities at each site were calculated as the surplus yield of 2’-OH acylation for experiments performed in the absence over the presence of excess competing drug.

**Fig. 2.**
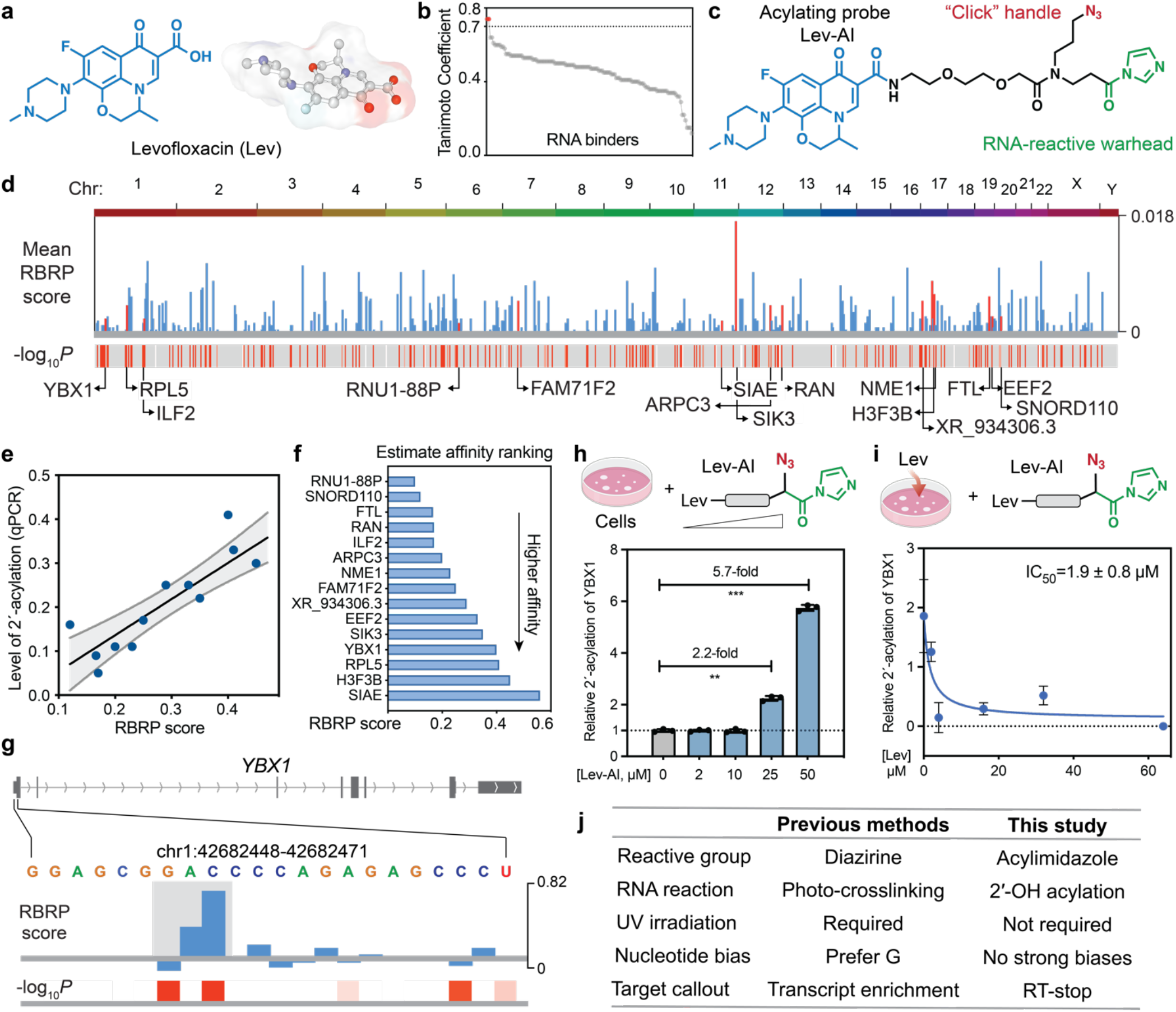
RBRP reveals transcriptome-interaction of Levofloxacin (Lev). **a.** Chemical structure of Levofloxacin (Lev) colored in blue. **b**, Dot plot showing the pairwise 2D Tanimoto coefficient (Tc) between Lev and known RNA binders. **c**, A modular design of acylating probe (Lev-AI) to identify transcriptome-interaction of Lev. The drug part is colored in blue, linker in black, “click” handle in red, and RNA-reactive warhead in green. **d**, UCSC tracks showing the mean RBRP scores (y-axis) in live HEK293 cells along the human chromosomes (*top*) and −Log_2_*P*-value of Welch *t*-test (*bottom*) of Lev-AI probe. Mean RBRP reactivity scores are scaled from 0 to 0.018. **e**, Linear regression analysis showing high concordance between RBRP score and 2’-acylation determined by independent qPCR for most targeted sites. Pearson’s correlation *r* = 0.90. **f**, RBRP scores enables estimated ranking of binding affinity of RNA interactions of Lev. **g**, UCSC tracks showing RBRP score at an RNA locus in YBX1 mRNA (*top*) and −Log_2_*P*-value of Welch *t*-test (*bottom*), suggesting binding of Lev. **h**, Lev-AI increases 2’-OH acylation in a dose-dependent manner at a binding site in YBX1 mRNA. Data shown are mean ± s.e.m., *n=3* independent biological replicates. **i**, Unmodified Lev dose-dependently reduces 2’-OH acylation at YBX1 binding site, with an IC50=1.9 ± 0.8 μM. Data shown are mean ± s.e.m., *n*=3 independent biological replicates. **j**, Comparing RBRP with previously reported diazirine-based methods for detecting RNA interactions of small molecules. Statistical significance was calculated with two-tailed Student’s *t*-tests: \*\**P*<0.01, \*\*\**P*<0.001, \*\*\*\**P*<0.0001.

To stringently eliminate the low-confidence transcriptome interactions, we validated all the putative drug binding sites with low-throughput RT-qPCR experiments (**Extended Data Fig. 4**). Deep sequencing results from RBRP showed reasonably strong correlation with RT-qPCR (Pearson correlation *r* = 0.90) for most putative binding sites, where significant RNA off targets are defined as nucleotides with RBRP score ≥ 0.12 (**Fig. 2e**). In addition, we computed the two-tailed *p*-value of Welch’s *t*-test comparing RBRP scores at each nucleotide transcriptome-wide in ± competitor experiments to further guide the calling of drug binding sites with high stringency.

These comparative RBRP data allowed us to identify probe-promoted, Lev-competable 2’-OH acylation as imprints of selective RNA-ligand interactions. Approximately 15 selective targets were identified in ~16,000 RNAs of HEK293 cells, all of which were independently confirmed by RT-qPCR (**Extended Data Fig. 4**). Because RBRP scores generally reflect level of competition by unmodified drug (**Fig. 2e**), we can approximately rank the binding affinity of targets based on their RBRP score (**Fig. 2f**). We found that RNA off-targets of Lev were enriched for exons of mature mRNA (**Supplementary file S3**). For example, Lev engaged with the open reading frames (ORF) of an mRNA that encodes Interleukin Enhancer Binding Factor 2 (ILF2). RBRP also showed that Lev targets non-coding RNAs (ncRNA) such as small nucleolar RNA C/D box 110 (SNORD110) and a U1 small nuclear 88 pseudogene RNA (RNU1-88P), which are polyadenylated within their annotated 3’ ends in HEK293 cells^30^ (**Supplementary Fig. 1**). These observations provide evidence for off-target transcriptome interactions of Levofloxacin in living human cells.

In addition, RBRP allowed us to estimate the binding affinity of RNA interactions of small-molecule drugs. For example, RBRP indicated that Lev targets the 5’ UTR of an mRNA that encodes the Y-box binding protein 1 (YBX1) (chr1:42,682,457-42,682,458) (**Fig. 2g**). With RT-qPCR, we showed dose-dependent 2’-OH acylation within the YBX1 5’ UTR in HEK293 cells (**Fig. 2h**). Pretreatment of HEK293 cells with increasing concentrations of unmodified Lev followed by treatment with Lev-AI demonstrated a dose-dependent reduction of 2’-OH acylation at the YBX1 5’ UTR site, with a clinically relevant IC_50_ value of 1.9 ± 0.8 μM (**Fig. 2i**).

To gather more evidence for transcriptome interactions of Lev and further evaluate the performance of RBRP, we directly compared RBRP to an orthogonal method that utilizes diazirine-conjugated probes to identify RNA *interactions*^6,12,31,32^. We constructed a photo-crosslinking analog of Lev (Lev-diazirine) which structurally resembles that of Lev-AI (**Extended Data Fig. 5a**). We conducted profiling experiments with Lev-diazirine in HEK293 cells with or without excess competitor drug (**Extended Data Fig. 5b)**, and photo-crosslinked RNAs were enriched following a reported protocol^6^. We then analyzed these crosslinked RNAs with deep sequencing and identified putative RNA hits with the enrichment of RNA transcripts in the absence or presence of excess competing drug. Profiling with Lev-diazirine identified 14 putative RNA interactions (**Extended Data Fig. 5c-d)**. Five “hits” discovered by RBRP were orthogonally confirmed by diazirine-based profiling experiments, with another four hits by RBRP less confidently validated by Lev-diazirine (**Extended Data Fig. 5e**). Both probes also identified additional sets of unique off-target interactions, suggesting complementarity between the two methods. Notably, the nucleotide composition in the RBRP sequencing libraries was found to be distributed almost evenly across all four nucleobases (**Extended Data Fig. 5f**). This is in contrast with diazirine-based profiling methods that display pronounced preference toward guanine nucleobase^20–22^. Taken together, the data highlight the viability of RBRP for profiling ligand-RNA interactions.

### Structural fingerprints of small-molecule drugs determine their transcriptome interactions

Existing protein-targeted drugs differ greatly in their physicochemical and molecular recognition features, covering a broad multidimensional chemical space (**Fig. 3a** and **Extended Data Fig.6**). To explore whether transcriptome-interaction of protein-targeted drugs extends beyond Levofloxacin, we next investigated two additional compounds that are structurally distinct from Lev and that have strongly divergent protein targets.

**Fig. 3.**
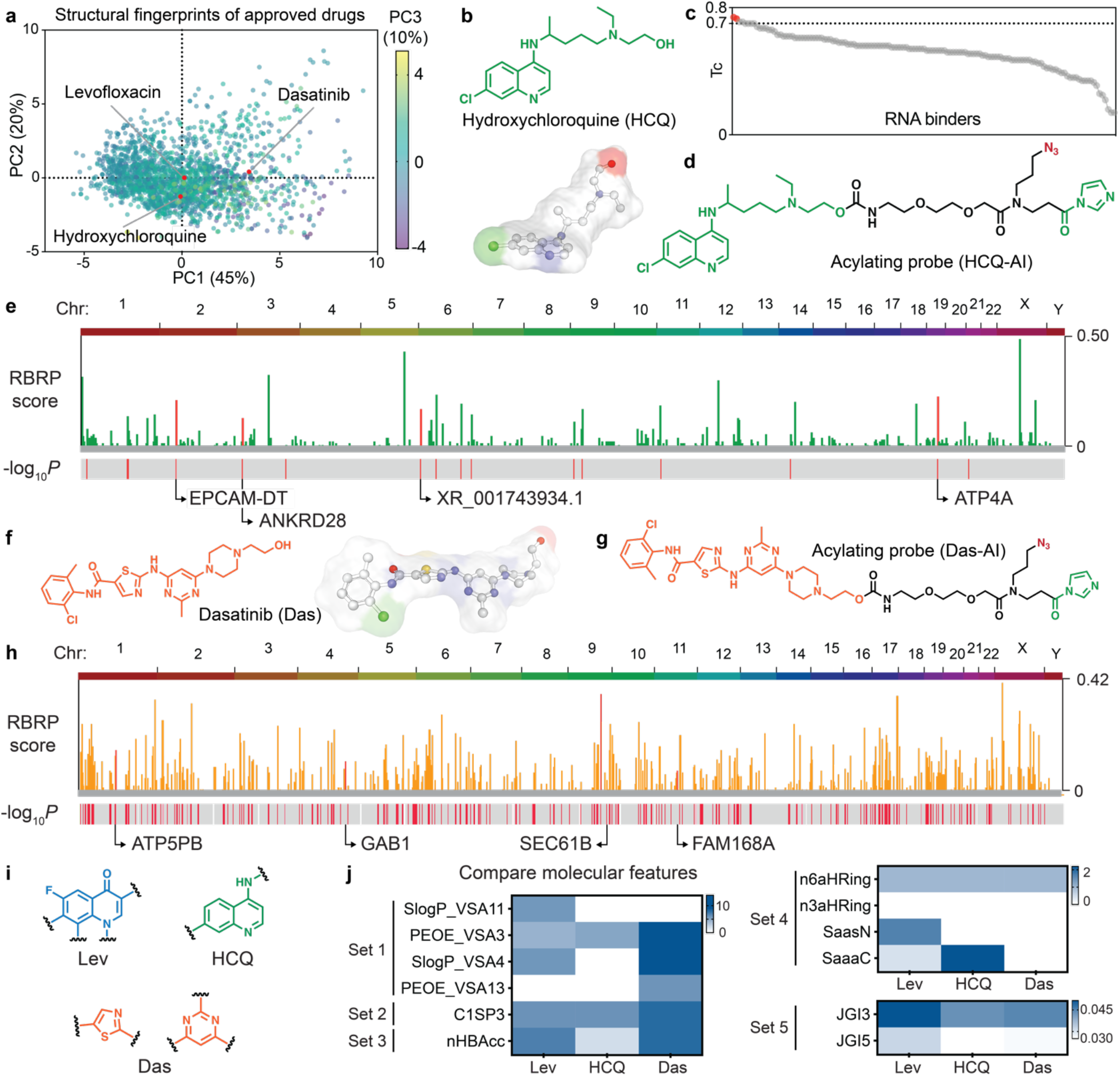
Structural fingerprints of small-molecule drugs determine their transcriptome interactions *in vivo*. **a**, Principal component analysis (PCA) describing the structural variance within the selected FDA-approved drug library. Multivariate plot showing PC1, PC2, and PC3. ChemMineTools calculate the Open Babel descriptors as structural fingerprints^28^. Contributions of each parameter and the loading plots for the first three principal components are shown in **Supplementary Tables S1**-**S3**. **b**, Chemical structures of Hydroxychloroquine (HCQ). **c**, Dot plot showing the pairwise 2D Tanimoto coefficient (Tc) between Lev and known RNA binders. RNA binders with Tc≥ 0.7 are colored red; other RNA binders are colored grey. **d**, Chemical structure of acylating probe (HCQ-AI) to identify transcriptome-interaction of HCQ. The drug part is colored in dark green, linker in black, “click” handle in red, and RNA-reactive warhead in green. **e**, UCSC tracks showing the RBRP scores (y-axis) in live HEK293 cells along the human chromosomes (*top*) and −Log_2_*P*-value of Welch *t*-test (*bottom*) of HCQ-AI probe. **f**-**g**, Chemical structures of Dasatinib (Das) (**f**) and acylating probe (Das-AI) (**g**). **h**, UCSC tracks showing the RBRP scores (y-axis) in live HEK293 cells along the human chromosomes (*top*) and −Log_2_*P*-value of Welch *t*-test (*bottom*) of Das-AI probe. **i**, Lev, HCQ, and Das contain different nitrogen heterocycles including fluoroquinolone, aminoquinoline, pyrimidine, and thiazole, which may affect RNA selectivity. **j**, Heatmap analysis showing that the three drugs have distinct structural features related to Van der Waals surface area (Set 1), sp3 character (Set 2), H-bond acceptor (Set 3), aromaticity and nitrogen rings (Set 4), and topological charge (Set 5), which are predictive of RNA-ligand interactions^29^.

We first characterized the *in vivo* transcriptome-interaction of hydroxychloroquine (HCQ), which was originally approved for the treatment of malaria and recently studied for treatment of COVID-19 infections, and causes retinopathy and cardiomyopathy of unknown origins^33,34^ (**Fig. 3b**). HCQ structurally resembles at least eight known RNA binders (Tc ≥ 0.7) and contains fused aromatic rings and positive charge under the physiological conditions that may add affinity for folded RNAs (**Fig. 3c**). After confirming that treatment with excess HCQ did not significantly alter the abundance of most transcripts (**Extended Data Fig. 2c**), we conducted RBRP experiments with an acylating analog of HCQ (HCQ-AI) in HEK293 at clinically relevant doses (**Fig. 3d**). The comparative RBRP workflow enabled us to identify probe-promoted, HCQ-competable 2’-OH acylation as imprints of selective RNA-ligand interactions. Approximately 4 selective targets were identified in ~16,000 RNAs of HEK293 cells (**Fig. 3e**), most of which were independently confirmed by RT-qPCR (**Extended Data Fig. 7b**).

We also profiled the transcriptome interactions of Dasatinib (Das), a multi-targeted kinase inhibitor known to cause neutropenia and myelosuppression, heart failure, and serious pulmonary arterial hypertension^35^ (**Fig. 3f**). The cationic Das contains heterocyclic flat structures, and amide linkages, reminiscent of groove-binding ligands for nucleic acids^36,37^. Base stacking and partial intercalation by Das are also likely feasible. RBRP with an acylating analog of Das (Das-AI) revealed three distinct off-target interactions in cells (**Fig. 3g-h**). We identified and validated several RNA loci that were targeted by Das in HEK293 cells (**Extended Data Fig. 7c**). This is consistent with a previous microarray-based drug screening that showed RNA interaction of FDA-approved kinase inhibitors^7^.

Profiling of these additional compounds via RBRP revealed distinct off-target interactions of drugs with the human transcriptome in cells, consistent with three drugs that differ greatly in their overall structures. The three drugs examined here contain different nitrogen heterocycles including fluoroquinolone, aminoquinoline, pyrimidine, and thiazole (**Fig. 3i**) that may contribute to their differential RNA selectivity^29,36^. The drugs also greatly differ in other features including Van der Waals surface area, sp3 character, aromaticity and nitrogen rings, and topological charge, which also likely influence their RNA recognition (**Fig. 3j**)^29,38^.

### Alteration of RNA structure and RBP binding associates with drug engagement

Despite the existence of pervasive transcriptome interactions, whether these off-target drug binding events may cause unintended biological effects remained unclear. To infer potential effects, we sought to identify the binding loci that engage with RBPs, which extensively associate with RNAs to exert biological functions^39^ (**Fig. 4a**). Taking the off-target small nucleolar RNA (SNORD110) as an example, SNORD110 binds to Lev and contains an oligo(A) tail at its 3’ end that allowed detection by RBRP^30^ (**Fig. 4b** and **Supplementary Information Fig. 1**). Cross-linking immunoprecipitation followed by sequencing (CLIP-seq) experiments^40^ showed that Lev engages with an RBP-binding sequence within SNORD110 (**Fig. 4b**). At this locus, Gene Ontology (GO) analyses demonstrated that the underlying RBPs possess numerous RNA-processing functions (*e.g*., rRNA modification) (**Fig. 4c**), suggesting potential perturbation of these functions by Lev. Two RBPs, nucleolar protein 58 (Nop58) and fibrillarin (FBL), form the small nucleolar ribonucleoprotein complex (SnoRNP) with SNORD110, which mediates site-specific 2’-OH methylation of nascent pre-rRNA at 18S rRNA:U1288 and impacts the translational functions of the mature ribosome (**Fig. 4d**)^41,42^. qPCR and deep sequencing validated the selective SNORD110 interaction of Lev (**Extended Data Fig. 4** and **Supplementary Information Fig. 2**). Titration of unmodified Lev in HEK293 cells revealed dose-dependent enhancement of 2’-OH methylation at 18S rRNA:U1288 at clinically relevant doses, with an EC50 of 10 ± 2.1 μM (**Fig. 4e**). Thus, comparative analysis of RBRP (drug binding) and existing iCLIP (RBP binding) data may allow us to generate testable hypotheses to infer possible biological outcomes of RNA off-target binding.

**Fig. 4.**
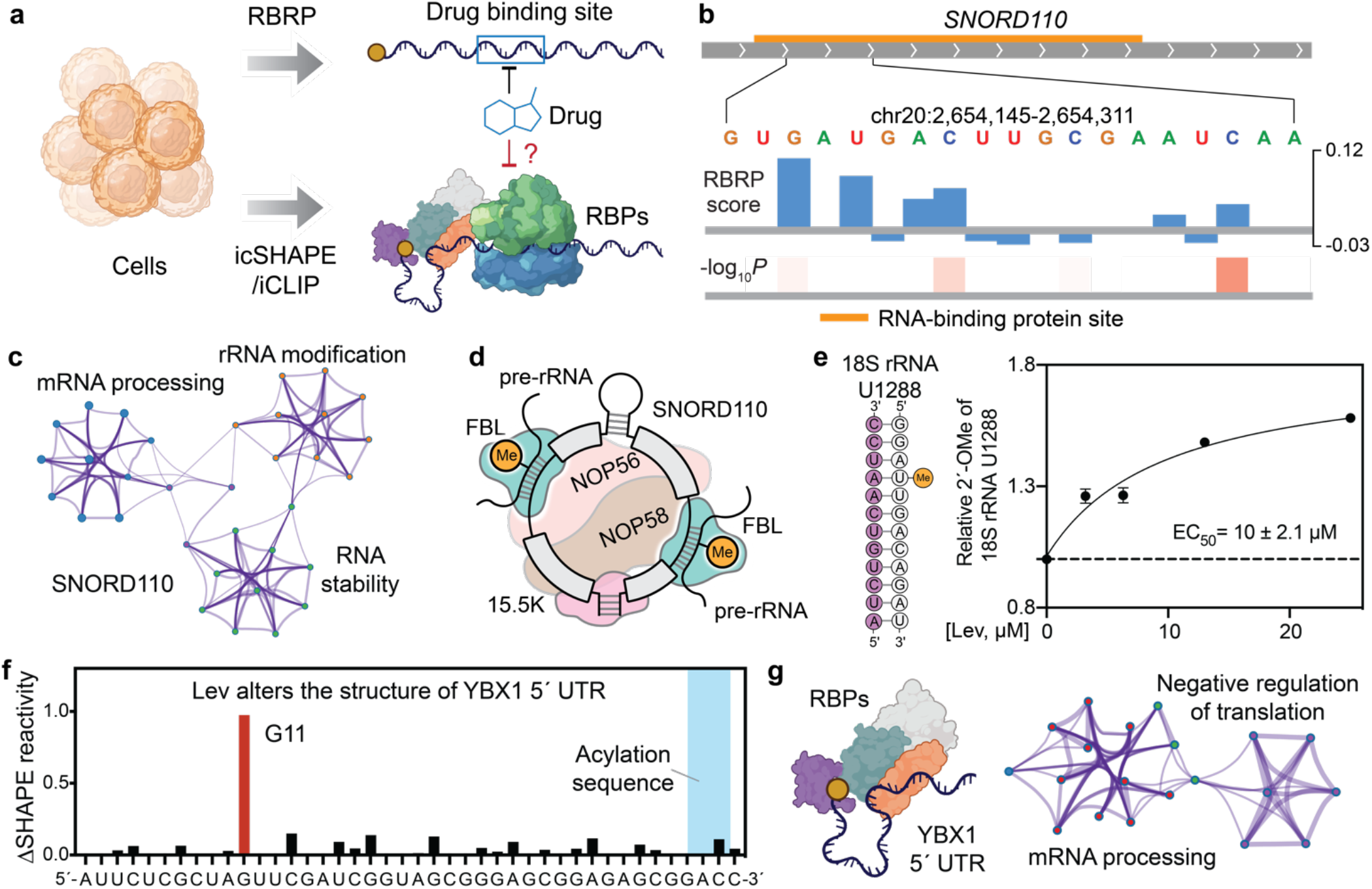
Altered RNA structure and RBP binding can associate with drug engagement. **a**, Meta-analysis reveals association among RBRP reactivity, RNA structural accessibility, and RBP binding to infer biological outcomes of off-target RNA interactions. **b**, UCSC tracks showing RBRP score at an RNA locus in SNORD110-expressing sequence (*top*) and −Log_2_*P*-value of Welch *t*-test (*bottom*), suggesting binding of Lev. RBP-binding sequence is colored orange. **c**, Metascape enrichment network showing the top non-redundant enrichment clusters of SNORD110-binding proteins proximal to the Lev-AI binding site. **d**, Small nucleolar ribonucleoprotein (snoRNP) complex guides 2’-OH methylation of the nascent pre-rRNA. The conserved sequences of SNORD110 are highlighted with box. Fibrillarin (FBL), NOP56, NOP58, and 15.5K are small nucleolar proteins. Me indicates the ribosomal 2’-OH to be methylated. **e**, Lev dose-dependently enhances 2’-O-methylation of 18S rRNA U1288, with an EC50 = 10±2.1 μM. Data shown are mean ± s.e.m., *n*=3 independent biological replicates. **f**, Acylation probing (SHAPE) showing Lev-enhanced structural accessibility of 2’-OH at G11 in YBX1 5’ UTR, a locus proximal to acylation sequence by Lev-AI. Values shown are the differential SHAPE reactivities at probe binding sites in the presence over the absence of Lev, determined by DNA fragment analysis. **g**, Illustration of RBPs binding to YBX1 5’ UTR (left panel) and Metascape enrichment network showing the top non-redundant enrichment clusters of YBX1 5’ UTR-binding proteins proximal to the Lev-AI binding site.

RNA structure also plays a central role in many cellular processes, including RNA transcription, translation, and degradation^43–45^, and is subject to extensive regulation *in vivo* by RBP interactions that can modulate intermolecular accessibility to structural probing reagents. Thus, we also searched for functional binding events that alter folded structure at RNA loci. SHAPE analysis of YBX1 5’ UTR in cell lysates demonstrated that Lev enhances the structural accessibility of G11 in YBX1 5’ UTR at an RBP-binding sequence (**Fig. 4f**). The underlying RBPs at this YBX1 locus are involved in numerous cellular functions including negative regulation of translation (**Fig. 4g**), leading us to wonder whether Lev may not only alter RNA structure at this locus, but also modulate the translation of YBX1 in cells in an off-target way.

### Levofloxacin targets a G-quadruplex in YBX1 5’ UTR and inhibits translation

To test whether Lev modulates YBX1 translation, we treated HEK293 cells with increasing concentrations of Lev and quantified the cellular expression level of YBX1 with Western-blot analysis. We observed a ~30% reduction in YBX1 expression level in cells treated with Lev at clinically relevant doses (**Fig. 5a**).

**Fig. 5.**
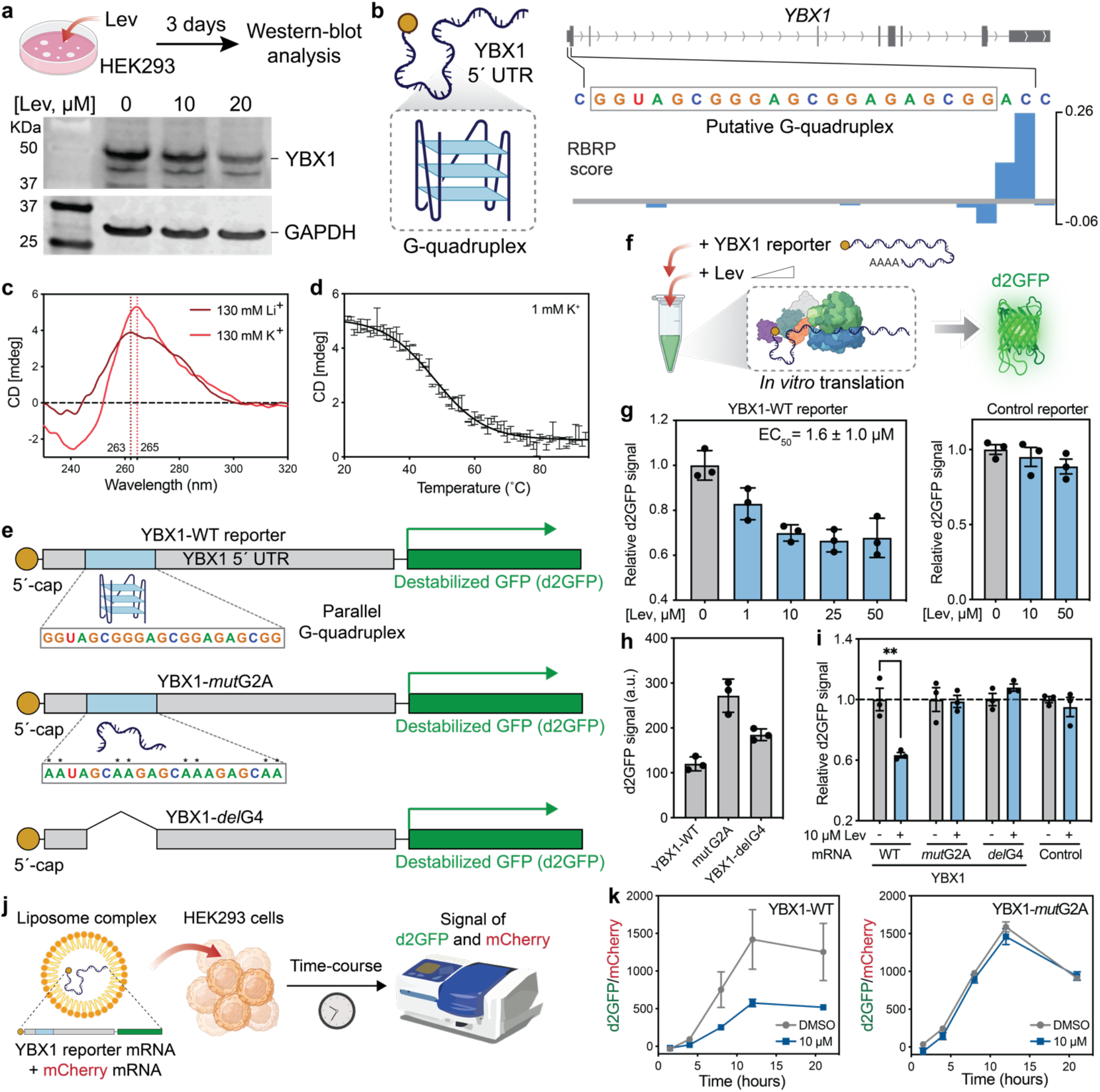
Levofloxacin targets a G-quadruplex in YBX1 5’ UTR and inhibits translation. **a**, Western-blot analysis showing dosedependent suppression of YBX1 expression level in HEK293 cells at clinically relevant doses. **b**, Schematic representation of a putative RNA G-quadruplex within YBX1 5’ UTR (left panel) and a UCSC track showing Lev-AI acylates 2’-OH groups immediately 3’ to the putative RNA G-quadruplex in YBX1 5’ UTR (right panel). **c**, Circular dichroism (CD) spectrum of YBX1 G-quadruplex at 10 μM strand concentration in 10 mM Tris, pH=7.0, and 130 mM of KCl or LiCl at 25 °C. **d**, Melting CD curve of YBX1 G-quadruplex at 10 μM strand concentration in 10 mM Tris, pH=7.0 and 1mM KCl. **e**, Schematic representation of chimeric d2GFP reporter mRNA constructs. YBX1-WT (top), full-length wild-type YBX1 5’ UTR; YBX1-*mot*G2A (middle), GG to AA mutation within G-quadruplex sequence to disrupt quadruplex formation; *YBX1-delG4* (bottom), G-quadruplex sequence deleted from YBX1 5’ UTR. **f**, Workflow determining translation efficiency with cell-free *in vitro* translation systems. **g**, Lev dose-dependently suppressed translation of YBX1-WT reporter, with an EC_50_= 1.6 ± 1.0 μM (left). Lev did not significantly alter the translation of a control d2GFP reporter with scrambled 5’ UTR (right). **h**, Translation efficiency of three chimeric YBX1 reporter mRNAs, as quantified by d2GFP signal. **i**, Relative translation efficiency of three chimeric YBX1 reporter mRNAs and a control mRNA with scrambled 5’ UTR in the presence or absence of 10 μM Lev. Statistical significance was calculated with two-tailed Student’s *t*-tests: \*\**P*<0.01, \*\*\**P*<0.001, \*\*\*\**P*<0.0001. **g**-**i**, Data shown are mean ± s.e.m., *n*=3 independent biological replicates. **j**, Workflow to determine translation efficiency of YBX1 reporter mRNA in HEK293 cells. Chimeric YBX1 reporters are co-transfected with mCherry mRNA. Fluorescence-based microplate reader monitors the translation of d2GFP and mCherry. **k**, Lev inhibits the translation of chimeric YBX1-WT reporter mRNA in HEK293 cells (left), while showing no effects on translation of YBX1-*mut*G2A at clinically relevant dose (10 μM) (right).

To probe the underlying mechanism of action, we first investigated Lev’s minimal binding sequence in YBX1 5’ UTR. RBRP provides information about proximal drug-binding sites at near single-nucleotide level, allowing the identification of the drug’s local binding target. The 5’ UTR of the Y-box binding protein 1 (YBX1) gene contains a putative G-quadruplex (G4) motif (5’- <u>GG</u>TAGC<u>GGG</u>AGC<u>GG</u>AGAGC<u>GG</u>-3’), which is located 18 nucleotides downstream of the 5’cap and 109 nucleotides upstream of the start codon (**Fig. 5b**). Lev-AI probe acylates the YBX1 5’ UTR at a dinucleotide sequence that is immediately adjacent 3’ to the putative G4 sequence. Notably, analogs of Levofloxacin, CX-5461 (Pidnarulex), have been implicated as G4 binders^46^. We have also shown tight engagement of Levofloxacin with YBX1 5’ UTR in cells at clinically relevant doses (**Fig. 2i**). Thus, we speculated that Lev might target the putative G4 sequence within YBX1 5’ UTR.

To confirm the existence of a stable quadruplex structure within YBX1 5’ UTR, we biophysically characterized a synthetic RNA corresponding to the isolated sequence from YBX1 5’ UTR with circular dichroism (CD) spectroscopy (**Fig. 5c-d**), confirming the formation of a parallel G4 structure within the isolated RNA (**Fig. 5c-d**)^47^.

To isolate YBX1 5’ UTR as the drug binding site of Lev, we biochemically quantified the impacts of Lev on the translation of genes encoded downstream of YBX1 5’ UTR with a cell-free translation system coupled to a reporter gene assay (**Fig. 5e-f**). Strong dose-dependent inhibition by Lev of d2GFP translation was seen, with an EC_50_= 1.6 ± 1.0 μM, which is consistent with binding affinity measured in cells (**Fig. 5g**), whereas no dose-dependent effect was seen with a scrambled UTR unable to form G4 structure. The results are consistent with previous observations that small molecules that target G-quadruplex in 5’ UTRs can modulate the translation of mRNA^48,49^.

Mutations can disrupt favorable target interactions of drugs, and can be applied to identify mechanism of drug actions^50^. Thus we engineered two mutants to disrupt G4 formation (**Fig. 5e**)^51^. G4-disrupting mutations (GG to AA) resulted in an increase in translation efficiency and loss of Lev response relative to the native sequence (WT) (**Fig. 5i**). These data also indicate that the natural YBX1 RNA G-quadruplex in its 5’ UTR has an inhibitory effect on translation. In addition, deletion of the G-quadruplex motif (YBX1-*del*G4) resulted in nearly complete resistance to Lev-mediated suppression of mRNA translation, which further documents this G4 as a Lev off-target binding site.

Finally, we showed that Lev targets the G4 quadruplex at YBX1 5’ UTR in living cells. YBX1 UTR reporter mRNAs were lipofected into HEK293 cells with or without Lev at a clinically relevant dose (10 μM). Translation kinetics of YBX1 reporter was monitor by green fluorescence signal in living HEK293 cells (**Fig. 5j**). We found that treatment with Lev markedly reduced the translation efficiency of G-quadruplex-containing YBX1-WT reporter (**Fig. 5k**, left panel). In contrast, Lev did not affect the cellular translation efficiency of YBX1-*mut*G2A, further supporting that Lev targets YBX1 G4 in cells (**Fig. 5k**, right panel). Thus, Lev-dependent reduction of YBX1 protein expression is at least partially caused by Lev targeting the translation-suppressive G-quadruplex within the YBX1 5’ UTR in living cells.

Taken together, our data strongly suggest that FDA-approved protein-targeted drugs can cause unintended biological outcomes by engaging with off-target cellular RNAs. Stringent transcriptome-wide validation of drug binding sites may provide more comprehensive insights for the potential mechanisms of dose-limiting drug side effects.

## Conclusions

It is generally understood that small-molecule drugs can exhibit toxicity by off-target interactions within the human proteome^52^. Our data suggest that, in addition to proteome interactions, off-target engagement with specific RNAs in the human transcriptome may also be a mechanism by which small molecules cause unintended biological consequences. Each ostensibly protein-targeted drug tested in this study was found to target numerous transcripts in cells. This is consistent with previous observations that known drugs including FDA-approved drugs can bind RNAs (e.g., Mitoxantrone targets pre-miR-21), which was discovered with an *in vitro* microarray-based screening and later validated in cells^7^. Considering the high chemical similarity of many existing protein-targeted drugs with known RNA-binding molecules (Tanimoto coefficient ≥0.7), this phenomenon is likely common among many FDA-approved drugs and those in clinical trials and preclinical studies. Given that ~90% of drugs fail to receive FDA approval and hundreds of previously approved drugs have been withdrawn^52,53^, RNA off targets may likely contribute to failure rates. Thus, stringent characterization of transcriptome interactions for smallmolecule drugs may be helpful in shedding light on these problems.

This study highlights the power of reactivity-based RNA profiling (RBRP) for target validation of small-molecule drugs, which requires the use of cell-permeable RNA acylation probes (reacting at 2’-OH groups) and a comparative workflow to scout off-target interactions of drugs in the human transcriptome. Direct comparison with an orthogonal diazirine-based RNA-ligand profiling method validated and highlighted the power of RBRP to identify transcriptome interactions of drugs, complementing existing diazirine-based methodology. RBRP bridges a gap in RNA-sequencing technologies that currently lack the ability to impartially identify drug binding sites at all four nucleotides (**Extended Data Fig. 5f**), without potential interference from extended incubation or UV light. The ability to quantify off-target interactions at each nucleotide in more than ~16,000 poly(A)+ RNAs enables unprecedented identification of altered RNA structural accessibility and RBP underlying drug engagement and allows the users to infer the cellular consequences of off-target RNA binding. Notably, we quantified and observed minimal correlation between the RBRP and icSHAPE reactivities transcriptome-wide (**Supplementary Information Fig. 3a-b**), confirming that RBRP signals are drug-promoted rather than simply reflecting structure-dependent stochastic acylation. This observation is also confirmed by the fact that RBRP signals were minimally correlated among three different drugs (**Supplementary Information Fig. 3c**).

Our results indicate that many existing drugs may exert unintended biological effects through off-target interactions with the human transcriptome *in vivo*. For example, we find that off-target transcriptome interaction may unexpectedly affect 2’-OH methylation levels of ribosomal RNAs. Our data also show that the antibiotic Levofloxacin targets a G-quadruplex in YBX1 5’ UTR and causes collateral inhibition of YBX1 translation in cells at clinically relevant doses. Not surprisingly, our findings provide evidence that transcriptome interaction of drugs depends greatly on their chemical structures^36,38,54^. Indeed, structural features such as positive charge and flat aromatic structures for groove binding and intercalation can add affinity for structured RNAs *in vitro* with low to moderate selectivity^36,37,55^. Consistently, our work shows that intercalator-like Lev extensively interacts with highly structured RNAs in cells, whereas Das is more structurally complex than a simple intercalator and has fewer RNA off targets. Although our work provides strong evidence that transcriptome-interaction can alter cellular outcomes in specific cases, it is likely that some off-target interactions within the human transcriptome will not cause unintended cellular consequences, if the binding does not alter RNA conformation or RPB interactions. In addition, it remains possible that some drug targets will prove to be cell type-dependent, which is not studied here, and will require further investigation.

More broadly, we suggest that rigorous characterization of transcriptome interaction should be essential not only for existing drugs, but also future drugs. Animal toxicity testing fails to predict side-effects and toxicity in ~50% of drugs between Phase I human trials and early post-market withdrawals^56^, calling for comprehensive target validation during preclinical studies. Others in the field have also postulated that off-target RNA interactions may likely fuel the failure of drug safety assessment^8^. Given that dose-limiting toxicity is a chief driver for clinical trial failures and drug withdrawal, stringent validation of drug targets in both the human proteome and transcriptome should not only reduce costs, but also allow the redistribution of resources towards lead compounds that are more likely to pass clinical trials. We envision that UV irradiation-free, minimally nucleobase-biased RBRP experiments can facilitate our understanding of RNA-mediated drug effects, indicate potential toxicity biomarkers, and characterize toxicity mechanisms.

## Supporting information

Supp file S1_Tanimoto coefficients of all drug-RNA binder pairs

Supp file S2_FDA approved RBinders

Supp file S3_RBRP results

Supp file S4_primers

## Methods

### Materials

DNA and RNA were purchased from IDT or Stanford PAN facility unless otherwise stated (see **Supplementary Table S4** for details). All chemicals purchased from commercial suppliers were used without further purification (see **Supplementary Table S5** for details). All enzymes, kits, bioreagents, and software were obtained from sources listed in Supplementary Table S5. Human cell line HEK293 (CRL-1573) was purchased from American Type Culture Collection (ATCC) and maintained in DMEM supplementary with 10% fetal bovine serum at 37 °C in a humidified incubator containing 5% CO_2_.

### Synthesis of acylimidazole probes

HCQ-AI, Lev-AI, and Das-AI were synthesized as described in the Supplementary Information and activated immediately before use: In general, the carboxylic acid precursors (1.0 equiv.) were dissolved in dry DMSO. To it was added a suspension containing 1.4 equivalent of 1,1’-carbonyldiimidazole (CDI) at room temperature. The solution was stirred at room temperature overnight under argon. After the reaction, the final solution can be used as a 50 mM acylimidazole stock without further purification. See Supplementary Information for details.

### Curation of small-molecule drugs database

Database curation of FDA-approved drugs. Lists of FDA-approved small-molecule drugs were acquired from a previous study^5^. Briefly, FDA-approved drugs were downloaded from DrugBank (v5.1.6, released on 04/22/2020) on 06/15/2020. The drug list was further filtered to remove molecules according to the reported criteria^5^. Molecules with molecular weights larger than 140 Da and smaller than 706 Da were used for statistical and principal components analysis.

Database curation of drugs in preclinical studies, clinical trials, and drugs withdrawn from the market. The small molecules were acquired from Drug Repurposing Hub^24^ on 06/05/2022. The molecules were annotated based on their disease areas and stages in clinical studies. Molecules with molecular weights that are larger than 140 Da and smaller than 706 Da were used for statistical analysis.

Database curation of RNA-binding small-molecule ligands (RNA binders). The small-molecule (SM) RNA binders were downloaded from R-bind 2.0^5^ on 05/24/2022. Molecules (131) with known PubChem Compound ID number (CID) were further used for Structural similarity analysis using the Tanimoto coefficient (Tc).

### Quantification of structural similarity with Tanimoto coefficient (Tc)

Substructure key-based 2D Tanimoto coefficients were calculated using PubChem Score Matrix Service. Lists of CIDs for 131 known RNA binders and molecules of interest were used as input to calculate the pairwise Tanimoto coefficients with each of 131 known RNA binders. Tanimoto coefficients of all drug-RNA binder pairs are listed as the matrix in **Supplementary file S1**.

### Chemoinformatic calculation of chemical fingerprints

Fourteen Open Babel descriptors of the selected FDA-approved drugs were calculated using ChemMine Tools^28^. See **Supplementary file S2** for the detailed list.

### Principal component analysis of chemical fingerprints of small-molecule drugs

Fourteen Open Babel descriptors were used as chemical fingerprints in the standardized principal component analysis (PCA). PCA was performed using Graphpad Prism 9. The principal components (PC1 and PC2) were selected based on parallel analysis, with eigenvalues greater than 95% percentile of Monte Carlo simulations (number of simulations: 1,000) on “random” data of equal dimension to the input data. See **Supplementary Tables S1-2** for details.

### Treatment of cells with acylimidazole probes, isolation of total cellular RNA, and enrichment of poly(A)+ transcripts

~2×10^6^ of cells were pretreated with DMSO or excess competing drugs for 30 minutes at 37 °C. The cells were then immediately scraped into a 15-mL falcon tube and resuspended in 1 mL of 1xPBS, pH=7.4 at 37 °C containing 50 μM of acylimidazole probe, in the absence or presence of excess competing drugs. After incubation at 37 °C for 30 minutes, the reaction was stopped by centrifugation. See Supplementary Information for details.

To isolate total cellular RNA, cells were lysed with 6 mL of Trizol LS reagent according to the Supplementary Information. RNAs were then purified with a Zymo Quick-RNA Midiprep kit according to the manufacturer’s protocol.

To enrich poly(A)+ transcripts, a 250-μg aliquot of total cellular RNA was diluted to 600 μg/mL with RNA storage buffer and then purified with Poly(A)Purist MAG kit according to the manufacturer’s protocol. The captured RNAs were further purified with RNeasy Mini columns according to the manufacturer’s protocol, and stored as 500-ng aliquots at −80 °C.

### Preparation of sequencing libraries

Sequencing libraries were prepared according to the following protocol based on a previous report^23,27^ with modifications: First, the probe-labeled RNAs were modified with biotin by “click” reaction. 500 ng of poly(A)+ RNAs were incubated with 2 μL of 1.85 mM DIBO-Biotin and 1 μL of RiboLock at 37 °C for 2 hours. After the reaction, modified RNAs were purified with a Zymo RNA Clean & Concentrator column-5 according to the manufacturer’s protocol.

(RNA fragmentation, 3’-end repair, and 3’-end ligation) RNAs were fragmented using RNA fragmentation reagent, with rendered RNA fragments with a medium length ~100 nt. After purification with a Zymo RNA Clean & Concentrator column-5, the 3’-end of RNA fragments were repaired by FastAP and T4 PNK for 1 hour at 37 °C according to the previously reported protocol^23,27^. Next, 3’-ligation was performed to install an 3’-blocked RNA ligand to the 3’-end of RNA fragments. Briefly, RNAs were incubated with T4 RNA ligase 1 and preadenylylated RNA links (Probe-treated sample: 3’-ddC blocked RNA linker /5rApp/AGA TCG GAA GAG CGG TTC AG/3ddC/ was used; For DMSO-treated samples, 3’-biotin blocked RNA linker /5rApp/AGA TCG GAA GAG CGG TTC AG/3Biotin/ was used. After the 3’-end ligation reaction, RNA was purified with a Zymo RNA Clean & Concentrator Column-5 and then resolved on a 6% (w/v) UreaGel denaturing PAGE. RNAs with a length above ~50 nt were extracted and further purified using a 10K Amicon filter as described in Supplementary Information.

(cDNA synthesis, enrichment of Biotin-modified RNAs, and cDNA purification) The following steps were performed according to the previously reported protocol with slight modifications^23,27^: The 3’-ligated RNA was dissolved in 11 μL of RNase-free water and 1 μL of 1 μM barcoded RT primer (/5phos/DDD NN<u>A ACC</u> NNN NAG ATC GGA AGA GCG TCG TGG A/iSp18/GGATCC/iSp18/TACTGAACCGC. D = A/G/T and N = A/T/G/C. The underlined four nucleotides are barcodes)(see Supplementary Table S4 for assignments of RT primers). The reaction was heated at 70 °C for 5 minutes and then cooled to 25 °C by 20°C/60 seconds. 8 μL of RT reaction mixture (4 μL of 5x First strand buffer, 0.75 μL of RiboLock, 1 μL of 100 mM DTT, 1 μL of 10 mM dNTPs, and 1.25 μL of SuperScript III) was then added to the above reaction. The resulting solution was heated at 25 °C for 3 mins, 42 °C for 5 mins, and 52 °C for 30 mins. Next, the cDNA-RNA heteroduplexes were enriched with MyOne C1 magnetic streptavidin beads and the cDNA was purified with a 6% (w/v) denaturing PAGE gel as described in the Supplementary Information.

(cDNA circularization and library amplification) The purified cDNA was circularized with CircLigase II for 3 hours at 60 °C and purified with Zymo DNA Clean & Concentrator Column-5 according to the manufacturers’ protocols. To amplify the library, 20 μL of 2x Phusion HF PCR master mix, 0.4 μL of SYBR Green I (25x), 1.0 μL of 10 μM P5-universal PCR primer (5’-AAT GAT ACG GCG ACC ACC GAG ATC TAC ACT CTT TCC CTA CAC GAC GCT CTT CCG ATC T-3’ and 10 μM P3-universal PCR primer (5’-CAA GCA GAA GAC GGC ATA CGA GAT CGG TCT CGG CAT TCC TGC TGA ACC GCT CTT CCG ATC T-3’). PCR amplification was monitored in real-time to avoid over-amplification. The PCR reaction was first denatured at 98 °C for 45 seconds, followed by cycles of denaturation (98 °C, 15 seconds), Anneal (65 °C, 20 seconds), and Extension (72 °C, 1 minute). The PCR products were purified with a 3% low melting point agarose gel and extracted with the QIAGEN MiniElute Gel extraction kit. The purified DNA was analyzed by BioAnalyzer for quality control and quantified with Qubit.

### Next-generation sequencing

Ten sub-libraries were combined as a sequence library. The sequence library was sequenced on an Illumina-based HiSeq X-ten with a 23% spike-in PhiX sequencing control. Paired-end 150 bp sequencing was performed. The raw data were transformed into single-end reads for the following bioinformatics analysis with the icSHAPE pipeline. Note: the DMSO-treated sequence sub-libraries are amplified for fewer cycle numbers and thus have higher complexity than the probe-treated sequence sub-libraries. Thus, more sequencing depth should be allocated to the probe-treated sequence sub-libraries.

### Bioinformatics and RBRP data visualization

We use a bioinformatics pipeline modified based on icSHAPE^23,27^ to calculate the RBRP score at each nucleotide transcriptome wide. See Supplementary Information for detailed protocols and command lines. Bigwig files can be visualized using alignment visualization tools such as Integrative Genomics Viewer (IGV) or UCSC Genome Browser.

### Identification of off-target RNA binding sites

RBRP scores at each site were calculated as the surplus yield of 2’-OH acylation for experiments performed in the absence over the presence of excess competing drug. We defined potential RNA-binding sites as nucleotides with RBRP scores ≥0.12 and read depths≥ 200 (see **Supplementary file S3** for all competed acylation sites).

### Target validation with RT-qPCR

HEK293 cells were treated with the acylation probe in the absence or presence of excess competing drugs. Total cellular RNA was extracted with the Trizol LS reagent and purified according to the protocol described above. 1 μg (2.5 μL, 400 ng/μL) of total RNA was mixed with 1 μL of 10 mM dNTPs, 1 μL of 2 μM RT primer, and 8.5 μL of water. The reaction mixture was heated at 65 °C for 5 minutes and then cooled on ice for > 1 minute. Next, 4 μL of 5x first-strand buffer, 1 μL of 0.1 M DTT, 1 μL of RiboLock, and 1 μL of SuperScript III was added to the reaction. The reaction solution was then incubated at 25 °C for 5 minutes, 55 °C for 30 minutes. After inactivation at 70 °C for 15 minutes, 0.5 μL of RNase H was added and incubated at 37 °C for 20 minutes.

qPCR. 1 μL of the above cDNA product was added to a solution containing 10 μL of Luna University qPCR Master Mix, 0.5 μL of 10 μM forward primer, 0.5 μL of 10 μM reverse primer, and add water to 20 μL. The resulting mixture was amplified using a StepOnePlus real-time qPCR system, starting with an initial denaturation step at 95 °C for 60 seconds, followed by 40 cycles of denaturation (95 °C, 15 seconds), anneal and extension (60 °C, 30 seconds). ΔCt value was calculated by subtracting Ct(U) by Ct(D): ΔCt=Ct(U)-Ct(D). qPCR amplified regions upstream (U) and downstream (D) of the acylation sites. An increased ΔCt value indicates acylation at the tested sites.

### Circular dichroism (CD) experiments

CD experiments were performed using a Jasco J-810 spectropolarimeter. The RNA solution in 10 mM Tris (pH=7.0) and desired concentration of KCl was heated at 95 °C for 3 minutes, then cooled to 25 °C over 1 hour. 250 μL of folded RNA was placed in a quartz cuvette with an optical path length of 1 mm. Three CD scans, over the wavelength range of 220 to 320 nm, were performed at 50 nm/minute with a 1-nm bandwidth. Average CD curve was used for further analysis. In addition, CD spectrum of the buffer was determined and subtracted from the spectrum obtained for the RNA-containing solution. Melting curve measurement: Folded RNA was heated from 20 °C to 95 °C at a 1 °C/minute ramping rate. CD ellipticity at 265 nm was measured with D.I.T.= 2 seconds.

### Cloning

We PCR-amplified the open reading frame and 3’ UTR sequence of a d2GFP-encoding plasmid according to a previous report^57^. Next, 5 ng of the above PCR product (8.4 fmol) is further PCR-amplified for ten cycles using Q5 High-Fidelity 2x Master Mix (NEB) with sequence-specific forward primers to install the T7 promoter sequence, variants of YBX1 5’ UTR, and poly(T)_120_ tail (**Supplementary file S4**). Finally, 1 μL of the second PCR product is further amplified with two universal primers (T7-PCR-fwd and Tail-PCR-rev) using Q5 High-Fidelity 2x Master Mix. This PCR product is purified with 1% agarose gel.

### Synthesis of reporter mRNA

The dsDNA template of YBX1-WT, YBX1-mutG2A, and YBX1-delG4 were transcribed using HiScribe T7 Quick High Yield RNA Synthesis Kit (NEB) and 5’-capped with Vaccinia Capping System according to manufacturer’s protocol.

### *In vitro* translation

600 ng of mRNA (YBX1-WT, YBX1-mutG2A, YBX1-delG4) was diluted into 1 x rG4 folding buffer (10 mM Tris, pH7.0, 0.1 mM KCl) to 17.82 μL. mRNAs were folded by heating at 90°C for 3 min and then cooling down to 25 °C at 0.1°C/second. Next, 0.18 μL of 100x Levofloxacin solution at desired concentration or DMSO was added and incubate at 25°C for 30 minutes. 6 μL of the above mRNA solution was added to a cell-free translation system containing 17.5 μL of rabbit reticulocyte lysate, 0.5 μL of RNasin^®^ Ribonuclease Inhibitor, 1 μL of 1 mM complete amino acid mixtures, and 0.19 μL corresponding 100x Levofloxacin solution or DMSO. The resulting mixture was incubated at 30°C for 90 min. After incubation, the green fluorescence signal was determined by a BioTek microplate reader (excitation/emission= 485 nm/520 nm).

### SHAPE experiment

1 μg of YBX1-WT mRNA was added to 1 x rG4 folding buffer (10 mM Tris, pH 7.0, 0.1 mM KCl) to 11.88 μL. Next, 0.12 μL of 100x levofloxacin solution or DMSO was added. After incubation at 25 °C for 30 minutes, 12 μL of the above mRNA solution was added to a cell-free system containing 35 μL of rabbit reticulocyte lysate, 1 μL of RNasin^®^ Ribonuclease Inhibitor, 1 μL of 1 mM complete amino acid mixtures, and 0.38 μL of 100x levofloxacin solution (1 mM to 5 mM) or DMSO. After incubation at 30 °C for 30 minutes, 5.5 μL of 2.0 M NAIN3 reagent was added and incubated at 25 °C for 15 minutes. The reaction mixture was purified by Zymo RNA clean-up and concentrator-5 column following manufacture’s protocol. The purified mRNA was mixed with 0.2 μL of 10 μM FAM-labeled RT-primer, 0.5 μL of 10 mM dNTP, and water to 6.5 μL. The mixture was heated at 75°C for 5 min and chilled on ice for 2 min. Next, reverse transcription was performed by adding 2 μL of 5x first strand buffer, 1 μL of 0.1 M DTT, 0.25 μL of 40 U/μL RiboLock RNase inhibitor, and 0.25 μL 200 U/μL SuperScript III. The reaction was heated at 25 °C for 10 minutes, 50 °C for 50 minutes, and 55 °C for 50 minutes. After reaction, 1 μL of 1 N NaOH was added and incubated at 95 °C for 3 min, followed by the addition of 12 μL of 8M urea loading buffer and heating at 95 °C for additional 3 minutes. The resulting samples were resolved on an 8% denaturing urea PAGE gel and imaged on an Amersham Typhoon gel imager.

### Quantification of the 2’-O-methylation level of human rRNA

To quantify 2’-O-methylation of rRNA, we performed Reverse Transcription at low deoxy-ribonucleoside triphosphate (dNTP) concentrations followed by polymerase chain reaction (PCR) (RTL-P)^58^. HEK293 cells were seeded on a 6-well plate in 2 mL DMEM containing 10% FBS. 20 μL of DMSO or concentrated DMSO solution of drug were added to the cells to reach desired drug concentrations. The cells were incubated at 37 °C in a humidified incubator containing 5% CO_2_ for two days. After two days, cells were washed with 2 mL of DPBS and then lysed with 600 μL of Trizol LS reagent, followed by RNA purification with Quick RNA-miniprep kits and on-column digestion of genomic DNA.

For RTL-P, reverse transcription was performed with 1 μg extracted RNA (5 μL), 1 μL of dNTP at low concentration (10 μM), 1 μL of 2 μM RT primer. The solution was heated at 65 °C for 5 minutes and then incubated on ice for > 1 minute. Next, 4 μL of 5x First-strand buffer, 1 μL of 0.1 M DTT, 1 μL of RiboLock, and 1 μL of SuperScript III were added to the reaction. The reaction mixture was then incubated at 55 °C for 50 minutes, followed by incubation at 70 °C for 15 minutes. 1 μL of the above cDNA product was added to a solution containing 10 μL of Luna University qPCR Master Mix, 0.5 μL of 10 μM forward primer, 0.5 μL of 10 μM reverse primer, and add water to 20 μL. The resulting mixture was amplified using a StepOnePlus real-time qPCR system, starting with an initial denaturation step at 95 °C for 60 seconds, followed by 40 cycles of denaturation (95 °C, 15 seconds), anneal and extension (60 °C, 30 seconds). ΔCt value was calculated by subtracting Ct(U) by Ct(D): ΔCt=Ct(U)-Ct(D). qPCR amplified regions upstream (U) and downstream (D) of the 2’-OMe sites. An increased ΔCt value indicates acylation at the tested sites. ΔCt value was calculated by subtracting Ct(U) by Ct(D): ΔCt=Ct(U)-Ct(D). An increased ΔCt value indicates enhanced of 2’-O-methylation at the targeted sites.

### Statistical analysis

For all statistical tests (unless otherwise noted), a one-tailed or two-tailed Student’s *t*-test was used to compare means between two samples. Significance is denoted as follows: * = *P*< 0.05, ** = *P*< 0.01, *** = *P*< 0.001, **** = *P*< 0.0001. Student’s t-tests were performed in GraphPad Prism9 software. Quantifications shown are mean ± s.e.m. unless otherwise stated.

## Data availability

All sequencing data are available through the Gene Expression Omnibus (GEO). Data supporting the findings of this study are available in the article, Supplementary Information, and Supplementary files S1-S4. Source data and bedgraphs of RBRP scores are also freely available at Figshare at https://doi.org/10.6084/m9.figshare.20326824.

## Code availability

The RBRP scripts used for bioinformatics analysis are freely available at https://github.com/linglanfang/RBRP.

## Acknowledgements

We acknowledge support from U.S. National Institutes of Health (GM130704 and GM145357) for support of this work. We thank Anita Đonlić and Emily G. Swanson for sharing the list of FDA-approved drugs. We thank Theresa McLaughlin at Vincent Coates Foundation Mass Spectrometry Laboratory, Stanford University Mass Spectrometry (RRID:SCR_017801) for acquiring the HRMS data. We thank Steven Lynch for his help with NMR studies. We thank staffs at Stanford PAN facility for supports with oligo synthesis. This work was also supported in part by NIH P30 CA124435 utilizing the Stanford Cancer Institute Proteomics/Mass Spectrometry Shared Resource and NIH High End Instrumentation grant (1 S10 OD028697-01).

---

**Extended Data Fig. 1.**
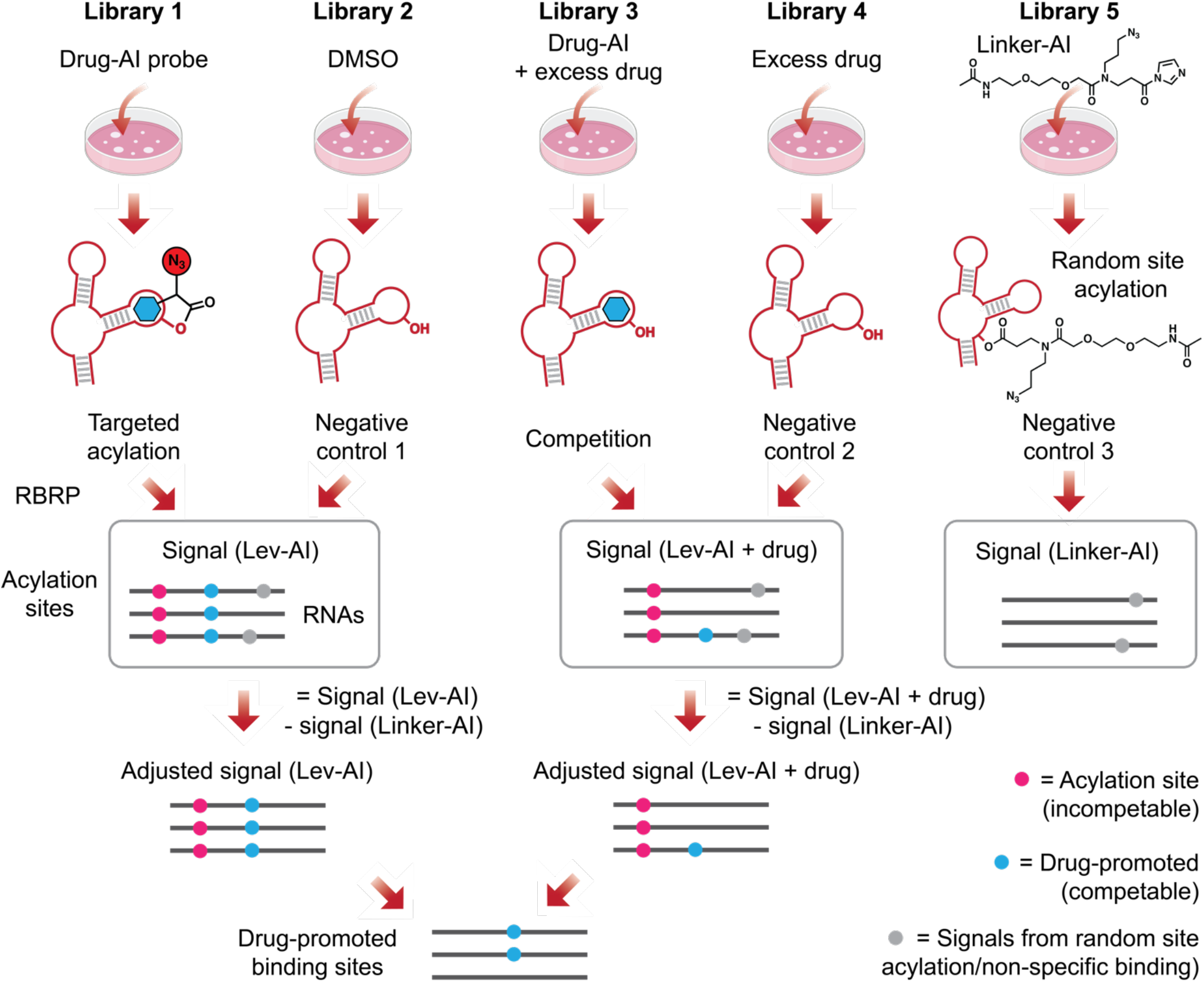
RBRP workflow and experimental setup. **Library 1**: Cells were treated with the drug-conjugated acylating probe. RBRP workflow identifies RBRP signals from drug-promoted acylation, random site acylation, and non-specific binding events during biotin-mediated pulldown. **Library 2**: Cells were treated with DMSO. RBRP workflow identifies RBRP signals from non-specific binding events. **Library 3**: Cells were treated with drug-conjugated acylating probe and the unmodified drug. RBRP workflow identifies RBRP signals from random site acylation, non-specific binding events during biotin-mediated pulldown, and changes in transcript abundance. **Library 4**: Cells were treated with excess unmodified drugs. RBRP workflow identifies RBRP signals from non-specific binding events during biotin-mediated pulldown and changes in transcript abundance. **Library 5**: Cells were treated with Linker-AI. RBRP workflow identifies RBRP signals from random site acylation and non-specific binding events during biotin-mediated pulldown.

**Extended Data Fig. 2.**
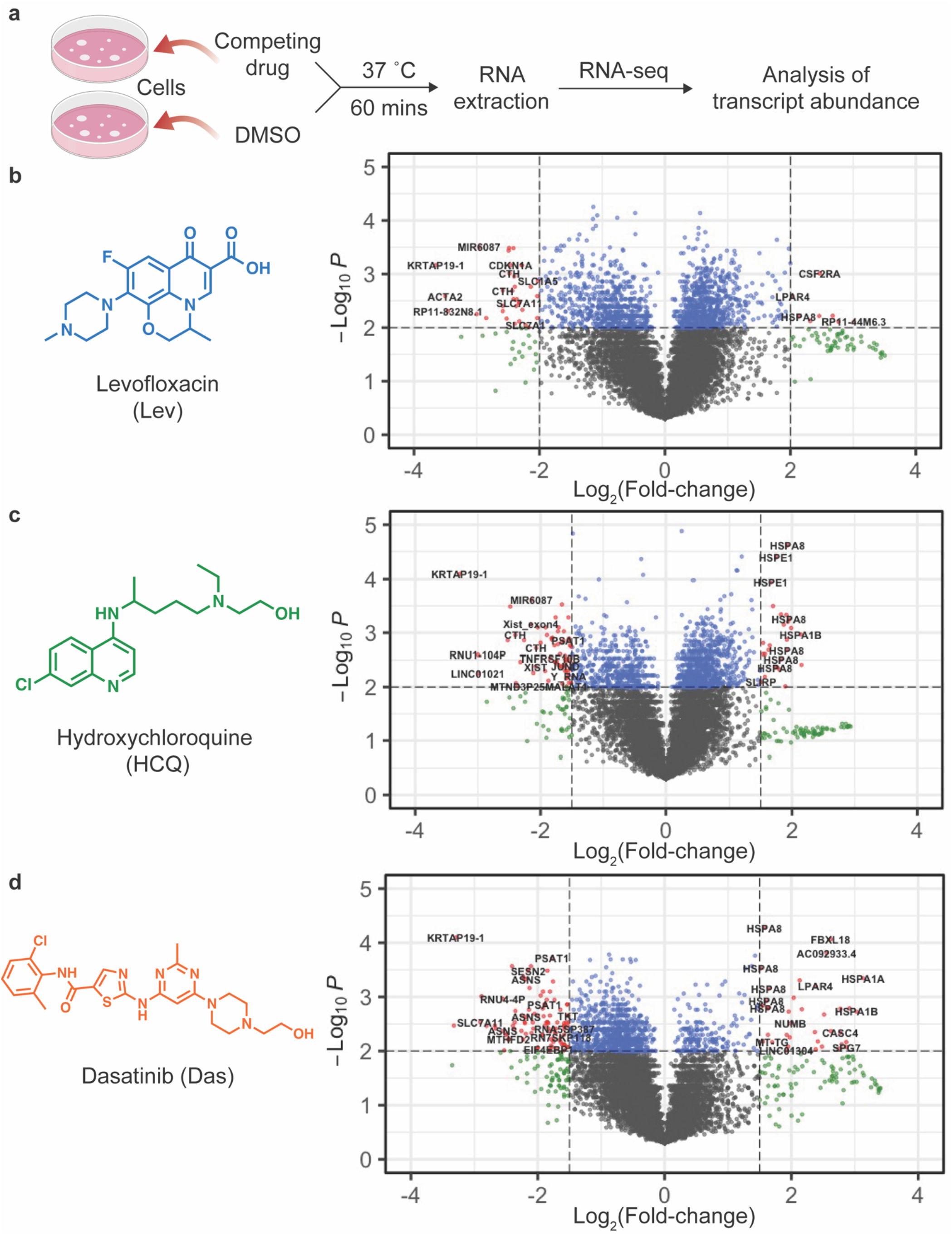
RNA-seq determines how excess unmodified drug influences transcripts abundance in HEK293 cells. **a**, Workflow of RNA-seq with HEK293 cells treated with unmodified drugs or DMSO. **b**-**d**, Volcano plot showing effects of unmodified Lev (**b**), HCQ (**c**), and Das (**d**) on transcript abundance. X-axis: log-transformed ratio of transcript abundance in the presence over the absence of unmodified drug. Y-axis: the negative value of log-transformed *P*-value calculated with DESeq2.

**Extended Data Fig. 3.**
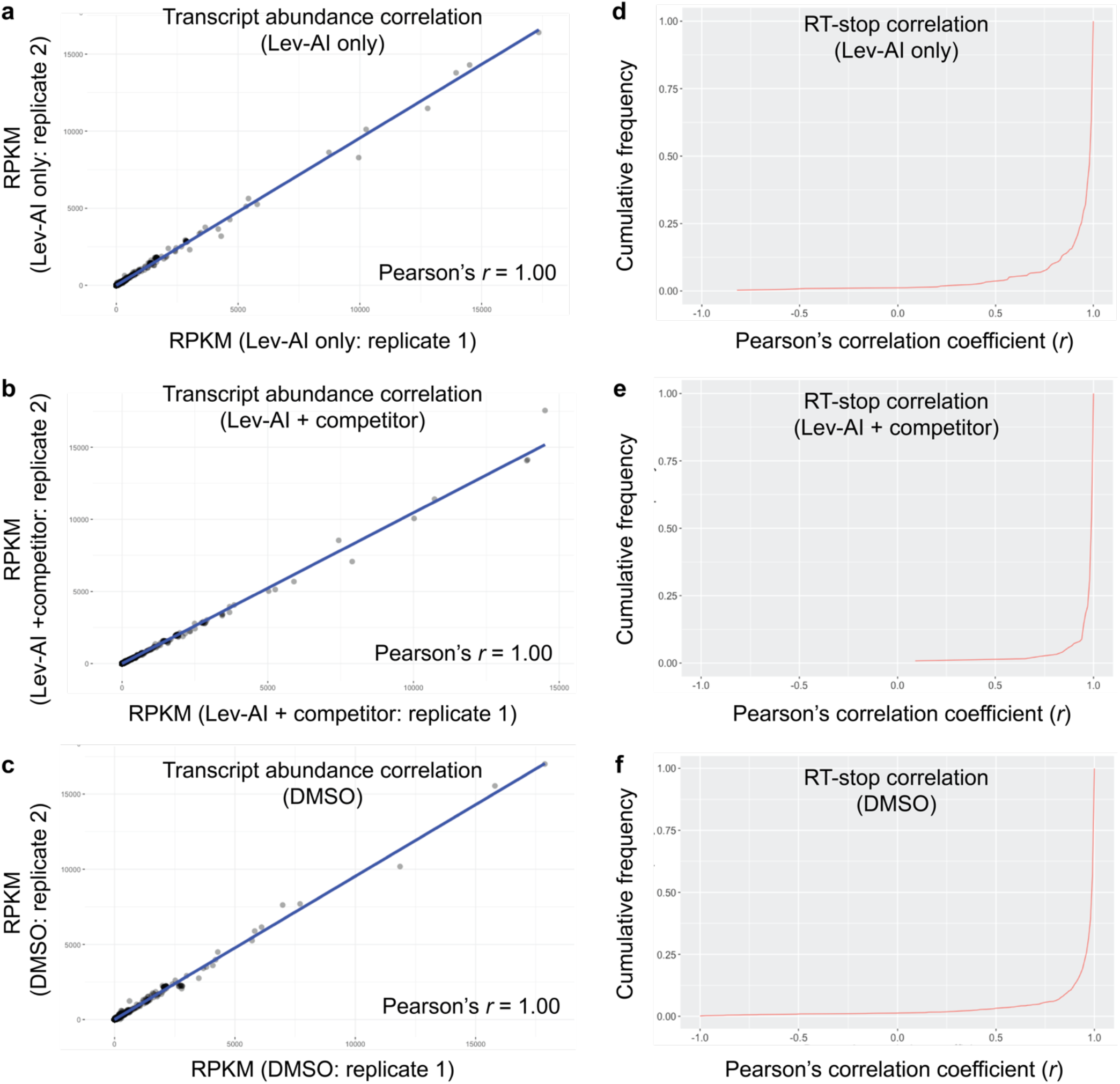
Transcript abundance and RT-stop frequencies are strongly concordant between RBRP sequencing libraries from two biological replicates. **a**-**c**, Scatter plot showing very strong correlation of transcript expression value (RPKM) between two biological replicates (Pearson correlation *r* = 1.00) in HEK293 cells treated with Lev-AI only (**a**), Lev-AI and excess unmodified Lev (**b**), and DMSO (**c**). **d**-**f**, The concordance of RT-stop frequencies is high for most transcripts of read depth higher than the optimized cutoff value (200) in sequencing libraries of Lev-AI only (**d**), Lev-AI and excess unmodified Lev (**e**), and DMSO (**f**).

**Extended Data Fig. 4.**
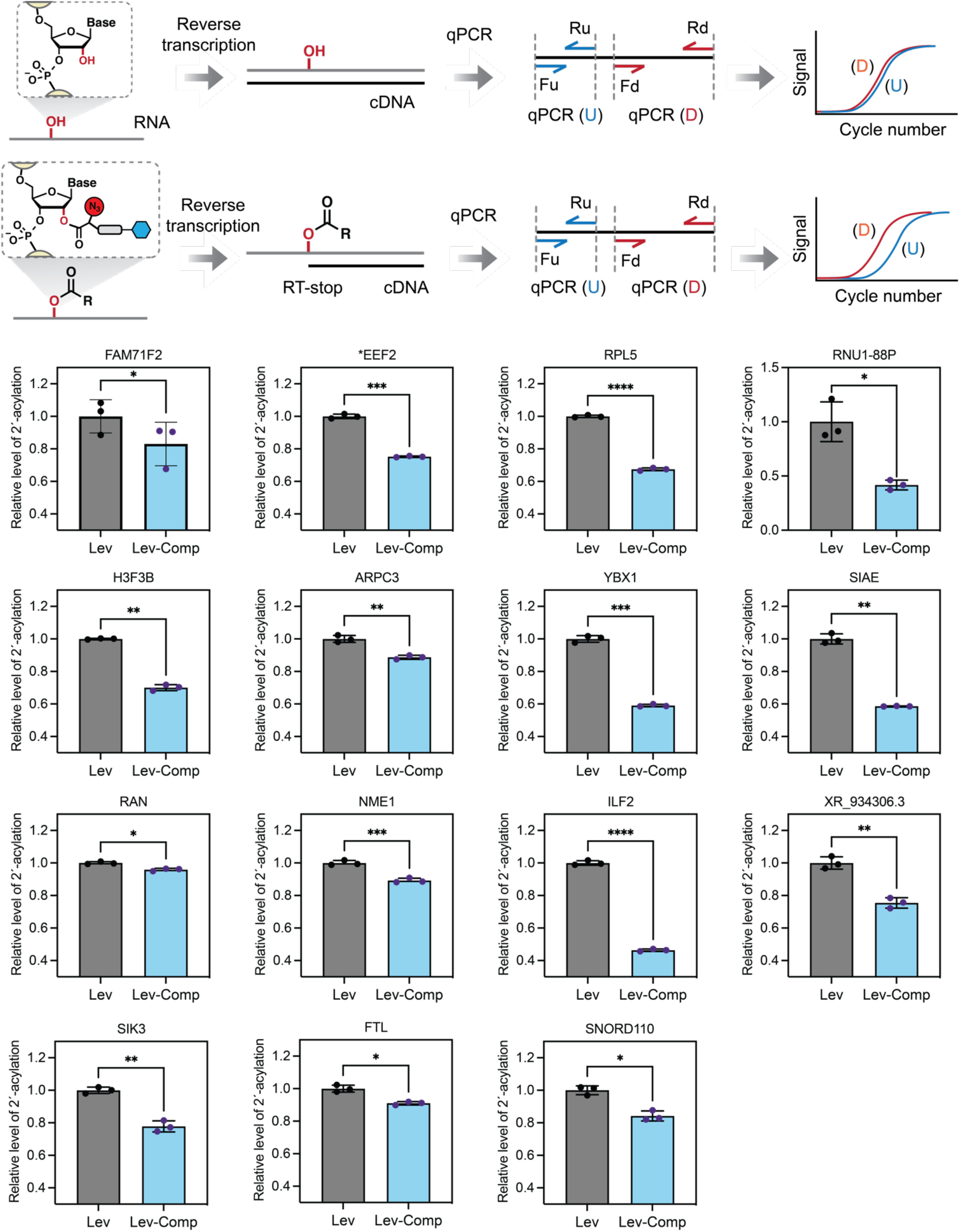
qPCR independently validates the transcriptome interactions of Levofloxacin (Lev) at 15 transcriptome binding sites in HEK293 cells. Workflow showing the strategies of validating competable 2’-OH acylation sites with qPCR (Top panel). qPCR validated and quantified the relative level of 2’-OH acylation at the drug-binding loci.

**Extended Data Fig. 5.**
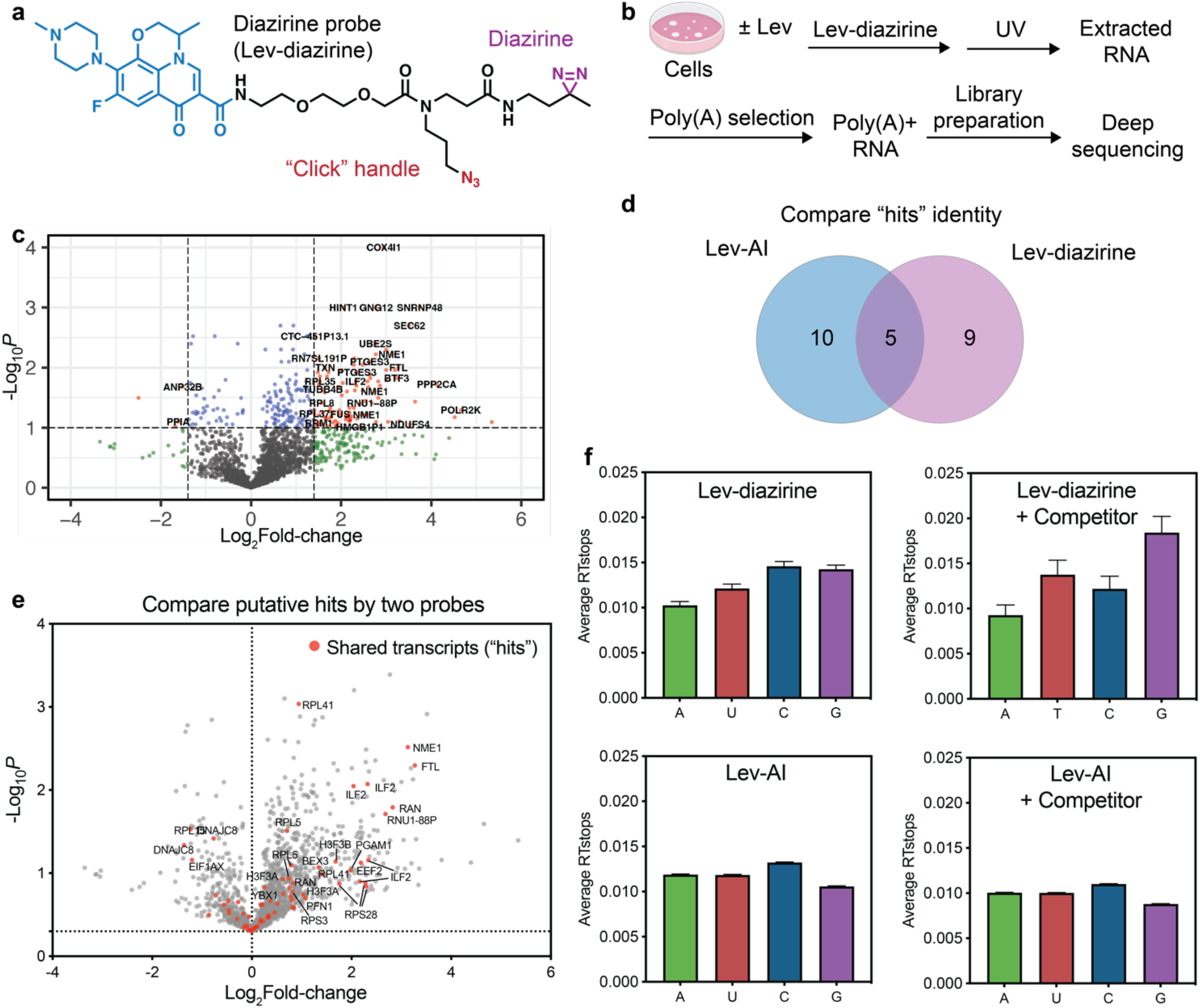
RBRP complements the existing diazirine-based profiling method. **a**, Chemical structure of diazirine-conjugated analog of Levofloxacin (Lev-diazirine). The drug moiety is colored blue, “click” handle” is colored red, and diazirine moiety is colored purple. **b**, Workflow for profiling RNA targets of Lev with Lev-diazirine probe. **c**, Volcano plot showing transcripts that are confidently enriched by Lev-diazirine in HEK293 cells. **d**, Volcano plot showing shared RNA targets that are identified by both Lev-AI and Lev-diazirine. X-axis: log-transformed ratio of transcript abundance in the presence over the absence of unmodified drug. Y-axis: the negative value of log-transformed *P*-value calculated with DESeq2. **d**, Venn diagram comparing RNA targets that were identified by Lev-AI and Lev-diazirine. **f**, Bar plots comparing the reactivities of Lev-AI and Lev-diazirine towards four nucleotides.

**Extended Data Fig. 6.**
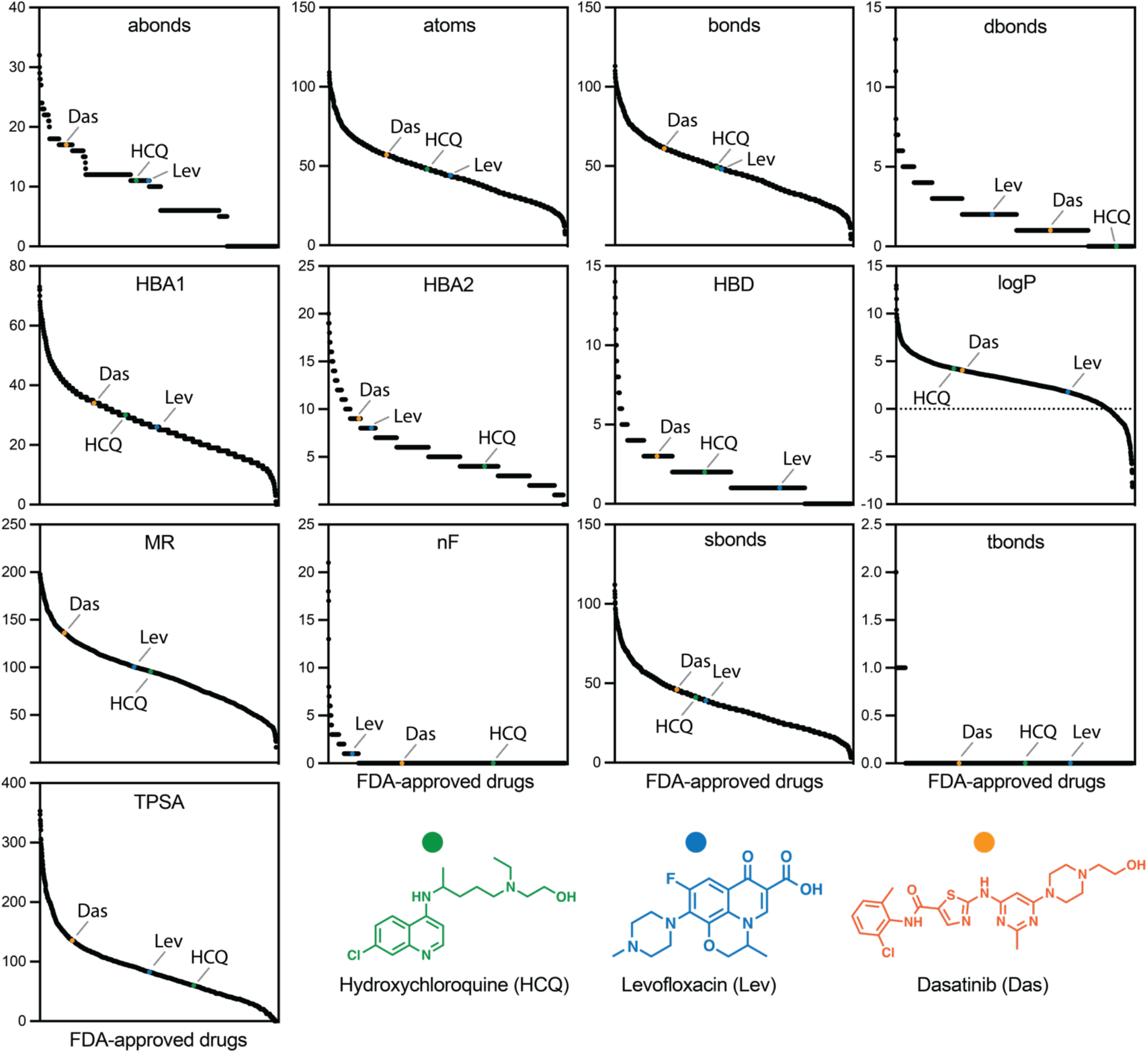
Plots comparing the structural fingerprints (Open Babel descriptors) of HCQ, Lev, and Das to an acquired list of FDA-approved small-molecule drugs (2076 drugs). Open Babel descriptors are abonds (Number of aromatic bonds), atoms (Number of atoms), bonds (Number of bonds), dbonds (Number of double bonds), HBA1 (Number of Hydrogen Bond Acceptors 1), HBA2 (Number of Hydrogen Bond Acceptors 2), HBD (Number of Hydrogen Bond Donors), logP (Octanol/water partition coefficient), MR (Molar refractivity), MW (Molecular Weight), nF (Number of Fluorine Atoms), sbonds (Number of single bonds), tbonds (Number of triple bonds), and TPSA (Topological polar surface area).

**Extended Data Fig. 7.**
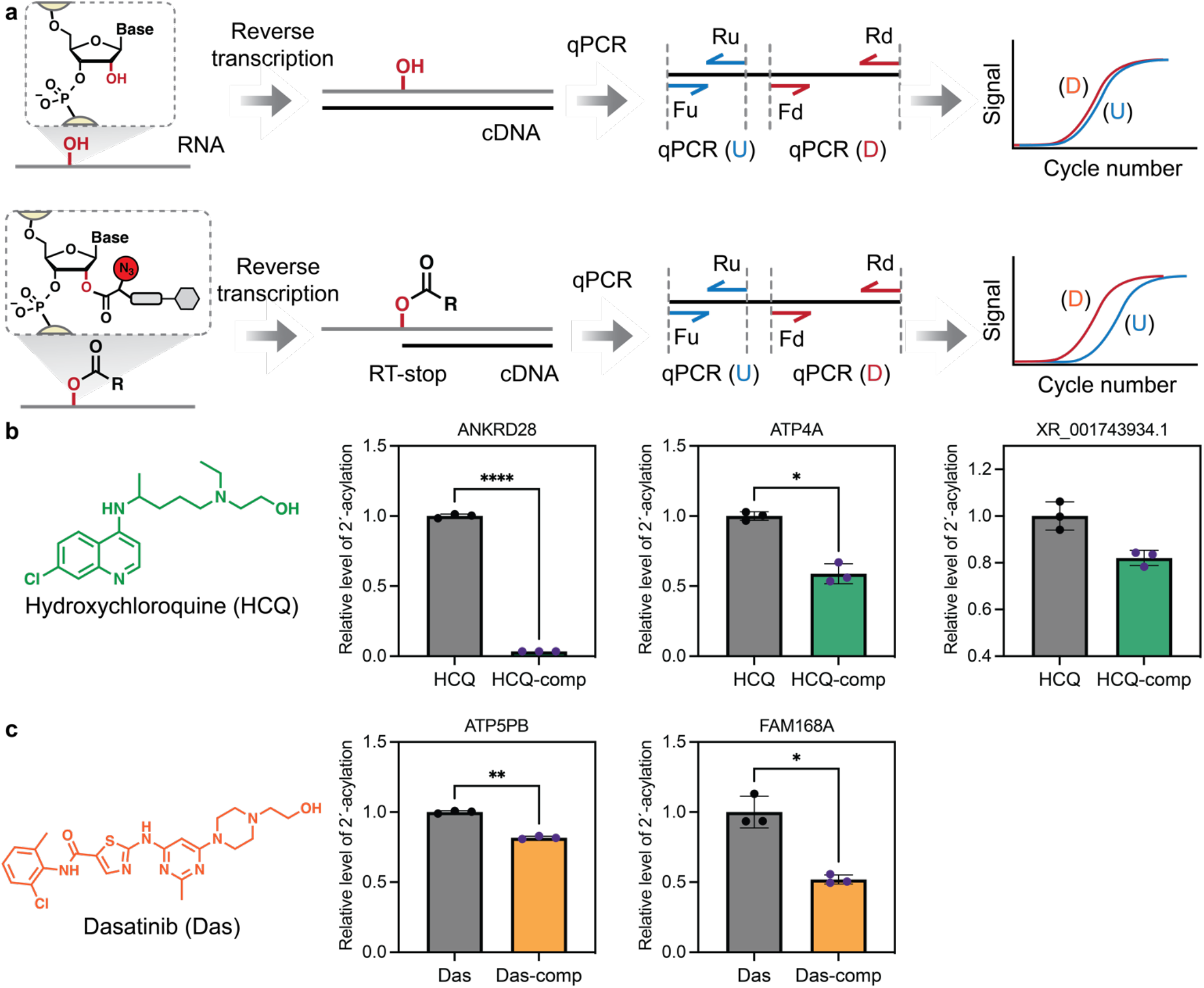
qPCR independently validates the transcriptome interactions of Hydroxychloroquine (HCQ) and Dasatinib (Das) at several transcriptome binding sites in HEK293 cells. **a**, Workflow showing the strategies of validating competable 2’-OH acylation sites with qPCR (Top panel). **b**-**c**, qPCR validated and quantified the relative level of 2’-OH acylation at the drug-binding loci of HCQ (**b**) and Das (**c**).

