## Supplementary Information for "Pervasive Transcriptome Interactions of Protein-Targeted Drugs"


- Supplementary Information Fig. 1-3 S2-S4
- Supplementary Table S1. Summary of selected principal components (PCs) for approved drugs. S5
- Supplementary Table S2. Eigenvalues, proportion of variance, and cumulative proportion of variance of all principal components. S5
- Supplementary Table S3. Summary of structural fingerprints of three drugs tested in this study. S6
- Supplementary Table S4. List of oligonucleotides used in this study S7
- Supplementary Table S5. List of reagents and materials S8-S9
- Preparation of sequencing library (detailed protocol) S10-S13
- Bioinformatics and command lines S14-S16
- Synthesis of acylimidazole probes S17-S27
- NMR spectra S28-S60
- HRMS data S61-65
- References S66

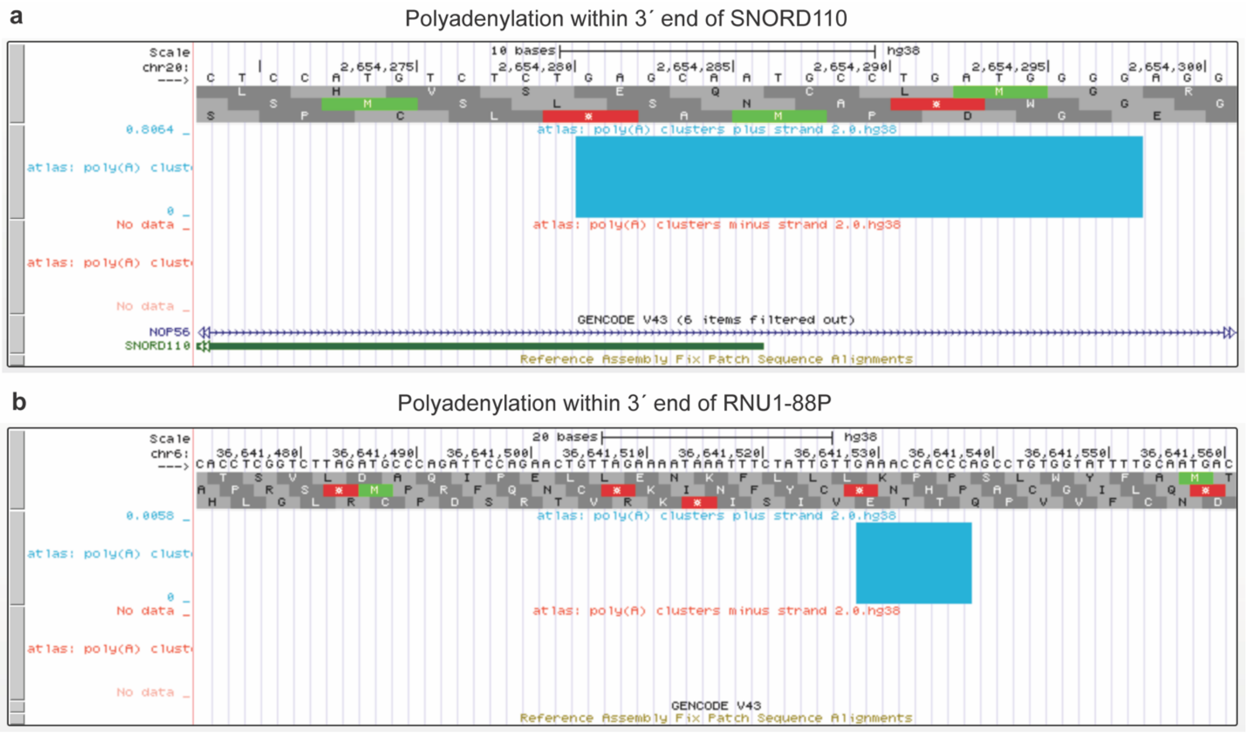


**Supplementary Information Fig. 1 | SNORD110 and RNU1-88P transcripts can be polyadenylated within their annotated 3´ ends**. **a**, UCSC track showing Poly(A) sites within the annotated 3´ end of SNORD110 (chr20:2654281-2654298; Positive strand). **b**, UCSC track showing Poly(A) sites within 3´ end of RNU1-88PP (chr6:36641529-36641538; Positive strand). The UCSC tracks were generated with PolyASite 2.0[1], which was constructed from publicly available human 3´ end sequencing datasets.

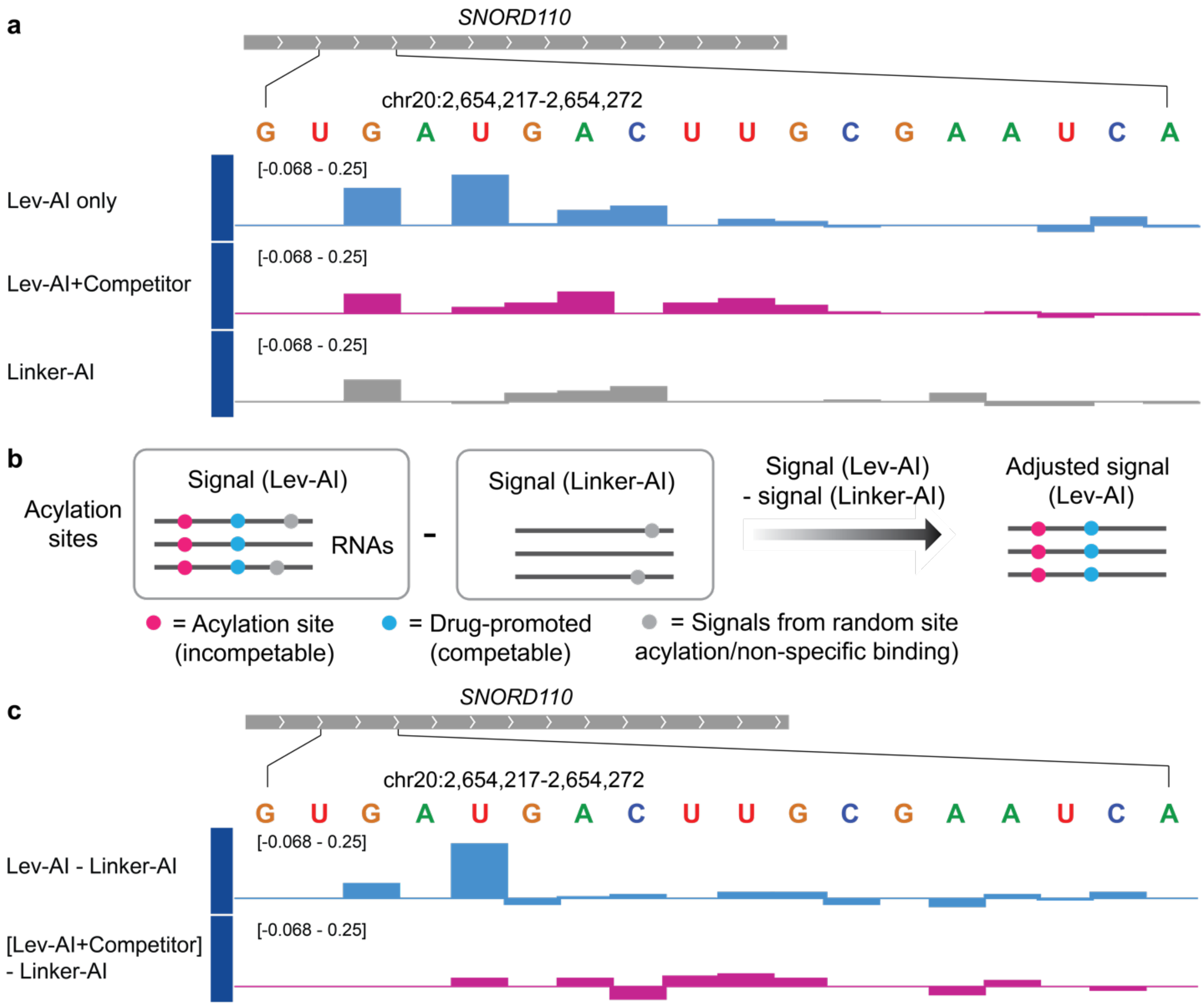


**Supplementary Information Fig. 2 | Deep sequencing validates selective SNORD110 interaction of Lev**. **a**, UCSC tracks showing calculated acylation yield (range from -0.068 to 0.25; an acylation yield 1.0 represents 100% 2´-OH acylation) at an RNA locus in SNORD110-expressing sequence by Lev-AI (top, colored blue), Lev-AI with competitor (middle, colored magenta), and Linker-AI (bottom, colored grey). **b**, Bioinformatics strategies to remove background signals that stem from random site 2´-OH acylation by Linker-AI or non-specific binding during sequencing library preparation. **c**, UCSC tracks showing adjusted acylation yield (range from -0.068 to 0.25; an acylation yield 1.0 represents 100% 2´-OH acylation) in SNORD110-expressing sequence. Acylation yields of Linker-AI were subtracted from those of Lev-AI and Lev-AI with competitor. Our data suggest that background acylation signals by Linker-AI can be removed by this bioinformatics workflow. For example, RBRP signal at G3 in the sequence shown was fully removed. This analysis also confirms RBRP scores are due to Lev-AI selectively targets SNORD110 rather than background acylation by the linker structure of Lev-AI.

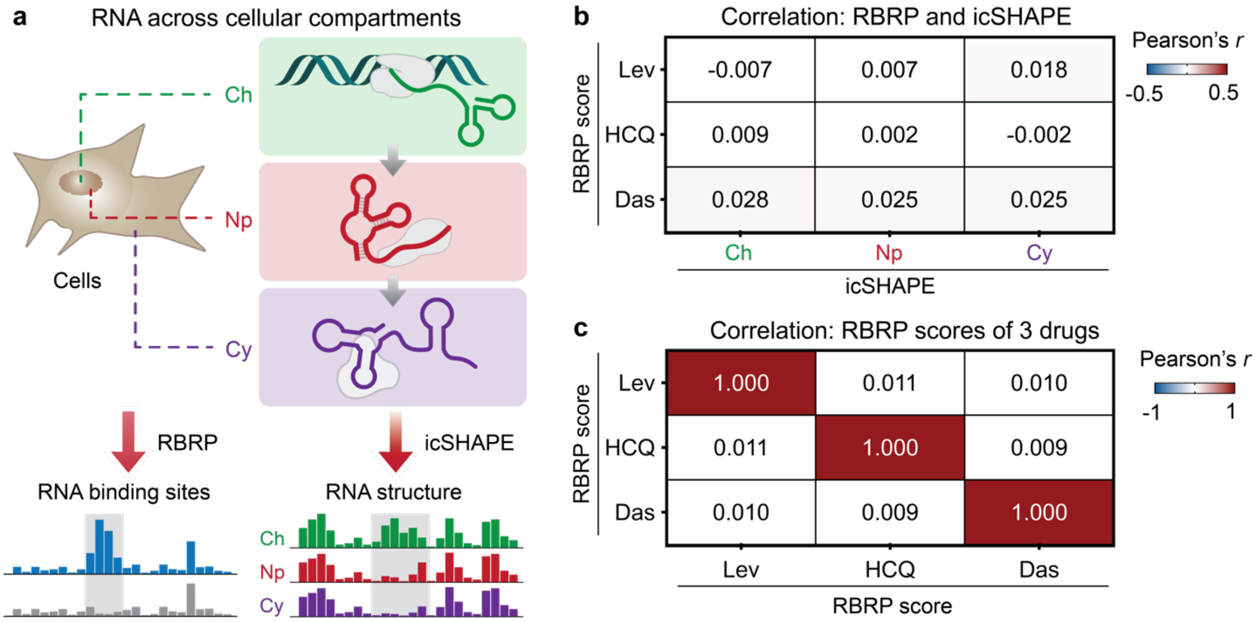


**Supplementary Information Fig. 3 | Meta-analysis reveals a minimal correlation between RBRP score and RNA structure transcriptome-wide. a**, Meta-analysis to quantify correlation between RBRP score and RNA structural accessibility (icSHAPE). Ch, chromatin. Np, nucleoplasm. Cy, cytoplasm. **b**, Heatmap showing minimal Pearson correlation between RBRP scores and icSHAPE reactivities in three cellular compartments transcriptome wide. **c**, Heatmap showing minimal Pearson correlation among RBRP scores of three drugs (Lev, HCQ, and Das) transcriptome wide.

**Supplementary Table S1.** Summary of selected principal components (PCs) for approved drugs (*n*=1,801).

| Parameter | PC1 | PC2 | Notes |
|---|---|---|---|
| abonds | 0.18 | -0.33 | Number of aromatic bonds |
| atoms | 0.97 | -0.20 | Number of atoms |
| bonds | 0.97 | -0.21 | Number of bonds |
| dbonds | 0.43 | 0.33 | Number of double bonds |
| HBA1 | 0.94 | -0.05 | Number of Hydrogen Bond Acceptors 1 |
| HBA2 | 0.58 | 0.71 | Number of Hydrogen Bond Acceptors 2 |
| HBD | 0.28 | 0.77 | Number of Hydrogen Bond Donors |
| logP | 0.34 | -0.81 | Octanol/water partition coefficient |
| MR | 0.94 | -0.25 | Molar refractivity |
| MW | 0.93 | -0.02 | Molecular Weight |
| nF | 0.03 | -0.19 | Number of Fluorine Atoms |
| sbonds | 0.90 | -0.13 | Number of single bonds |
| tbonds | -0.01 | -0.11 | Number of triple bonds |
| TPSA | 0.45 | 0.83 | Topological polar surface area |

**Supplementary Table S2.** Eigenvalues, proportion of variance, and cumulative proportion of variance of all principal components.

| | PC1 | PC2 | PC3 | PC4 | PC5 | PC6 | PC7 |
|---|---|---|---|---|---|---|---|
| Eigenvalue | 6.262 | 2.86 | 1.353 | 0.9938 | 0.9867 | 0.8903 | 0.3473 |
| Proportion | 44.73% | 20.43% | 9.67% | 7.10% | 7.05% | 6.36% | 2.48% |
| Cumulative proportion | 44.73% | 65.16% | 74.82% | 81.92% | 88.97% | 95.33% | 97.81% |

| | PC8 | PC9 | PC10 | PC11 | PC12 | PC13 |
|---|---|---|---|---|---|---|
| Eigenvalue | 0.1553 | 0.08583 | 0.03988 | 0.01611 | 0.008844 | 0.000571 |
| Proportion | 1.11% | 0.61% | 0.28% | 0.12% | 0.06% | 0.004% |
| Cumulative proportion | 98.92% | 99.53% | 99.82% | 99.93% | 100.00% | 100.00% |

**Supplementary Table S3.** Summary of structural fingerprints of three drugs tested in this study.

| Compound | abonds | atoms | bonds | dbonds | HBA1 | HBA2 | HBD |
|---|---|---|---|---|---|---|---|
| Levofloxacin | 11 | 46 | 49 | 2 | 26 | 7 | 1 |
| Dasatinib | 17 | 59 | 62 | 1 | 35 | 9 | 3 |
| Hydroxychloroquine | 11 | 49 | 50 | 0 | 30 | 4 | 2 |

| Compound | logP | MR | MW | nF | sbonds | tbonds | TPSA |
|---|---|---|---|---|---|---|---|
| Levofloxacin | 1.547 | 101.83 | 361 | 1 | 36 | 0 | 75.01 |
| Dasatinib | 3.462 | 138.63 | 488 | 0 | 44 | 0 | 134.75 |
| Hydroxychloroquine | 3.856 | 98.57 | 336 | 0 | 39 | 0 | 48.39 |

**Supplementary Table S4. List of oligonucleotides used in this study**

| **Oligo Nucleotides** | | |
|---|---|---|
| ***Primers for sequence library preparation*** | | |
| Probe-treated RNAs require a pre-adenylylated and 3′-ddC blocked RNA linker | IDT | (Probe-treated samples, HPLC-purified): /5rApp/AGATCGGAAGAGCGGTTCAG/3ddC/ |
| *DMSO-treated RNAs* require a pre-adenylylated and 3′-biotin blocked RNA linker | IDT | (DMSO samples, HPLC-purified): /5rApp/AGATCGGAAGAGCGGTTCAG/3Biotin/ |
| P5-universal PCR primer | IDT | (HPLC-purified) AATGATACGGCGACCACCGAGATCTACACTCTTTCCCTACACGACGCTCTTCCGATCT |
| P3-universal PCR primer | IDT | (HPLC-purified) CAAGCAGAAGACGGCATACGAGATCGGTCTCGGCATTCCTGCTGAACCGCTCTTCCGATCT |
| RT primer-1 | IDT | (Standard desalting) /5phos/DDDNNAACCNNNNAGATCGGAAGAGCGTCGTGGA/iSp18/GGATCC/iSp18/TACTGAACCGC |
| RT primer-2 | IDT | (Standard desalting) /5phos/DDDNNACAANNNNAGATCGGAAGAGCGTCGTGGA/iSp18/GGATCC/iSp18/TACTGAACCGC |
| RT primer-3 | IDT | (Standard desalting) /5phos/DDDNNAGGTNNNNAGATCGGAAGAGCGTCGTGGA/iSp18/GGATCC/iSp18/TACTGAACCGC |
| RT primer-4 | IDT | (Standard desalting) /5phos/DDDNNATTGNNNNAGATCGGAAGAGCGTCGTGGA/iSp18/GGATCC/iSp18/TACTGAACCGC |
| RT primer-5 | IDT | (Standard desalting) /5phos/DDDNNGCCANNNNAGATCGGAAGAGCGTCGTGGA/iSp18/GGATCC/iSp18/TACTGAACCGC |
| RT primer-6 | IDT | (Standard desalting) /5phos/DDDNNGACCNNNNAGATCGGAAGAGCGTCGTGGA/iSp18/GGATCC/iSp18/TACTGAACCGC |
| RT primer-7 | IDT | (Standard desalting) /5phos/DDDNNGATTNNNNAGATCGGAAGAGCGTCGTGGA/iSp18/GGATCC/iSp18/TACTGAACCGC |
| RT primer-8 | IDT | (Standard desalting) /5phos/DDDNNTTGANNNNAGATCGGAAGAGCGTCGTGGA/iSp18/GGATCC/iSp18/TACTGAACCGC |
| RT primer-9 | IDT | (Standard desalting) /5phos/DDDNNCTGGNNNNAGATCGGAAGAGCGTCGTGGA/iSp18/GGATCC/iSp18/TACTGAACCGC |
| RT primer-10 | IDT | (Standard desalting) /5phos/DDDNNTCCTNNNNAGATCGGAAGAGCGTCGTGGA/iSp18/GGATCC/iSp18/TACTGAACCGC |
| RT primer-11 | IDT | (Standard desalting) /5phos/DDDNNGGAANNNNAGATCGGAAGAGCGTCGTGGA/iSp18/GGATCC/iSp18/TACTGAACCGC |
| RT primer-12 | IDT | (Standard desalting) /5phos/DDDNNCAAGNNNNAGATCGGAAGAGCGTCGTGGA/iSp18/GGATCC/iSp18/TACTGAACCGC |
| ***Primers for target validation and cloning*** | | |
| Please see **Supplementary file S4** for details. | | |

**Supplementary Table S5**. **List of all reagents and materials**

| REAGENTS | SOURCE | NOTES |
|---|---|---|
| **Reagents and Enzymes** | | |
| Trizol LS Reagent | Thermo Scientific | # 10296028 |
| Dulbecco's Modified Eagle Medium (DMEM) | gibco | # 11995-065 |
| 1xPBS, pH=7.4 | gibco | # 10010-023 |
| UltraPure DNase/RNase-free Distilled water | Thermo Scientific | # 10977023 |
| 96% Ethanol | Fisher | # BP8202-500 |
| Fetal bovine serum (FBS) | gibco | # 26140-079 |
| RNA Gel loading dye (2x) | Thermo Scientific | # R0641 |
| SYBR Gold Nucleic Acid gel stain (10,000X) | Thermo Scientific | # S11494 |
| SYBRGreen I Nucleic Acid gel stain (10,000X) | Thermo Scientific | # S7563 |
| RiboLock RNase Inhibitor (40 U/µL) | Thermo Scientific | # EO0381 |
| SuperScript III Reverse Transcriptase | Thermo Scientific | # 18080044 |
| Gel Loading Dye, Orange (6X) | New England Biolabs | # B7022S |
| Rabbit Reticulocyte Lysate, Nuclease-Treated | Promega | # L4960 |
| Luna Universal qPCR Master Mix | New England Biolabs | # M3003L |
| dNTP Mix (10 mM each) | Thermo Scientific | # R0191 |
| DIBO-Biotin | Millipore Sigma | # 760749 |
| DTT (dithiothreitol) | Thermo Scientific | # R0861 |
| NaCl (5 M), RNase-free | Thermo Scientific | # AM9759 |
| EDTA (0.5 M), pH 8.0, RNase-free | Thermo Scientific | # AM9260G |
| Tween-20 | Millipore Sigma | # P7949 |
| CircLigase™ II ssDNA Ligase | VWR | # 76081-610 |
| Phusion High-Fidelity PCR Master Mix with HF Buffer | New England BioLabs | # M0531S |
| T4 Polynucleotide Kinase (T4 PNK) | New England BioLabs | # M0201S |
| FastAP Thermosensitive Alkaline Phosphatase (1 U/µL) | Thermo Scientific | # EF0651 |
| RNA Fragmentation Reagents | Thermo Scientific | # AM8740 |
| TopVision Agarose | Thermo Scientific | # R0491 |
| **Cell Lines** | | |
| HEK293 | ATCC | # CRL-1573 |
| **Chemicals** | | |
| Chloroform | Fisher Scientific | # AC610281000 |
| 1,1'-Carbonyldiimidazole (CDI) | AK Scientific | # A069 |

| | | |
|---|---|---|
| UltraPure 1 M Tris-HCl buffer, pH=7.5 | Thermo Scientific | # 15567-027 |
| d6-DMSO | ACROS | # 320770075 |
| Hydroxychloroquine sulfate | AstaTech | # H10433 |
| Dasatinib | AstaTech | # 62242 |
| Levofloxacin | AURUM Pharmatech | # U22961 |
| **Commercial Kits** | | |
| Quick-RNA Midiprep kit | Zymo | # R1056 |
| RNA Cleanup & Concentrator Column-5 | Zymo | # R1014 |
| MinElute Gel Extraction Kit | QIAGEN | # 28604 |
| Poly(A)Purist MAG kit | Thermo Scientific | # AM1922 |
| RNeasy Mini kits | QIAGEN | # 74904 |
| Dynabeads™ MyOne™ Streptavidin C1 | Thermo Scientific | # 65001 |
| Amicon Ultra 10K filter | Millipore Sigma | # UFC501024 |
| Zymo DNA Clean & Concentrator Column-5 | Zymo | # D4014 |
| **Software & online source** | | |
| GraphPad Prism 9 | GraphPad Software | https://www.graphpad.com/ |
| ImageStudioLite | Li-COR biosciences | https://www.licor.com/bio/ |
| Adobe Illustrator 2021 | Adobe | https://www.adobe.com/ |
| MestReNova | Mestrelab | https://www.mestrelab.com/ |
| DrugBank | DrugBank | https://go.drugbank.com/ |
| Drug Repurposing Hub | Broad Institute | https://www.broadinstitute.org/drug-repurposing-hub |
| PubChem Score Matrix Service | NIH | https://pubchem.ncbi.nlm.nih.gov/score_matrix/score_matrix.cgi |
| ChemMine Tools | University of California Riverside | https://chemminetools.ucr.edu/ |
| icSHAPE bioinformatics pipeline | Github | https://github.com/qczhang/icSHAPE |
| UCSC Genome Browser | UCSC Genomics Institute | https://genome.ucsc.edu/ |
| Integrative Genomics Viewer (IGV) | Broad Institute | https://software.broadinstitute.org/software/igv/ |
| wiggletools | Github | https://github.com/Ensembl/WiggleTools |

**Preparation of sequencing library**

**Activation of acylimidazole probes.** The acylimidazole probes were generated immediately before the profiling experiments. Specifically, carboxylic precursors of acylimidazole probes were treated with a slightly excess amount of 1,1'-carbonyldiimidazole (CDI) in dry DMSO to render activated acylimidazole solution, which contains 1:1.3 acylimidazole compound and imidazole. The acylimidazole solution was further used without purification.

To generate the Hydroxychloroquine-acylimidazole probe (HCQ-AI), a 5 µL solution of 100 mM HCQ-acid was treated with 5 µL of freshly prepared 130 mM CDI solution in dry d6-DMSO. The reaction mixture was incubated overnight to yield a solution of ~50 mM activated HCQ-AI.

To generate the Dasatinib-Acylimidazole probe (Das-AI), 1.7 mg of *Dasatinib-COOH* was dissolved in 20.4 µL of dry d6-DMSO and sequentially added 20.4 µL of 130 mM CDI in dry d6-DMSO. The reaction mixture was incubated under argon overnight to render a solution of ~50 mM activated Das-AI.

To generate the Levofloxacin-Acylimidazole probe (Lev-AI), 1.6 mg of Levofloxacin-COOH was dissolved in 45 µL of dry d6-DMSO and then treated with 5 µL of 617 mM CDI in dry d6-DMSO. The reaction mixture was incubated under argon overnight to yield a solution of ~47 mM activated Lev-AI.

**Cell culture, cell harvesting, and pretreatment with competitor drugs**. HEK293 cells were grown in antibiotic-free high-glucose DMEM containing 10% FBS on 15-cm plates until ~90% confluency. (a) To harvest cells for probe treatment in the absence of competing drugs, cells were washed with 10 mL of warm 1x PBS, pH=7.4 once. 4 mL of 1x PBS, pH=7.4 containing 2% DMSO was added to the cells. The cells were then gently scraped, and ~2x$10^6$ cells were transferred to a 15-mL falcon tube. The supernatant was removed by centrifugation at 800*g* for 2 minutes at room temperature. (b) To harvest cells for probe treatment in the presence of competitor drug, cells were pretreated with 10 mL of DMEM containing 10% FBS and Hydroxychloroquine (300 µM), Dasatinib (200 µM), or Levofloxacin (300 µM) for 30 minutes at 37 ˚C in a humidified incubator under 5% $CO_2$. After incubation, cells were washed with 10 mL warm 1xPBS, pH=7.4 containing Hydroxychloroquine (300 µM), Dasatinib (200 µM), or Levofloxacin (300 µM) once, then gently scraped into 4 mL of 1xPBS, pH=7.4 containing the indicated concentration of corresponding competitor drugs. ~2x$10^6$ cells were transferred to a 15-mL falcon tube. The supernatant was removed by centrifugation at 800*g* for 2 minutes at room temperature.

**Treatment of live cells with acylimidazole probes**. Cells were resuspended in 1 mL of 1xPBS containing 50 µM acylimidazole probe (HCQ-AI, Das-AI, or Lev-AI) or DMSO as a negative control. Cells were transferred to a 2-mL Eppendorf tube and was incubated at 37 ˚C on a sample revolver for 30 mins. The reaction was stopped by centrifuging samples at 2500g for 1 min at 4˚C.

**Cell lysis and isolation of total cellular RNA**. The cells were resuspended in 500 µL 1xPBS, pH=7.4, and then immediately transferred to a 15-mL falcon tube containing 6 mL of Trizol LS. The resulting sample was homogenized for ~15s by vortexing. Next, 1.2 mL of chloroform (0.2x by volume) was added to the solution and mixed for ~15s by vortexing. The resulting mixture was incubated at room temperature for 5 minutes and then centrifuged at 2500*g* for 15 minutes at 4˚C. The aqueous phase (*top*) was collected into a new RNase-free tube.

Next, total cellular RNA was isolated using a Zymo Quick-RNA Midiprep kit according to the manufacturer's protocol with on-column digestion of genomic DNA. Total cellular RNA was eluted with 400 µL of RNase-free water. The concentration of eluted RNA was determined by Nanodrop. RNA was then stored as 250-µg aliquots at -80 ˚C. RNA quality was assessed by 1% agarose gel (stained by 1x SYBR Gold). Clear 18S and 28S rRNA bands should be observed.

**Isolating poly(A)+ RNA fraction with Poly(A)Purist MAG kit.** A 250-µg aliquot of total cellular RNA was diluted to a final volume of 400 µL with RNA storage solution to reach a total concentration of 600 µg/mL.

Poly(A)+ RNAs were then purified with Poly(A)Purist MAG kit according to the manufacturer's protocol. Briefly, poly(A)+ RNAs were pulled down with beads containing poly(dT) and eluted with 200 µL of warm (60-80 ˚C) RNA storage solution twice.

To further desalt and cleanup the RNA, eluted RNA was first lyophilized and then dissolved in 100 µL of water. Next, the resulting RNA solution was purified by RNeasy Mini columns according to the manufacturer's protocol. Finally, RNA was eluted twice with 50 µL of water (100 µL final volume) and stored as 500-ng aliquots at -80˚C.

**Biotin conjugation of Probe-labeled RNAs by "click" chemistry**. RNA aliquots (~16 µL) were thawed and treated with 2 µL of 1.85 mM DIBO-Biotin (solution in DMSO) and 1 µL of RiboLock. The reaction was incubated at 37˚C for 2 hours on a sample revolver. After the reaction, the biotin-conjugated transcripts were purified with a Zymo RNA Clean & Concentrator column-5 according to the manufacturer's protocol. RNA was eluted with 15 µL of water and freeze-dry on a lyophilizer.

**RNA fragmentation and RNA end repair**. Biotin-conjugated RNAs were resuspended in 9 µL of RNase-free water and heated at 95 ˚C for 60 seconds. 1 µL of 10x RNA fragmentation reagent was then added to the heated RNA solution and incubated at 95 ˚C for 50 seconds, which rendered fragmented RNAs with a medium length of ~100 nt. The reaction was immediately quenched with 1 µL of 10x stop solution on ice. The resulting RNA fragments were then purified using a Zymo RNA cleanup & concentrator column-5 accordingly to the manufacturer's protocol. The eluted RNA fragments were then freeze-dried on a lyophilizer.

3'-end repair of RNA fragments was performed according to the previously reported protocol[2,3]. Briefly, RNA was solubilized in 10 µL of RNA end-repair mix that contains 1 µL of 10x T4 PNK buffer, 1 µL of RiboLock (U/µL), 1 µL of FastAP (1 U/µL), 2 µL of T4 PNK (10 U/µL), and 5 µL of RNase-free water. The reaction was incubated at 37˚C for 1 hour.

**RNA 3'-end ligation**. The 3'-end ligation was performed according to the previously reported protocol with modifications[2,3]. For probe-treated samples, 15.875 µL of the 3' RNA ligation mixture (as tabulated below) was added to each sample. The resulting reaction mixture was incubated at 25 ˚C for 4 hours in a thermocycler.

| **Component** | **Amount (µL)** |
|---|---|
| RNA ligase buffer, 10x | 1.5 |
| DTT, 200 mM | 0.625 |
| Preadenylylated and **3'-ddC** blocked RNA linker, 10 µM | 2.5 |
| T4 RNA ligase 1 (10,000 U/mL) | 3.75 |
| PEG8000, 50% (v/v) | 7.5 |
| Completed end-repair reaction | 10 |
| Total | 25.8 |

3′-ddC blocked RNA linker: (Probe-treated samples, IDT, HPLC-purified): /5rApp/AGA TCG GAA GAG CGG TTC AG/3ddC/

For DMSO-treated samples, 15.875 µL of the 3' RNA ligation mixture (as tabulated below) was added to each sample. The resulting reaction mixture was incubated at 25 ˚C for 4 hours in a thermocycler.

| **Component** | **Amount (µL)** |
|---|---|
| RNA ligase buffer, 10x | 1.5 |
| DTT, 200 mM (Fresh) | 0.625 |
| Preadenylylated and **3'-biotin** blocked RNA linker, 10 µM | 2.5 |
| T4 RNA ligase 1 (10,000 U/mL) | 3.75 |

| | |
|---|---|
| PEG8000, 50% (v/v) | 7.5 |
| Completed end-repair reaction | 10 |
| Total | 25.8 |

**3′-biotin** blocked RNA linker: (DMSO samples, IDT, HPLC-purified): /5rApp/AGA TCG GAA GAG CGG TTC AG/3Biotin/

After the 3'-end ligation reaction, RNA was purified with a Zymo RNA cleanup & concentrator column-5. The eluted RNA was freeze-dried on a lyophilizer. Next, RNA was redissolved in 7 µL of gel loading buffer II and resolved on a 6% (w/v) UreaGel denaturing PAGE and run at 20 W for 10 mins. The resulting gel was stained in SYBR Gold mix for 3 min at room temperature. The portions of gel with a length above ~50 nt were sliced, crushed, and transferred into 1.5-mL Eppendorf tubes. 250 µL of RNase-free water was added to the crushed gel slices and heated at 67 ˚C for 15 minutes on an orbital shaker. The extracted RNA solution was then desalted with a 0.5-mL 10K Amicon filter according to the manufacturer's protocol. The eluted RNA was then freeze-dried on a lyophilizer.

**cDNA synthesis, enrichment of Biotin-modified RNAs, and cDNA purification.** The following steps were performed according to the previously reported protocol with slight modifications[2,3]. The 3'-ligated RNA was solubilized in 11 µL of RNase-free water and 1 µL of 1 µM RT primer (see **Supplementary Table S4** for assignments of RT primers). Barcoded primers for reverse transcription: /5phos/DDDNN**AACC**NNNNAGATCGGAAGAGCGTCGTGGA/iSp18/GGATCC/iSp18/TACTGAACCGC. D = A/G/T and N = A/T/G/C. The underlined four nucleotides are barcodes.) The reaction was heated at 70 ˚C for 5 minutes and then cooled to 25 ˚C by 20˚C/60 seconds. 8 µL of RT reaction mixture (4 µL of 5x First strand buffer, 0.75 µL of RiboLock, 1 µL of 100 mM DTT, 1 µL of 10 mM dNTPs, and 1.25 µL of SuperScript III) was then added to the above reaction. The resulting solution was heated at 25 ˚C for 3 mins, 42 ˚C for 5 mins, and 52 ˚C for 30 mins.

Next, the biotin-conjugated RNAs were enriched with 5 µL of MyOne C1 magnetic streptavidin beads by incubating at 25˚C for 45 minutes on a sample revolver. The captured RNA was stringently washed with 400 µL wash buffer (100 mM Tris, pH=7.0, 4 M NaCl, 10 mM EDTA, 0.2% Tween-20) for five times and 500 µL of 1xPBS, pH=7.4 for two times. Next, the beads were resuspended in 50 µL of an RNase mixture ( 5 µL of 10x RNaseH buffer, 12.5 µL of 50 mM D-biotin, 1 µL of 1 µM P3-universal PCR primer, 1 µL RNase A/T1 enzyme mix, 1 µL of RNase H, and 30.5 mL of water). The reaction mixture was incubated at 37˚C for 30 mins on an orbital shaker. P3- PCR primer (IDT, HPLC-purified): 5′-CAAGCAGAAGACGGCATACGAGATCGGTCTCGGCATTCCTG CTGAACCGCTC TTCCGATCT-3′

After the reaction, 1 µL DMSO was added to the solution, and the resulting mixture was heated at 95˚C for 4 mins. The supernatant was collected, and cDNA was further purified with Zymo DNA Clean & Concentrator column-5 according to the manufacturer's protocol. cDNA was eluted with water and freeze-dried on a lyophilizer. cDNA was then solubilized in 7 µL Gel loading buffer II and resolved on a 6% (w/v) denaturing PAGE. Gel portions above ~80 nt were collected, crushed, and transferred into 1.5-mL Eppendorf tubes. 250 µL of RNase-free water was added to the crushed gel slices and heated at 67 ˚C for 15 minutes on an orbital shaker. The extracted cDNA solution was then desalted with a 0.5-mL 10K Amicon filter according to the manufacturer's protocol and further purified with a Zymo DNA Clean & Concentrator column-5. cDNA was eluted twice in 8.5 µL water (final ~16 µL).

**cDNA circularization and library amplification**. The purified cDNA was circularized with CircLigase II according to the manufacturer's protocol. Briefly, to ~16 µL solution of cDNA was added 2 mL of 2x CirLigase II buffer, 1 µL of 50 mM $MnCl_2$, and 1 µL CircLigase II. The reaction was incubated at 60 ˚C for 3 hours before purification with Zymo DNA Clean & Concentrator column-5 according to the manufacturer's protocol. Circularized cDNA was eluted in 20 µL water and then transferred to optical PCR tubes for qPCR.

To amplify the library, 20 µL of 2x Phusion HF PCR master mix, 0.4 µL of SYBR Green I (25x), 1.0 µL of 10 µM P5-universal PCR primer (5′-AATGATACGGCGACCACCGAGATCTACACTCTTTCCCTACACGACGCTCTTCCGATCT-3′ and 10 µM P3-universal PCR primer (5′-CAAGCAGAAGACGGCATACGAGATCGGTCTCGGCATTCCTGCTGAACCGCTCTTCCGATCT-3′). PCR amplification was monitored in real-time to avoid over-amplification. The PCR reaction was first denatured at 98 ˚C for 45 seconds, followed by cycles of denaturation (98 ˚C, 15 seconds), Anneal (65 ˚C, 20 seconds), and Extension (72 ˚C, 1 minute). The PCR products were purified with Zymo DNA Clean & Concentrator column-5 according to the manufacturer's protocol and then freeze-dried on a lyophilizer. Next, the DNA was redissolved in 6 µL of gel loading buffer and resolved on 3% low melting point agarose in 1 % TBE buffer containing 1x SYBRGold. The gel slices corresponding to the PCR products were then extracted using the QIAGEN MiniElute Gel extraction kit. The purified DNA was analyzed by BioAnalyzer for quality control and quantified with Qubit.

**Bioinformatics.** We use a modified bioinformatics pipeline based on icSHAPE[2,3] to calculate averaged frequency of reverse transcription stops as the RBRP reactivity at each nucleotide transcriptome-wide. Step-wise protocols are as follows:

1. Raw sequencing data was unzipped and changed to FASTQ format.

```
$ gunzip RBRP.fq.gz
$ mv RBRP.fq RBRP.fastq
```

### this will change the data format to .fastq for barcode splitting in the next step.

2. The sequencing data were demultiplexed based on the barcode of each sub-library. The following example shows how to demultiplex raw data for a library consisting of ten sub-libraries.

```
$ splitFastq.pl -U RBRP.fastq -l GGTT:LIB-1::TTGT:LIB-2::ACCT:LIB-3::CAAT:LIB-
4::TGGC:LIB-5::GGTC:LIB-6::AATC:LIB-7::TCAA:LIB-8::CCAG:LIB-9::AGGA:LIB-
10::others:unmatched -b 5:4 -d library_split -s hiseq_barcode.stat
```

Barcode-1: GGTT (AACC in RT primer)
Barcode-2: TTGT (ACAA in RT primer)
Barcode-3: ACCT (AGGT in RT primer)
Barcode-4: CAAT (ATTG in RT primer)
Barcode-5: TGGC (GCCA in RT primer)
Barcode-6: GGTC (GACC in RT primer)
Barcode-7: AATC (GATT in RT primer)
Barcode-8: TCAA (TTGA in RT primer)
Barcode-9: CCAG (CTGG in RT primer)
Barcode-10: AGGA (TCCT in RT primer)

3. The quality of reads were evaluated with FastQC for each sub-libraries.

```
$ fastqc -o out_directory LIB_x.fastq
```

4. All sequencing reads were collapsed and PCR duplicates were removed.

```
$ readCollapse.pl -U LIB_x.fastq -o LIB_x.rmdup.fastq -f LIB_x.fa
```

### -LIB_x.fastq is the input file from Step 3; LIB_x.rmdup.fastq is the output file after removing PCR duplicates; LIB_x.fa records the frequency of each sequence in the input LIB_x.fastq file.

5. The adapter and barcode sequences were trimmed from reads.

```
$ trimming.pl -U LIB_x.rmdup.fastq -o LIB_x.trimmed.fastq -l 13 -t 0 -c phred33
-a adapter.fa -m 25
```

6. The trimmed reads were aligned to the human transcriptome with bowtie 2.

```
$ bowtie2 -U LIB_x.trimmed.fastq -S LIB_x.sam -x /PATH/transcripts --non-
deterministic --time
```

7. Abundance of identified transcripts were calculated.

```
$ estimateRPKM.pl -i LIB_x.sam -o LIB_x.rpkm
```

8. RT stop frequencies at each nucleotide in each transcript were quantified.

```
$ calcRT.pl -i LIB_x.sam -o LIB_x.rt -r LIB_x.rpkm -c 1
```

9a. The background RT stop frequencies were calculated by combining two replicates of DMSO-treated libraries.

```
$ combineRTreplicates.pl -i DMSO-1.rt:DMSO-2.rt -o DMSO.rt
```

9b. The RT stop frequencies of each sequence sub-library were normalized.

```
$ normalizeRTfileScaleFactor1.pl -i LIB_x.rt -o LIB_x.normalized.rt -d 32 -l 32
```

9c. RBRP score at each nucleotide for all transcripts was calculated.

```
$ calcRBRPminusCtrl.pl -f LIB_x.normalized.rt -b LIB_DMSO.normalized.rt -o LIB_x.tmp.out -e dividing -y 0.5
```

10. For probe-treated libraries, filter RBRP score based on foreground RT stops (in the probe-treated samples) and the background sequencing depth (background base density in DMSO-treated samples):

```
$ filterRTstopBD.pl -i LIB_x.tmp.out -o LIB_x.out -t 200 -s 5 -e 30
```

11. Bedgraph files were generated for each sub-library to visualize RBRP score transcriptome-wide.

```
$ enrich2Bedgraph.pl -i LIB_x.out -o LIB_x.bedgraph -g /PATH/Homo_sapiens.GRCh38.104.gtf -a /PATH/transcriptome.fa
```

12. Bigwig files were generated for each sub-library.

```
$ sort -k1,1 -k2,3n LIB_x.bedgraph -o LIB_x.sorted.bedgraph
$ uniqueTrack.pl LIB_x.sorted.bedgraph LIB_x.sorted.uniq.bedgraph
$ cut -f1-4 LIB_x.sorted.uniq.bedgraph | grep -v NULL > LIB_x.sim.bedgraph
$ bedGraphToBigWig LIB_x.sim.bedgraph /PATH/genome.size LIB_x.sim.bw
```

**Data Visualization.** The following codes generate the final bigwig files that show the RBRP scores and -Log*P*-value from Welch's *t*-test. Lev is used as an example to show these codes below.

1. Generate mean track of Linker:

```
$ wiggletools write mean_Linker.wig mean Linker-1.sim.bw Linker-2.sim.bw
$ wigToBigWig mean_Linker.wig genome.size mean_Linker.sim.bw
```

2. Adjust each replicate of Lev:

```
$ wiggletools write adj.Lev-1.wig diff Lev-1.sim.bw mean_Linker.sim.bw
$ wiggletools write adj.Lev-2.wig diff Lev-2.sim.bw mean_Linker.sim.bw
$ wigToBigWig adj.Lev-1.wig genome.size adj.Lev-1.sim.bw
$ wigToBigWig adj.Lev-2.wig genome.size adj.Lev-2.sim.bw
```

3. Adjust each replicate of Lev-comp:

```
$ wiggletools write int.Lev-comp-1.wig diff Lev-comp-1.sim.bw mean_Linker.sim.bw
$ wiggletools write adj.Lev-comp-1.wig sum int.Lev-comp-1.wig deltaExp-Lev.sim.bw
$ wiggletools write int.Lev-comp-2.wig diff Lev-comp-2.sim.bw mean_Linker.sim.bw
$ wiggletools write adj.Lev-comp-2.wig sum int.Lev-comp-2.wig deltaExp-Lev.sim.bw
$ wigToBigWig adj.Lev-comp-1.wig genome.size adj.Lev-comp-1.sim.bw
$ wigToBigWig adj.Lev-comp-2.wig genome.size adj.Lev-comp-2.sim.bw
```

4. Calculate *P*-value with Welch *t*-test:

```
$ wiggletools write p-value.wig ttest adj.Lev-1.sim.bw adj.Lev-2.sim.bw : adj.Lev-comp-1.sim.bw adj.Lev-comp-2.sim.bw
$ wiggletools write logP-value.wig log 10 p-value.wig
$ wiggletools write neglog10P-value.wig scale -1 logP-value.wig
$ wigToBigWig neglog10P-value.wig genome.size neglog10P-value.sim.bw
```

5. Generate mean track of each condition:

```
$ wiggletools write mean_adj.Lev.wig mean adj.Lev-1.sim.bw adj.Lev-2.sim.bw
$ wiggletools write mean_adj.Lev-comp.wig mean adj.Lev-comp-1.sim.bw adj.Lev-comp-2.sim.bw
$ wigToBigWig mean_adj.Lev.wig genome.size mean_adj.Lev.sim.bw
$ wigToBigWig mean_adj.Lev-comp.wig genome.size mean_adj.Lev-comp.sim.bw
```

6. Generate differential track of mean_adj.Lev - mean_adj.Lev-comp using Wiggletools:

```
$ wiggletools write diff_RBRP.wig diff mean_adj.Lev.sim.bw mean_adj.Lev-comp.sim.bw
$ wigToBigWig diff_RBRP.wig genome.size diff_RBRP.sim.bw
```

##### Synthesis of acylimidazole probes

**General Synthetic Procedures**. All chemicals purchased from commercial suppliers were used without further purification, as described in Supplementary Table 5. The purities of compounds were determined by NMR and/or LC/MS. The NMR spectra were recorded on Varian 300, 400, or 500 MHz NMR spectrometers at ambient temperature. Chemical shifts were reported in parts per million (ppm) and coupling constants in hertz (Hz). $^1$H-NMR spectra were referenced to the residual solvent peaks as internal standards (2.50 ppm for d6-DMSO; 7.26 ppm for $CDCl_3$).

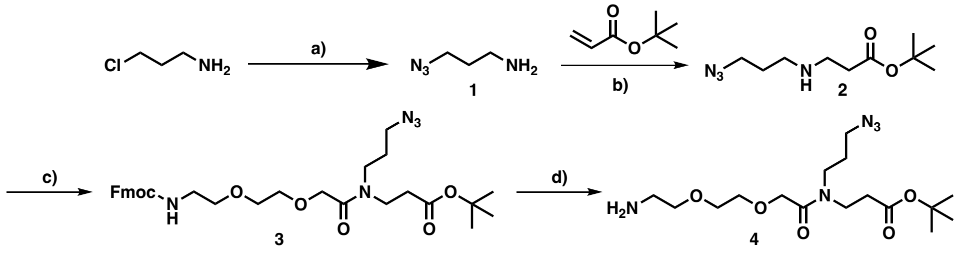


**Scheme 1 | Synthesis of intermediate 4.** a) Sodium azide, water, 80˚C, 15 hours. b) DBU, acetonitrile, room temperature, 20 hours. c) EDCI-HCl, HOBT, DIPEA, DMF, 0 ˚C, 12 hours. d) 20% piperidine in dichloromethane, room temperature, 1 hour.

###### Synthesis of 3-azidopropan-1-amine (1)

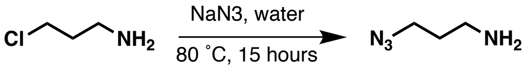


3-azidopropan-1-amine was synthesized according to a previously reported protocol (PMID: 31644285). Briefly, 5.0 g of 3-chloropropan-1-amine hydrochloride (1.0 equiv.) was dissolved in 50 mL $H_2O$ and 7.5 g of $NaN_3$ (3.0 equiv.) was slowly added. After heating at 80°C for 15 h, the reaction mixture was cooled to room temperature and 5.5 g of KOH pellets were added on ice, at which the orange color disappeared. After extracting with diethyl ether and drying with $MgSO_4$, the solvent was evaporated, yielding *3-azidopropan-1-amine* (**1**) as a clear liquid (2.1 g, 55%). $^1$H-NMR (500 MHz, $D_2O$) δ=3.42 (t, *J* = 6.9 Hz, 2H), 2.72 (td, *J* = 6.9, 1.3 Hz, 2H), 1.76 (ddd, *J* = 8.3, 6.6, 1.4 Hz, 2H). Note: The $^1$H-NMR is consistent with the reported data (PMID: 31644285), thus was used in the next step without further characterization.

###### Synthesis of *tert*-butyl 3-((3-azidopropyl)amino)propanoate (2)

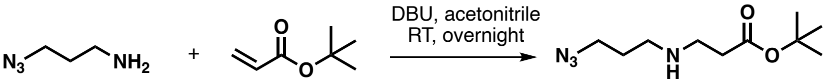


3-Azidopropan-1-amine (**1**) (1.7 g, 1.2 equiv.), *tert*-butyl acrylate (1.9 g, 1.0 equiv.), and DBU (1.1 g, 0.50 equiv.) were dissolved in 30 mL acetonitrile and stirred for 20 hours at room temperature under argon. Reaction progress was monitored by measuring the consumption of *tert*-butyl acrylate by NMR. After completion, the reaction mixture was concentrated in vacuum. Purification by silica column chromatography (eluent gradient: 0% to 10% methanol in dichloromethane) rendered *tert*-butyl 3-((3-azidopropyl)amino)propanoate (**2**) as a clear yellow liquid (1.5 g, 44%). Rf=0.31 in 10% methanol in dichloromethane. $^1$H-NMR (300 MHz, $CDCl_3$) δ= 3.35 (t, *J* = 6.7 Hz, 2H), 2.82 (t, *J* = 6.4 Hz, 2H), 2.69 (t, *J* = 6.9 Hz, 2H), 2.41 (t, *J* = 6.4 Hz, 2H), 1.75 (p, *J* = 6.8 Hz, 2H), 1.43 (s, 9H). $^{13}$C NMR (101 MHz, $CDCl_3$) δ= 171.92, 80.34, 49.26, 46.44, 44.94, 35.50, 28.96, 27.88. HRMS $C_{10}H_{20}N_4O_2$ $[M+H]^+$ calculated 229.1659, found 229.1657.

**Synthesis of *tert*-butyl 13-(3-azidopropyl)-1-(9*H*-fluoren-9-yl)-3,12-dioxo-2,7,10-trioxa-4,13-diazahexadecan-16-oate (3)**

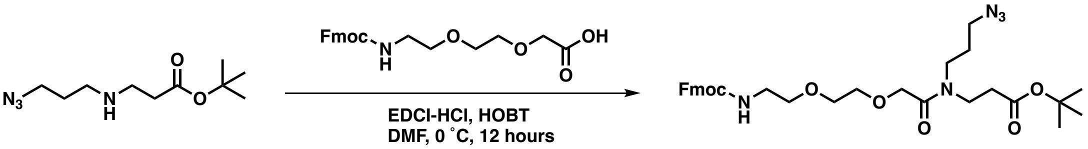


1-(9H-fluoren-9-yl)-3-oxo-2,7,10-trioxa-4-azadodecan-12-oic acid (1.0 g, 1.2 equiv.) and HOBT (0.45 g, 1.5 equiv.) were dissolved in 8.0 mL DMF under argon and stirred at 0 °C for 5 minutes. EDC·HCl (0.63 g, 1.5 equiv.) was then added and the reaction was stirred at 0 °C for 5 minutes. Next, compound (**2**) (0.50 g, 1.0 equiv.) and DIPEA (0.40 mL, 1.0 equiv.) were added, and the reaction mixture was stirred at 0 °C for 12 hours. Upon completion, the reaction was diluted with ethyl acetate, washed with brine, and dried with anhydrous $MgSO_4$. After concentrating in vacuum, the concentrated residue was purified by silica column chromatography (eluent gradient: 0% to 3% methanol in dichloromethane), yielding *tert-butyl 13-(3-azidopropyl)-1-(9H-fluoren-9-yl)-3,12-dioxo-2,7,10-trioxa-4,13-diazahexadecan-16-oate* (**3**) as a thick yellow liquid (1.1 g, 83%). Rf= 0.59 in 3% methanol in dichloromethane. $^1$H-NMR (400 MHz, CDCl3) δ 7.73 (d, *J* = 7.5 Hz, 2H), 7.59 (d, *J* = 7.5 Hz, 2H), 7.41 – 7.32 (m, 2H), 7.32 – 7.24 (m, 2H), 5.73 – 5.54 (m, 1H), 4.36 (s, 2H), 4.28 – 4.14 (m, 3H), 3.72 – 3.44 (m, 8H), 3.44 – 3.18 (m, 6H), 2.57 – 2.40 (m, 2H), 1.85 – 1.73 (m, 2H), 1.44 – 1.37 (m, 9H). $^{13}$C-NMR (101 MHz, $CDCl_3$) δ= 171.30, 170.46, 169.39, 169.17, 156.64, 144.08, 141.33, 127.69, 127.07, 125.19, 119.98, 81.60, 80.91, 70.71, 70.34, 70.21, 70.09, 66.66, 49.22, 48.54, 47.31, 44.90, 43.36, 34.88, 33.75, 28.09, 27.03. HRMS $C_{31}H_{41}N_5O_7$ $[M+H]^+$ calculated 596.3084, found 595.3006.

***Tert*-butyl 3-(2-(2-(2-aminoethoxy)ethoxy)-*N*-(3-azidopropyl)acetamido)propanoate (4)**

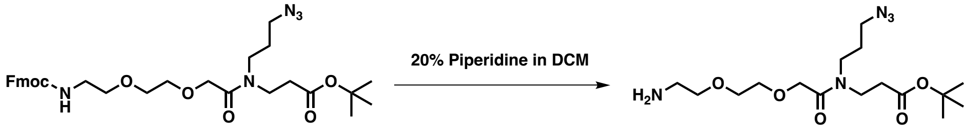


Compound (**3**) (2.3 g) was stirred in 20% piperidine/DCM (10 mL) for 1 hour at room temperature and concentrated. The reaction mixture was purified by silica column chromatography (eluent gradient: 0% to 10% methanol and 1% ammonia in dichloromethane), yielding *tert-butyl 3-(2-(2-(2-aminoethoxy)ethoxy)-N-(3-azidopropyl)acetamido)propanoate* **(4)** as a thick yellow liquid (1.2 g, 82%). Rf= 0.21 in 10% methanol and 1% ammonia in dichloromethane. $^1$H-NMR (400 MHz, CDCl3) δ= 4.30 (s, 2H), 3.86 (t, *J* = 5.0 Hz, 2H), 3.78 – 3.66 (m, 5H), 3.56 – 3.27 (m, 6H), 3.23 (t, *J* = 5.0 Hz, 2H), 2.51 (t, *J* = 7.0 Hz, 2H), 1.89 – 1.75 (m,

2H), 1.49 – 1.39 (m, 9H). $^{13}$C-NMR (101 MHz, $CDCl_3$) δ= 171.34, 170.40, 170.00, 169.78, 81.78, 81.14, 70.71, 70.14, 69.03, 67.25, 49.28, 48.60, 44.78, 43.61, 42.87, 42.42, 39.83, 34.70, 33.86, 28.42 – 27.84 (m), 27.06. HRMS $C_{16}H_{31}N_5O_5$ $[M+H]^+$ calculated 374.2403, found 374.2396.

**Synthesis of Das-AI**

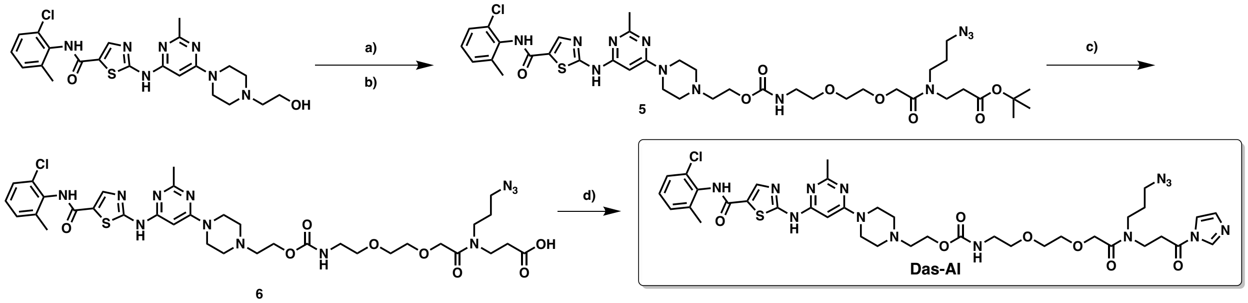


**Scheme 2 | Synthesis of Das-AI**. a) CDI, DMF, RT, 2.5 hours. b) Intermediate **4**, DMAP, DIPEA, DMF, 70 ˚C, 22 hours. c) 50% TFA in DCM, RT, 1 hour. d) CDI, DMSO, RT.

**Synthesis of *tert*-butyl 14-(3-azidopropyl)-1-(4-(6-((5-((2-chloro-6-methylphenyl)carbamoyl)thiazol-2-yl)amino)-2-methylpyrimidin-4-yl)piperazin-1-yl)-4,13-dioxo-3,8,11-trioxa-5,14-diazaheptadecan-17-oate (5)**

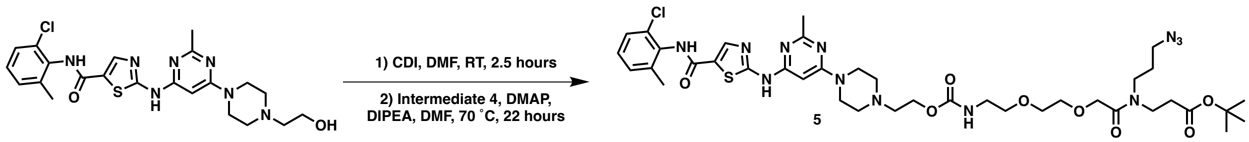


Dasatinib (0.30 g, 1.0 equiv.) and CDI (0.17 g, 1.7 equiv.) were dissolved in dry DMF (3 mL) and stirred at room temperature for 2 hours. Activation of dasatinib was monitored by NMR, and another portion of CDI (0.051 g) was added and stirred for 30 min at room temperature until the starting material was fully activated. Intermediate **4** (0.41 g, 1.8 equiv.), DMAP (0.0075 g, 0.1 equiv.), and DIPEA (0.159 mL, 2 equiv.) were added and stirred for 22 hours at 70°C. The reaction mixture was diluted with ethyl acetate, washed with brine, dried with anhydrous $MgSO_4$. Purification with silica column chromatography (eluent gradient: 0% to 20% methanol in dichloromethane) yielded *tert-butyl 14-(3-azidopropyl)-1-(4-(6-((5-((2-chloro-6-methylphenyl)carbamoyl)thiazol-2-yl)amino)-2-methylpyrimidin-4-yl)piperazin-1-yl)-4,13-dioxo-3,8,11-trioxa-5,14-diazaheptadecan-17-oate* **(5)** as a white solid (0.44 g, 80%). Rf=0.82 in 20% methanol in dichloromethane. $^1$H-NMR (400 MHz, $CD_3OD$) δ= 8.26 (s, 1H), 8.13 (s, 1H), 7.27 (d, $J$ = 7.2 Hz, 1H), 7.21 – 7.10 (m, 2H), 5.93 (s, 1H), 4.28 (s, 2H), 4.20 (s, 1H), 4.16 (t, $J$ = 5.6 Hz, 2H), 3.65 – 3.54 (m, 9H), 3.54 – 3.44 (m, 4H), 3.38 – 3.20 (m, 6H), 2.62 (t, $J$ = 5.6 Hz, 2H), 2.58 – 2.43 (m, 6H), 2.41 (s, 3H), 2.27 (s, 3H), 1.85 – 1.70 (m, 2H), 1.39 (s, 9H). $^{13}$C-NMR (101 MHz, $CD_3OD$) δ= 172.58, 172.19, 171.68, 171.43, 167.24, 165.09, 164.28, 163.12, 158.62, 158.36, 142.17, 140.24, 134.26, 134.11, 130.05, 129.42, 128.22, 126.72, 83.89, 82.24, 81.90, 79.55, 79.22, 78.90, 71.71, 71.66, 71.15, 70.91, 70.64, 70.52, 62.95, 58.01, 54.80,

53.97, 50.13, 45.85, 44.84, 44.19, 43.96, 43.22, 41.67, 35.35, 34.49, 28.82, 28.35, 27.88, 25.76, 18.85. HRMS $C_{39}H_{55}ClN_{12}O_8S$ $[M+H]^+$ calculated 887.3753, found 887.3762.

**Synthesis of 14-(3-azidopropyl)-1-(4-(6-((5-((2-chloro-6-methylphenyl)carbamoyl)thiazol-2-yl)amino)-2-methylpyrimidin-4-yl)piperazin-1-yl)-4,13-dioxo-3,8,11-trioxa-5,14-diazaheptadecan-17-oic acid (6)**

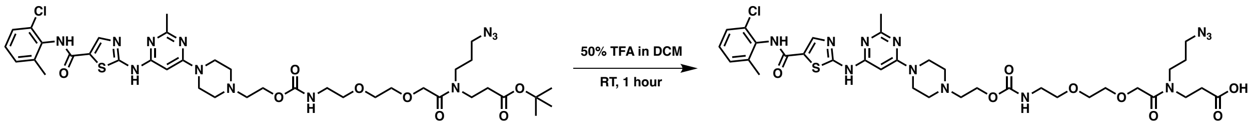


Compound **(5)** (216 mg) was dissolved in 50% TFA/DCM (4 mL) and stirred at room temperature for 3 h until full conversion of the starting material. After evaporating twice with toluene, the reaction mixture was purified on a short silica gel column (~2 cm) (eluent: 20% methanol in dichloromethane), yielding *14-(3-azidopropyl)-1-(4-(6-((5-((2-chloro-6-methylphenyl)carbamoyl)thiazol-2-yl)amino)-2-methylpyrimidin-4-yl) piperazin-1-yl)-4,13-dioxo-3,8,11-trioxa-5,14-diazaheptadecan-17-oic acid* **(6)** as a solid (192 mg, 95%). Rf= 0.39 in 20% methanol in dichloromethane. $^1$H-NMR (300 MHz, $CD_3OD$) δ= 8.97 (s, 1H), 8.20 (s, 1H), 7.35 – 7.24 (m, 1H), 7.24 – 7.12 (m, 2H), 6.63 (s, 1H), 4.40 (d, *J* = 5.8 Hz, 2H), 4.35 (s, 1H), 4.26 (s, 1H), 3.93 (s, 2H), 3.69 – 3.20 (m, 24H), 2.59 (t, *J* = 7.0 Hz, 2H), 2.53 (s, 3H), 2.27 (s, 3H), 1.88 – 1.70 (m, 2H). $^{13}$C-NMR (75 MHz, $CD_3OD$) δ= 175.33, 174.72, 171.99, 171.79, 165.28, 162.84, 162.11, 161.69, 157.97, 155.48, 144.76, 140.27, 139.76, 134.05, 130.24, 129.80, 128.37, 127.50, 119.34, 115.51, 85.23, 71.53, 71.18, 71.02, 70.33, 70.09, 59.68, 57.19, 52.77, 45.72, 44.22, 43.77, 43.38, 42.74, 41.70, 33.84, 33.15, 28.71, 27.81, 23.37, 18.72. Note: The compound is a mixture of rotamers. HRMS $C_{35}H_{47}ClN_{12}O_8S$ $[M+H]^+$ calculated 831.3127, found 831.3114.

**Synthesis of Das-AI**

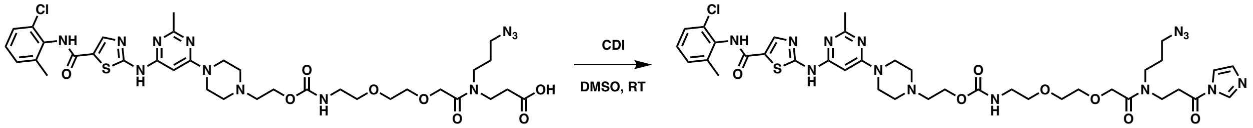


Compound (**6**) (45 mg) was dissolved in anhydrous d6-DMSO (54 µL) as a 1 M solution under argon. To it was added 54 µL of 1.4 M solution of 1,1´-carbonyldiimidazole (CDI). The reaction was stirred at room temperature for 2 hours at room temperature and used as a 500 mM stock solution for biological experiments without further purification. The resulting solution of 2-(4-(6-((5-((2-chloro-6-methylphenyl)carbamoyl)thiazol-2-yl)amino)-2-methylpyrimidin-4-yl)piperazin-1-yl)ethyl (2-(2-(2-((3-(1H-imidazol-1-yl)-3-oxopropyl)(3-azidopropyl) amino)-2-oxoethoxy)ethoxy)ethyl)carbamate (Das-AI) is colorless. 1H-NMR (300 MHz, DMSO) δ= 8.27 (s, 1H), 8.25 (s, 1H), 7.59 (d, J = 1.6 Hz, 1H), 7.38 (d, J = 7.2 Hz, 1H), 7.32 – 7.18 (m, 3H), 7.07 (s, 1H), 6.14 (s, 1H), 4.27 – 4.11 (m, 4H), 3.74 – 3.05 (m, 20H), 2.84 – 2.74 (m, 6H), 2.54 – 2.46 (m, 2H), 2.41 (s, 3H), 2.24 (s, 3H), 1.86 – 1.70 (m, 2H). $^{13}$C-NMR (75 MHz,

DMSO) δ= 165.29, 162.51, 162.28, 160.02, 158.89, 158.48, 157.07, 140.94, 138.89, 137.27, 133.48, 132.52, 130.32, 129.05, 128.17, 127.03, 125.80, 117.56, 115.24, 82.94, 69.83, 69.43, 69.13, 60.35, 56.09, 52.04, 48.61, 48.26, 42.73, 42.29, 25.59, 18.32. HRMS $C_{38}H_{49}ClN_{14}O_7S$ $[M+H]^+$ calculated 881.3396, found 881.3393.

**Synthesis of HCQ-AI**

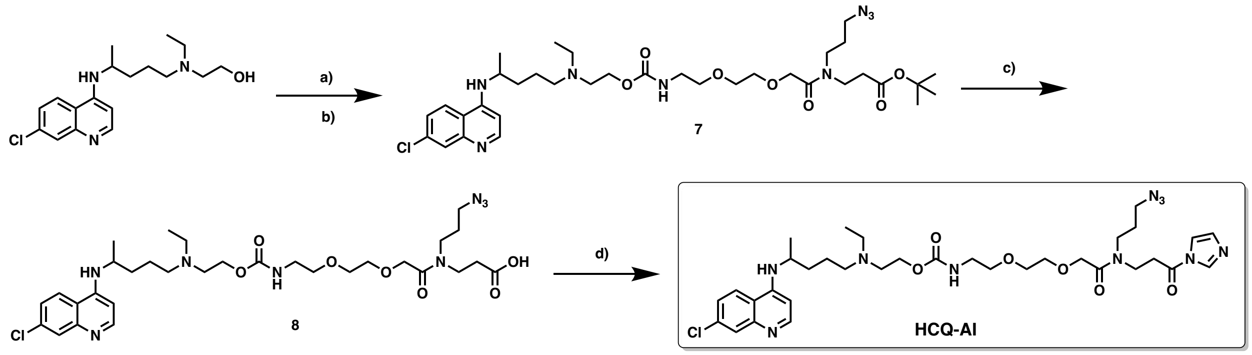


**Scheme 3 | Synthesis of HCQ-AI**. a) CDI, DMF, under argon, 2.5 hours, RT. b) Intermediate **4**, DMAP, DIPEA, DMF, RT, overnight. c) 50% TFA in DCM, RT, 2.5 hours.

**Synthesis of *tert*-butyl 4-(3-azidopropyl)-22-((7-chloroquinolin-4-yl)amino)-18-ethyl-5,14-dioxo-7,10,15-trioxa-4,13,18-triazatricosanoate (7)**

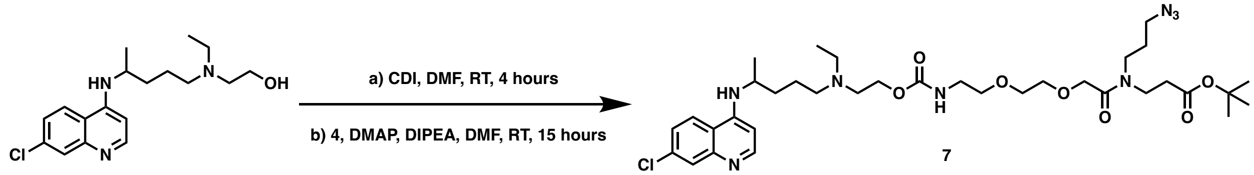


Hydroxychloroquine (300 mg, 1.0 equiv.) was dissolved in 3 mL of anhydrous DMF. CDI (590 mg, 4.0 equiv.) was added to the solution in one portion under argon. The resulting reaction mixture was stirred at room temperature for 2.5 hours before quenching by 3 mL of water on ice. The reaction was then diluted with 30 mL of ethyl acetate, washed with water (15 mL x 2) and brine (15 mL x 1), and dried over anhydrous $Na_2SO_4$. The solution was concentrated in vacuum. The residue was redissolved in 1.5 mL of anhydrous DMF. To it was added *tert*-butyl 3-(2-(2-(2-aminoethoxy)ethoxy)-*N*-(3-azidopropyl)acetamido)propanoate (**4**) (410 mg, 2.0 equiv.), DMAP (11 mg, 0.1 equiv.) and DIPEA (290 mg, 2.5 equiv.). The resulting solution was stirred at room temperature overnight under argon. The reaction was diluted with ethyl acetate, washed with brine, and dried over anhydrous $Na_2SO_4$. Purification on silica gel (eluent gradient: 0% to 20% methanol in dichloromethane) afforded *tert*-butyl 4-(3-azidopropyl)-22-((7-chloroquinolin-4-yl)amino)-18-ethyl-5,14-dioxo-7,10,15-trioxa-4,13,18-triazatricosanoate (**7**) as a pale yellow solid (115 mg, 22%). Rf= 0.33 in 20% methanol in dichloromethane. $^1$H-NMR (400 MHz, $CDCl_3$) δ= 8.31 (d, *J* = 9.1 Hz, 1H), 8.23 (d, *J* = 6.1 Hz, 1H), 7.87 (s, 1H), 7.24 (d, *J* = 8.8 Hz, 1H), 7.12 (s, 1H), 6.38 (d, *J* = 6.3 Hz, 1H), 5.59 (t, *J* = 6.0 Hz, 1H), 4.16 (s, 1H), 4.09 (s, 1H), 4.03 (t, *J* = 6.1 Hz, 2H), 3.69 (p, *J* = 6.6 Hz, 1H), 3.58 – 3.14 (m, 14H), 2.62 (t, *J*

= 6.1 Hz, 2H), 2.55 – 2.36 (m, 6H), 1.86 – 1.43 (m, 6H), 1.32 (d, *J* = 4.8 Hz, 9H), 1.26 (d, *J* = 6.3 Hz, 3H), 0.91 (t, *J* = 7.1 Hz, 3H). $^{13}$C-NMR (101 MHz, $CDCl_3$) δ= 171.00, 170.23, 169.33, 169.09, 156.45, 151.93, 147.16, 144.57, 136.64, 125.79, 124.25, 123.88, 116.61, 98.43, 81.41, 80.77, 70.52, 70.01, 69.88, 62.06, 53.36, 51.78, 49.31, 49.05, 48.39, 48.14, 44.71, 43.17, 42.89, 42.00, 40.64, 34.67, 33.57, 33.49, 27.92, 26.88, 23.61, 19.86, 11.11. HRMS $C_{35}H_{55}ClN_8O_7$ $[M+H]^+$ calculated 735.3960, found 735.3948.

**Synthesis of 4-(3-azidopropyl)-22-((7-chloroquinolin-4-yl)amino)-18-ethyl-5,14-dioxo-7,10,15-trioxa-4,13,18-triazatricosanoic acid (8)**

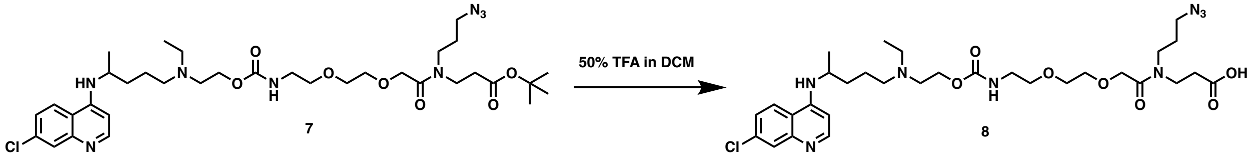


Tert-butyl 4-(3-azidopropyl)-22-((7-chloroquinolin-4-yl)amino)-18-ethyl-5,14-dioxo-7,10,15-trioxa-4,13,18-triazatricosanoate (**7**) (77 mg, 1.0 equiv.) was dissolved in 1 mL of DCM. The solution as cooled to 0 ˚C on ice and TFA (1 mL) was added. The resulting solution was stirred and allowed to warm to room temperature over two hours. The solvent was removed in vacuum. Purification of the residue by chromatography on a short (~2 cm) silica column (eluent: 20% methanol in dichloromethane) afforded 4-(3-azidopropyl)-22-((7-chloroquinolin-4-yl)amino)-18-ethyl-5,14-dioxo-7,10,15-trioxa-4,13,18-triazatricosanoic acid as a pale-yellow solid (42 mg, 60%). Rf= 0.37 in 20% methanol in dichloromethane. $^{1}$H-NMR (400 MHz, $CD_3OD$) δ= 8.50 (d, *J* = 9.1 Hz, 1H), 8.36 (d, *J* = 7.1 Hz, 1H), 7.87 (s, 1H), 7.63 (d, *J* = 9.1 Hz, 1H), 6.90 (d, *J* = 7.9 Hz, 1H), 4.49 – 4.02 (m, 5H), 3.68 – 3.16 (m, 21H), 2.68 – 2.50 (m, 2H), 1.82 (p, *J* = 14.5 Hz, 6H), 1.38 (d, *J* = 6.4 Hz, 3H), 1.28 (t, *J* = 7.2 Hz, 4H). $^{13}$C-NMR (75 MHz, $CD_3OD$) δ= 175.22, 174.63, 171.95, 171.73, 158.02, 157.09, 143.91, 140.90, 140.15, 128.49, 126.36, 120.31, 116.97, 99.89, 71.52, 71.14, 71.00, 70.36, 70.14, 59.97, 53.97, 52.69, 51.12, 50.20, 49.97, 45.75, 44.19, 43.78, 43.38, 41.64, 33.85, 33.41, 33.12, 28.75, 27.85, 21.87, 19.90, 8.99. HRMS $C_{31}H_{47}ClN_8O_7$ $[M+H]^+$ calculated 679.3334, found 679.3326.

**Synthesis of HCQ-AI**

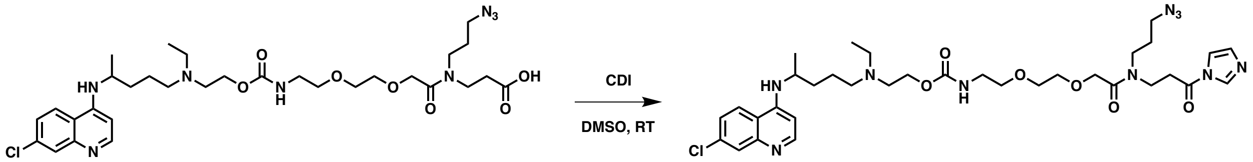


Compound (**8**) (48 mg) was dissolved in anhydrous d6-DMSO (71 µL) as a 1 M solution under argon. To it was added 71 µL of 1.4 M solution of 1,1´-carbonyldiimidazole (CDI). The reaction was stirred at room temperature for 2 hours at room temperature and used as a 500 mM stock solution for biological experiments without further purification. The resulting solution of 2-((4-((7-chloroquinolin-4-yl)amino)pentyl)(ethyl)amino)ethyl (2-(2-(2-((3-(1H-imidazol-1-yl)-3-oxopropyl)(3-azidopropyl)amino)-2-oxoethoxy)ethoxy)ethyl)carbamate (HCQ-AI) is colorless. $^{1}$H-NMR (400 MHz, DMSO) δ= 8.61 (d, *J* = 9.0 Hz, 1H), 8.48 (d, *J* = 6.4 Hz, 1H), 8.25 (s, 1H), 7.93 (d, *J* = 2.2 Hz, 1H), 7.64 – 7.56 (m, 2H), 7.10 – 7.04

(m, 1H), 6.77 (d, *J* = 6.5 Hz, 1H), 4.26 – 4.11 (m, 4H), 4.00 – 3.91 (m, 1H), 3.66 – 3.03 (m, 14H), 3.03 – 2.87 (m, 4H), 2.59 – 2.40 (m, 4H), 1.87 – 1.53 (m, 6H), 1.28 (d, *J* = 6.3 Hz, 3H), 1.10 (t, *J* = 7.1 Hz, 3H). $^{13}$C-NMR (101 MHz, DMSO) δ= 173.02, 172.87, 169.02, 168.89, 158.83, 158.51, 155.83, 155.77, 152.70, 149.02, 146.65, 143.05, 137.30, 136.21, 130.34, 125.52, 125.45, 122.52, 118.74, 117.58, 116.33, 115.77, 98.80, 69.84, 69.44, 69.17, 69.13, 59.54, 52.29, 50.74, 48.63, 48.27, 47.66, 44.22, 42.37, 42.28, 40.07, 33.29, 32.38, 32.30, 27.57, 26.54, 21.17, 19.58, 9.46. HRMS $C_{34}H_{49}ClN_{10}O_6$ $[M+H]^+$ calculated 729.3603, found 729.3594.

**Synthesis of Lev-AI**

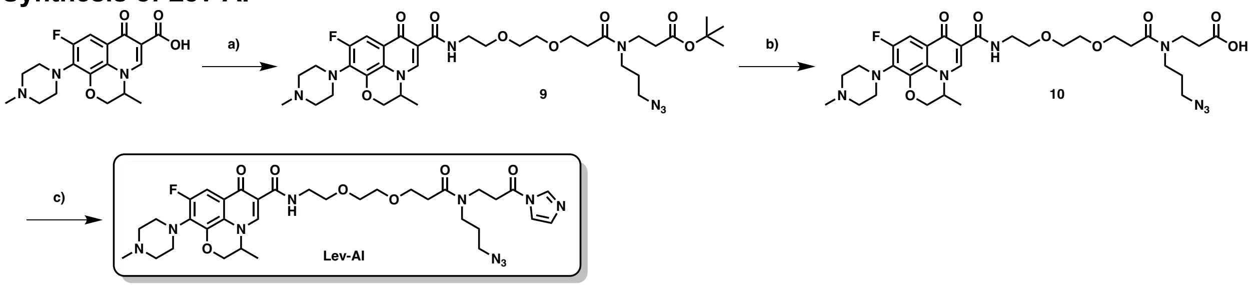

**Scheme 4 | Synthesis of Lev-AI**. a) Intermediate **4**, EDCI-HCl, HOBT, DMF, 0 ˚C to RT, overnight; b) 50% TFA in DCM, RT, 2.5 hours. c) CDI, DMSO, RT.

**Synthesis of *tert*-butyl 12-(3-azidopropyl)-1-(9-fluoro-3-methyl-10-(4-methylpiperazin-1-yl)-7-oxo-2,3-dihydro-7*H*-[1,4]oxazino[2,3,4-*ij*]quinolin-6-yl)-1,11-dioxo-5,8-dioxa-2,12-diazapentadecan-15-oate (9)**

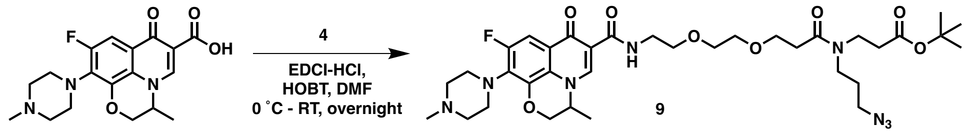

Levofloxacin (237 mg, 1.2 equiv.) was dissolved in 5 mL of DCM. The solution was cooled to 0 ˚C, and to it was added *tert*-butyl 3-(2-(2-(2-aminoethoxy)ethoxy)-*N*-(3-azidopropyl)acetamido)propanoate (**4**) (212 mg, 1.0 equiv.), HOBT (136 mg, 1.3 equiv.), EDCI-HCl (112 mg, 1.5 quiv.), and diisopropylethylamine (106 mg, 1.5 equiv.). The reaction was stirred and allowed to warm to room temperature overnight under argon. The reaction mixture was then diluted with ethyl acetate (80 mL) and washed with $NaHCO_3$, brine, and dried over anhydrous $Na_2SO_4$. Purification by chromatography on silica gel (eluent gradient: 0% to 20% methanol in dichloromethane) afforded *tert*-butyl 12-(3-azidopropyl)-1-(9-fluoro-3-methyl-10-(4-methylpiperazin-1-yl)-7-oxo-2,3-dihydro-7*H*-[1,4]oxazino[2,3,4-*ij*]quinolin-6-yl)-1,11-dioxo-5,8-dioxa-2,12-diazapentadecan-15-oate (**9**) as an off-white solid (257 mg, 65%). Rf= 0.59 in 20% methanol in dichloromethane. $^1$H-NMR (400 MHz, $CDCl_3$) δ= 9.97 (d, *J* = 5.4 Hz, 1H), 8.43 (d, *J* = 1.2 Hz, 0H), 7.41 (d, *J* = 12.5 Hz, 0H), 4.39 – 4.03 (m, 4H), 3.65 – 2.90 (m, 22H), 2.72 (t, *J* = 5.2 Hz, 1H), 2.40 – 2.28 (m, 4H), 2.15 (s, 3H), 2.04 (s, 3H), 1.70 – 1.56 (m, 2H), 1.37 (d, *J* = 6.8 Hz, 3H), 1.23 (d, *J* = 3.0 Hz, 9H) (Note:

Compound 9 is mixture of isomers). $^{13}C$-NMR (101 MHz, $CDCl_3$) δ= 174.77, 170.80, 170.24 – 169.79, 169.18, 169.07, 168.95, 168.82, 164.77, 156.59, 156.21, 154.13, 143.64, 139.23, 131.41, 124.00, 121.97, 110.54, 104.43, 81.16, 81.03, 80.89, 80.66 – 80.21, 71.88, 71.05, 70.28, 70.10, 69.88, 69.93 – 69.45, 69.01, 68.91, 67.91, 55.37, 54.49, 50.23, 48.85, 48.43 – 47.96, 46.08, 45.15, 44.75 – 44.35, 44.27, 44.11, 43.07 – 42.55, 42.42, 41.75, 41.05, 38.72, 36.81, 34.59, 34.48, 34.35, 33.33, 27.70, 26.67, 17.94, 14.08 (Note: Compound 9 is mixture of isomers). HRMS $C_{34}H_{49}FN_8O_8$ $[M+H]^+$ calculated 717.3735, found 717.3719.

**Synthesis of 12-(3-azidopropyl)-1-(9-fluoro-3-methyl-10-(4-methylpiperazin-1-yl)-7-oxo-2,3-dihydro-7*H*-[1,4]oxazino[2,3,4-*ij*]quinolin-6-yl)-1,11-dioxo-5,8-dioxa-2,12-diazapentadecan-15-oic acid (10)**

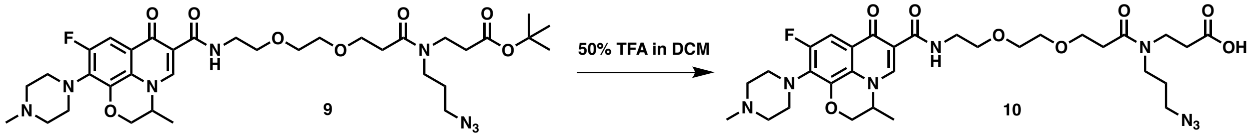


Tert-butyl 12-(3-azidopropyl)-1-(9-fluoro-3-methyl-10-(4-methylpiperazin-1-yl)-7-oxo-2,3-dihydro-7H-[1,4]oxazino[2,3,4-ij]quinolin-6-yl)-1,11-dioxo-5,8-dioxa-2,12-diazapentadecan-15-oate (**9**) (200 mg, 1.0 equiv.) was dissolved in 4 mL of 50% TFA in DCM at room temperature. The resulting solution was stirred at room temperature for 3 hours. The solvent was removed in vacuum and purification by a short (~2 cm) silica gel column (eluent: 20% methanol in dichloromethane) afforded (*S*)-11-(3-azidopropyl)-1-(9-fluoro-3-methyl-10-(4-methylpiperazin-1-yl)-7-oxo-2,3-dihydro-7*H*-[1,4]oxazino[2,3,4-*ij*]quinolin-6-yl)-1,10-dioxo-5,8-dioxa-2,11-diazatetradecan-14-oic acid (**10**) as an off-white solid (177 mg, 96%). Rf=0.25 in 20% methanol in dichloromethane. $^1H$ NMR (500 MHz, $CD_3OD$) δ= 8.65 (s, 1H), 7.47 (s, 1H), 4.63 (s, 1H), 4.53 – 4.24 (m, 6H), 3.82 – 3.05 (m, 20H), 2.95 (s, 3H), 2.65 – 2.48 (m, 3H), 1.87 – 1.73 (m, 2H), 1.49 (d, $J$ = 6.8 Hz, 3H). $^{13}C$-NMR (126 MHz, $CD_3OD$) δ= 176.19, 175.26, 175.08, 174.65, 174.52, 171.86, 171.65, 162.63, 162.35, 157.85, 156.78, 155.89, 146.09, 142.20, 130.83, 125.70, 124.48, 121.40, 119.08, 116.76, 114.44, 111.49, 104.92, 71.65, 71.37, 71.22, 71.08, 70.99, 70.41, 70.24, 70.10, 69.96, 69.7, 67.87, 56.54, 55.45, 50.19, 49.62, 45.66, 44.17, 43.93, 43.69, 43.52, 43.35, 40.45, 40.00, 33.83, 33.11, 28.71, 27.83, 18.18. HRMS $C_{30}H_{41}FN_8O_8$ $[M+H]^+$ calculated 661.3109, found 661.3098.

**Synthesis of Lev-AI**

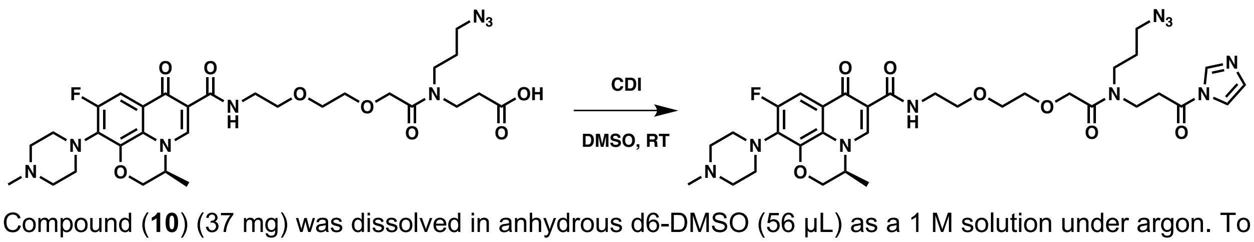


Compound (**10**) (37 mg) was dissolved in anhydrous d6-DMSO (56 µL) as a 1 M solution under argon. To it was added 56 µL of 1.4 M solution of 1,1´-carbonyldiimidazole (CDI). The reaction was stirred at room temperature for 2 hours at room temperature and used as a 500 mM stock solution for biological experiments without further purification. The resulting solution of (*S*)-*N*-(2-(2-(2-((3-(1*H*-imidazol-1-yl)-3-

oxopropyl)(3-azidopropyl)amino)-2-oxoethoxy)ethoxy)ethyl)-9-fluoro-3-methyl-10-(4-methylpiperazin-1-yl)-7-oxo-2,3-dihydro-7*H*-[1,4]oxazino[2,3,4-*ij*]quinoline-6-carboxamide (**Lev-AI**) is colorless. $^{1}$H-NMR (400 MHz, DMSO) δ= 10.02 (t, *J* = 5.5 Hz, 1H), 8.79 (s, 1H), 8.27 (s, 1H), 7.71 (d, *J* = 1.6 Hz, 1H), 7.55 (d, *J* = 12.4 Hz, 1H), 7.01 (d, *J* = 1.4 Hz, 1H), 4.85 (d, *J* = 7.0 Hz, 1H), 4.58 – 4.09 (m, 6H), 3.68 – 2.21 (m, 25H), 1.81 – 1.65 (m, 2H), 1.42 (d, *J* = 6.7 Hz, 3H). $^{13}$C-NMR (75 MHz, DMSO) δ= 174.12, 172.84, 168.86, 164.16, 158.80, 145.05, 136.02, 130.31, 129.55, 124.33, 122.50, 119.22, 116.64, 110.24, 103.51, 69.82, 69.37, 69.08, 68.57, 68.31, 54.05, 53.87, 48.61, 48.19, 44.19, 43.48, 42.32, 41.58, 33.28, 27.56, 26.52, 17.89. (Note: Lev-AI is mixture of isomers) HRMS $C_{33}H_{43}FN_{10}O_7$ $[M+H]^+$ calculated 711.3378, found 711.3362.

**Synthesis of Lev-diazirine**

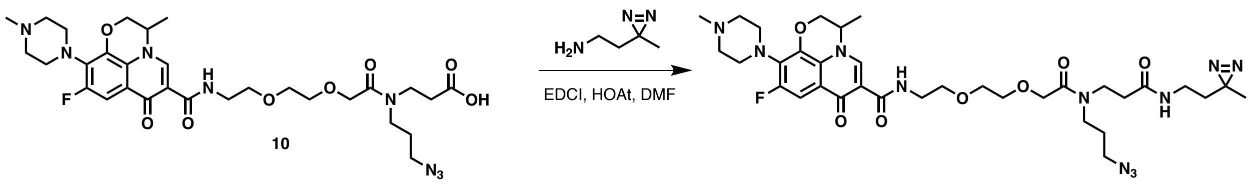


Compound (**10**) (40 mg, 1.2 equiv.) was dissolved in 2 mL of DCM. The solution was cooled to 0 ˚C, and to it was added 2-(3-methyl-3*H*-diazirin-3-yl)ethan-1-amine (5 mg, 1.0 equiv.), HOBT (10 mg, 1.t equiv.), EDCI-HCl (13 mg, 1.3 quiv.), and diisopropylethylamine (10 mg, 1.5 equiv.). The reaction was stirred and allowed to warm to room temperature overnight under argon. The reaction mixture was then diluted with ethyl acetate (30 mL) and washed with $NaHCO_3$, brine, and dried over anhydrous $Na_2SO_4$. Purification by chromatography on silica gel (eluent gradient: 0% to 20% methanol in dichloromethane) afforded *N*-(9-(3-azidopropyl)-15-(3-methyl-3*H*-diazirin-3-yl)-8,12-dioxo-3,6-dioxa-9,13-diazapentadecyl)-9-fluoro-3-methyl-10-(4-methylpiperazin-1-yl)-7-oxo-2,3-dihydro-7*H*-[1,4]oxazino[2,3,4-*ij*]quinoline-6-carboxamide (**Lev-diazirine**) as a brown solid (13 mg, 29%). $^{1}$H-NMR (500 MHz, DMSO) δ= 10.06 – 10.00 (m, 1H), 8.77 (s, 1H), 8.04 – 7.93 (m, 1H), 7.52 (dd, *J* = 12.6, 2.3 Hz, 1H), 4.87 – 4.79 (m, 1H), 4.56 – 4.12 (m, 4H), 3.64 – 2.13 (m, 29H), 1.81 – 1.64 (m, 2H), 1.47 – 1.38 (m, 5H), 1.00 (s, 3H). $^{13}$C-NMR (126 MHz, DMSO) δ= 174.11, 170.26, 168.70, 164.16, 156.15, 154.20, 144.89, 140.17, 130.74, 124.32, 121.87, 110.01, 103.33, 69.78, 69.58, 69.33, 69.24, 69.07, 68.09, 55.24, 53.96, 50.01, 48.59, 48.23, 45.93, 44.19, 42.71, 42.18, 42.06, 38.47, 34.73, 33.82, 33.64, 27.54, 26.43, 24.76, 19.27, 17.88. (Note: Lev-diazirine is mixture of isomers) Rf= 0.62 in 20% methanol in dichloromethane. HRMS $C_{34}H_{48}FN_{11}O_7$ $[M+H]^+$ calculated 742.3800, found 742.3788.

**Synthesis of *tert*-butyl 12-(3-azidopropyl)-2,11-dioxo-6,9-dioxa-3,12-diazapentadecan-15-oate (11)**

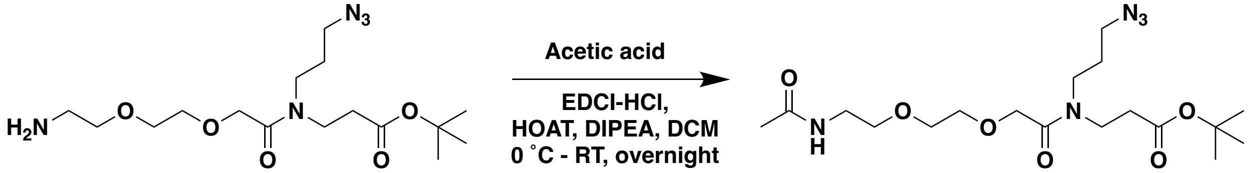

*Tert*-butyl 3-(2-(2-(2-aminoethoxy)ethoxy)-*N*-(3-azidopropyl)acetamido)propanoate (171 mg, 1.0 equiv.) was dissolved in 4 mL of DCM. The solution was cooled to 0 ˚C, and to it was added acetic acid (32 mg, 1.2 equiv.), HOAT (90 mg, 1.5 equiv.), EDCI-HCl (110 mg, 1.3 quiv.), and diisopropylethylamine (86 mg, 1.5 equiv.). The reaction was stirred and allowed to warm to room temperature overnight under argon. The reaction mixture was then diluted with ethyl acetate (30 mL) and washed with $NaHCO_3$, brine, and dried over anhydrous $Na_2SO_4$. Purification by chromatography on silica gel (eluent: 20% methanol in dichloromethane) afforded *tert*-butyl 12-(3-azidopropyl)-2,11-dioxo-6,9-dioxa-3,12-diazapentadecan-15-oate (**11**) as a yellow oil (155 mg, 84%). Rf= 0.38 in 10% methanol in dichloromethane. $^{1}$H-NMR (500 MHz, $CDCl_3$) δ 6.69 (s, 1H), 4.28 (s, 1H), 4.23 (s, 1H), 3.72 – 3.28 (m, 14H), 2.54 (t, *J* = 7.1 Hz, 2H), 2.00 (s, 2H), 1.85 (s, 3H), 1.45 (s, 9H). Note: This compound is a mixture of rotamers. $^{13}$C-NMR (126 MHz, $CDCl_3$) δ= 171.10, 170.29, 169.29, 80.89, 70.75, 69.97, 69.83, 69.57, 48.98, 44.23, 41.95, 39.24, 34.73, 33.64, 27.96, 26.90, 23.00. HRMS $C_{18}H_{33}N_5O_6$ $[M+H]^+$ calculated 416.2509, found 416.2502.

**Synthesis of 12-(3-azidopropyl)-2,11-dioxo-6,9-dioxa-3,12-diazapentadecan-15-oic acid (12)**

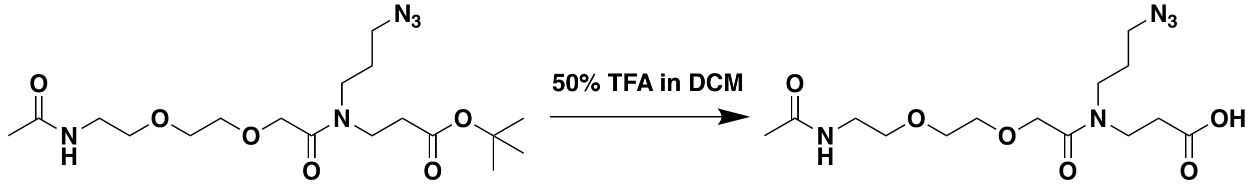


*Tert*-butyl 12-(3-azidopropyl)-2,11-dioxo-6,9-dioxa-3,12-diazapentadecan-15-oate (**11**) (150 mg, 1.0 equiv.) was dissolved in 4 mL of 50% TFA in DCM at room temperature. The resulting solution was stirred at room temperature for 1 hours. The solvent was removed in vacuum and purification by a short (~2 cm) silica gel column (eluent: 10% methanol in dichloromethane) afforded 12-(3-azidopropyl)-2,11-dioxo-6,9-dioxa-3,12-diazapentadecan-15-oic acid (**12**) as a yellow oil (104 mg, 81%). Rf= 0.54 in 10% methanol in dichloromethane. $^{1}$H-NMR (300 MHz, DMSO) δ= 10.81 (s, 1H), 7.88 (s, 1H), 4.20 (s, 1H), 4.14 (s, 1H), 3.63 – 3.07 (m, 14H), 2.61 – 2.38 (m, 2H), 1.80 (d, *J* = 1.3 Hz, 3H), 1.77 – 1.64 (m, 2H). $^{13}$C-NMR (75 MHz, DMSO) δ= 172.75, 169.24, 168.76, 69.78, 69.38, 69.12, 48.57, 48.22, 44.17, 42.24, 41.50, 38.56, 32.18, 26.49, 22.53. HRMS $C_{14}H_{25}N_5O_6$ $[M+H]^+$ calculated 360.1883, found 360.1877.

**Synthesis of Linker-AI**

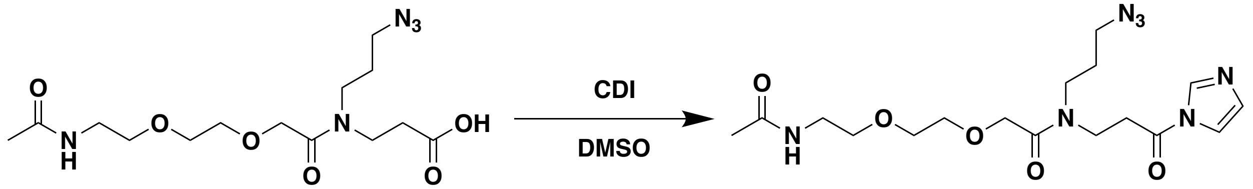


Compound (**12**) (45 mg) was dissolved in anhydrous d6-DMSO (125 µL) as a 1 M solution under argon. To it was added 125 µL of 1.4 M solution of 1,1´-carbonyldiimidazole (CDI). The reaction was stirred at room temperature for 2 hours at room temperature and used as a 500 mM stock solution for biological experiments without further purification. The resulting solution of *N*-(3-(1*H*-imidazol-1-yl)-3-oxopropyl)-2-(2-(2-acetamidoethoxy)ethoxy)-*N*-(3-azidopropyl)acetamide (**Linker-AI**) is colorless. $^{1}$H-NMR (300 MHz, DMSO) δ= 8.47 (s, 1H), 8.43 (s, 1H), 7.94 (s, 1H), 7.72 (d, *J* = 1.6 Hz, 1H), 7.07 (d, *J* = 4.3 Hz, 3H), 4.25 (s, 1H), 4.16 (s, 1H), 3.77 – 3.02 (m, 14H), 2.65 – 2.34 (m, 2H), 1.86 – 1.66 (m, 3H). $^{13}$C-NMR (75 MHz, DMSO) δ 169.39, 169.20, 169.02, 137.15, 130.34, 116.49, 69.88, 69.42, 69.14, 48.63, 48.28, 44.27, 42.38, 41.35, 41.00, 38.61, 33.98, 33.32, 27.53, 26.43, 22.55. HRMS $C_{17}H_{27}N_7O_5$ $[M+H]^+$ calculated 410.2152, found 410.2147

$^{1}H$-NMR spectrum of **1**

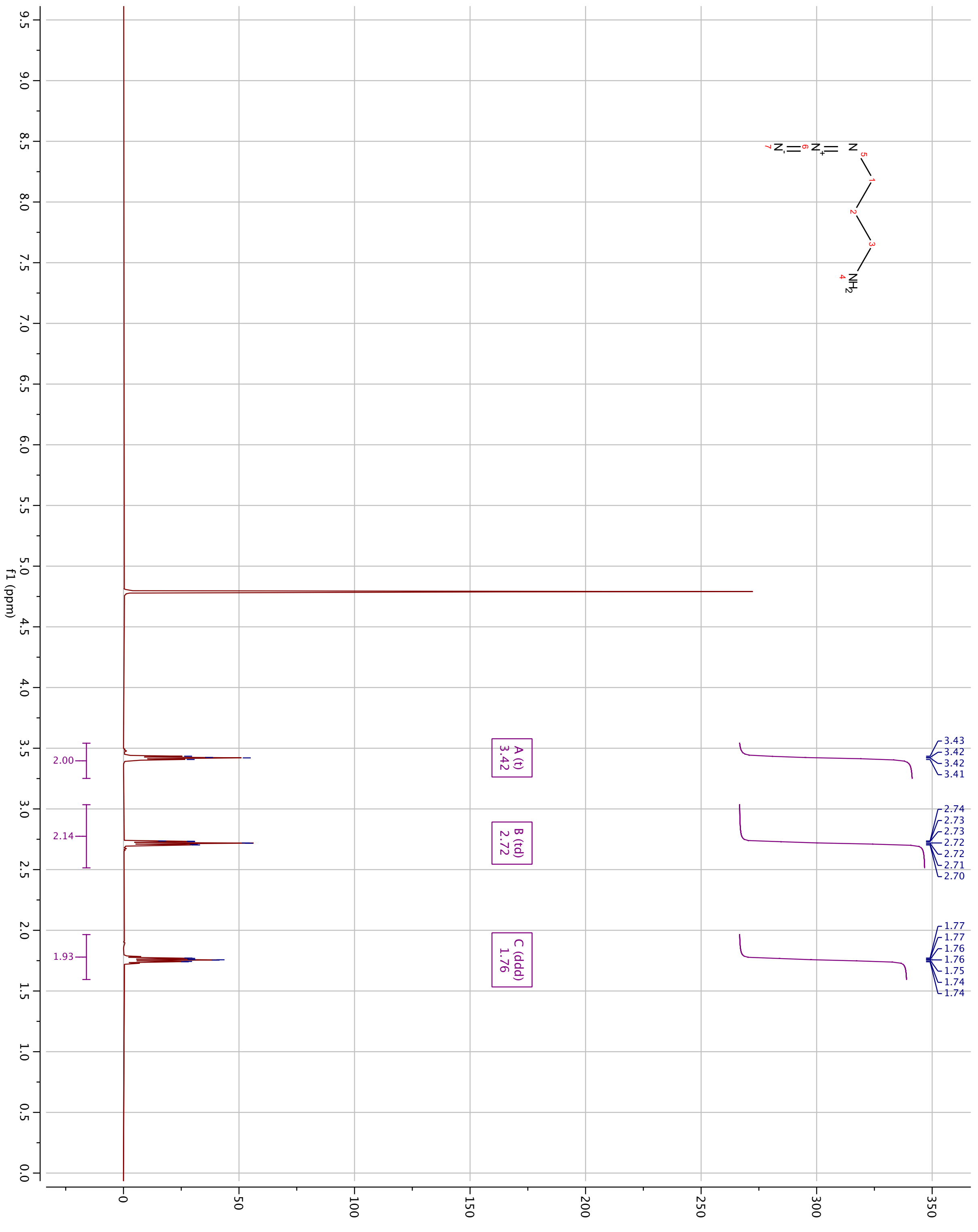

$^{1}$H-NMR spectrum of **2**

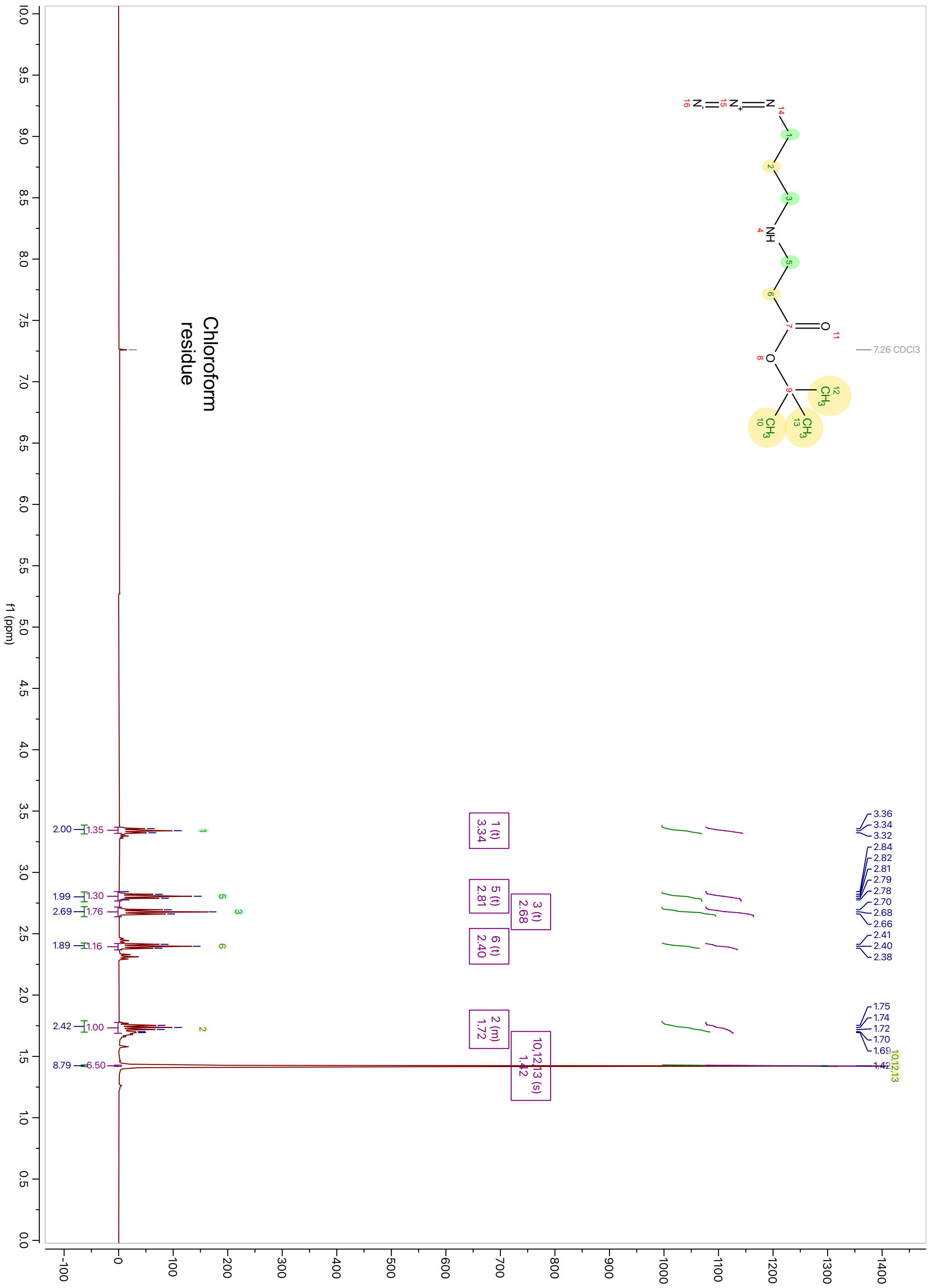

$^{13}$C-NMR spectrum of **2**

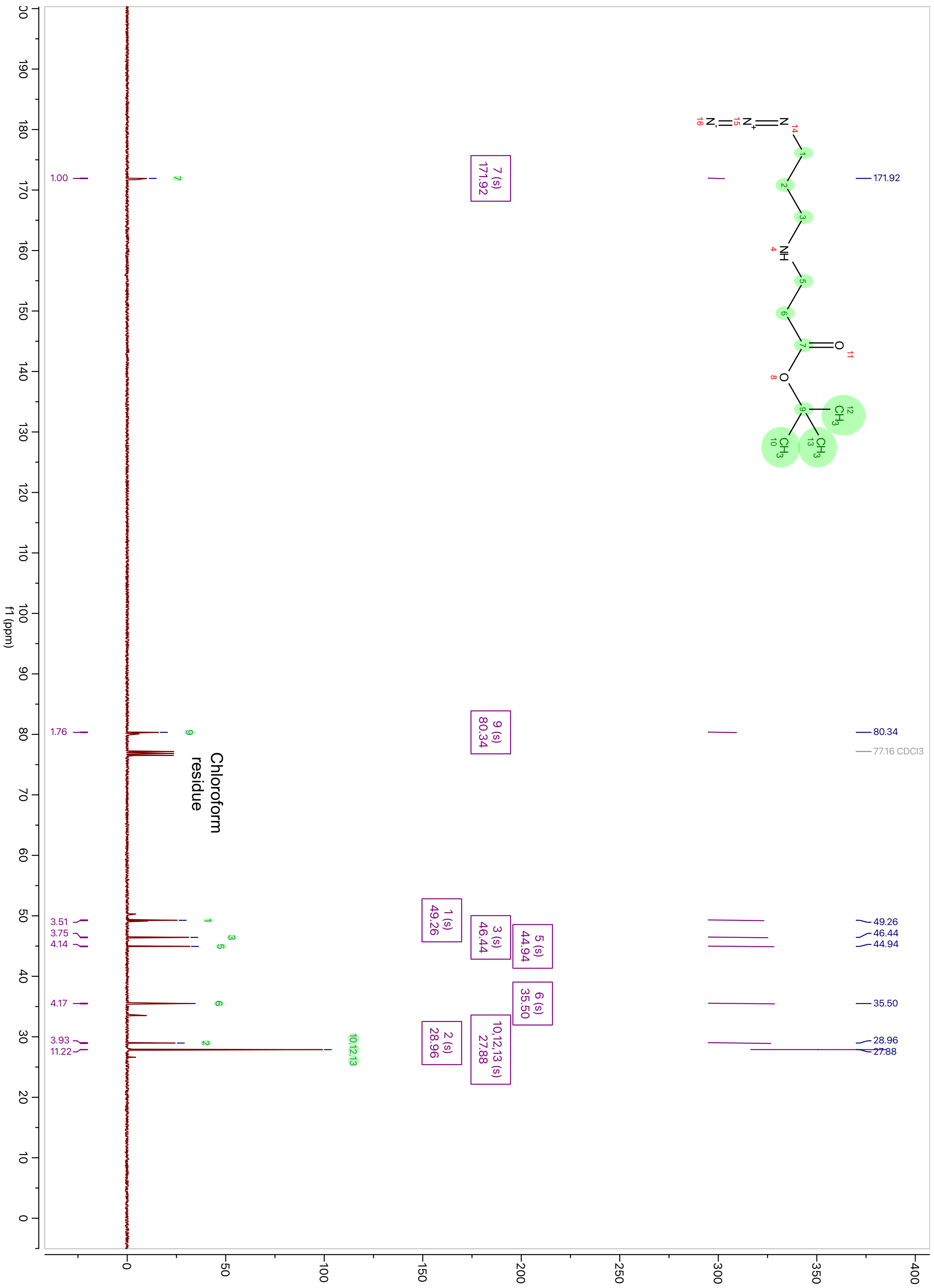

[1]H-NMR spectrum of **3**

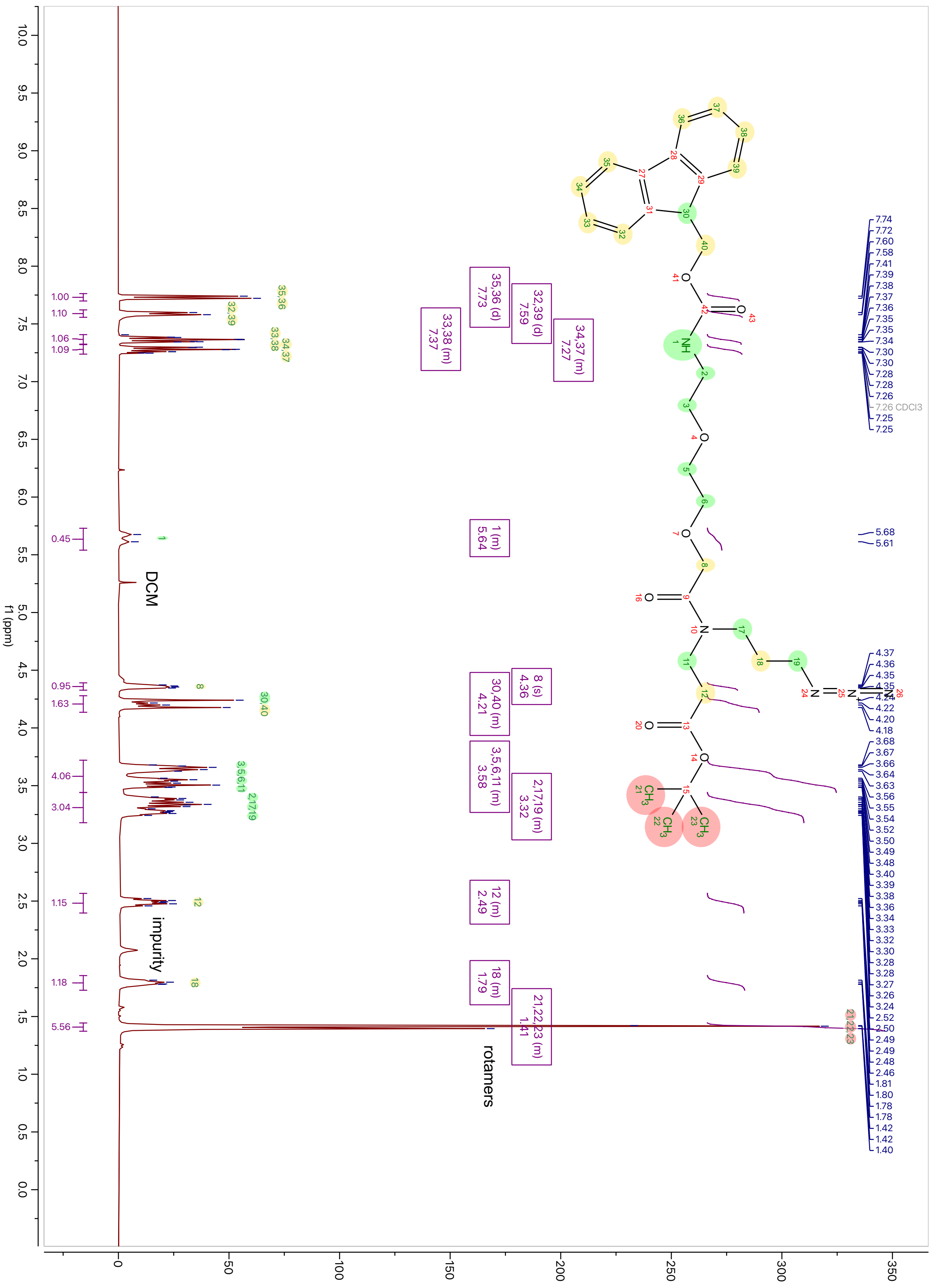

$^{13}$C-NMR spectrum of **3**

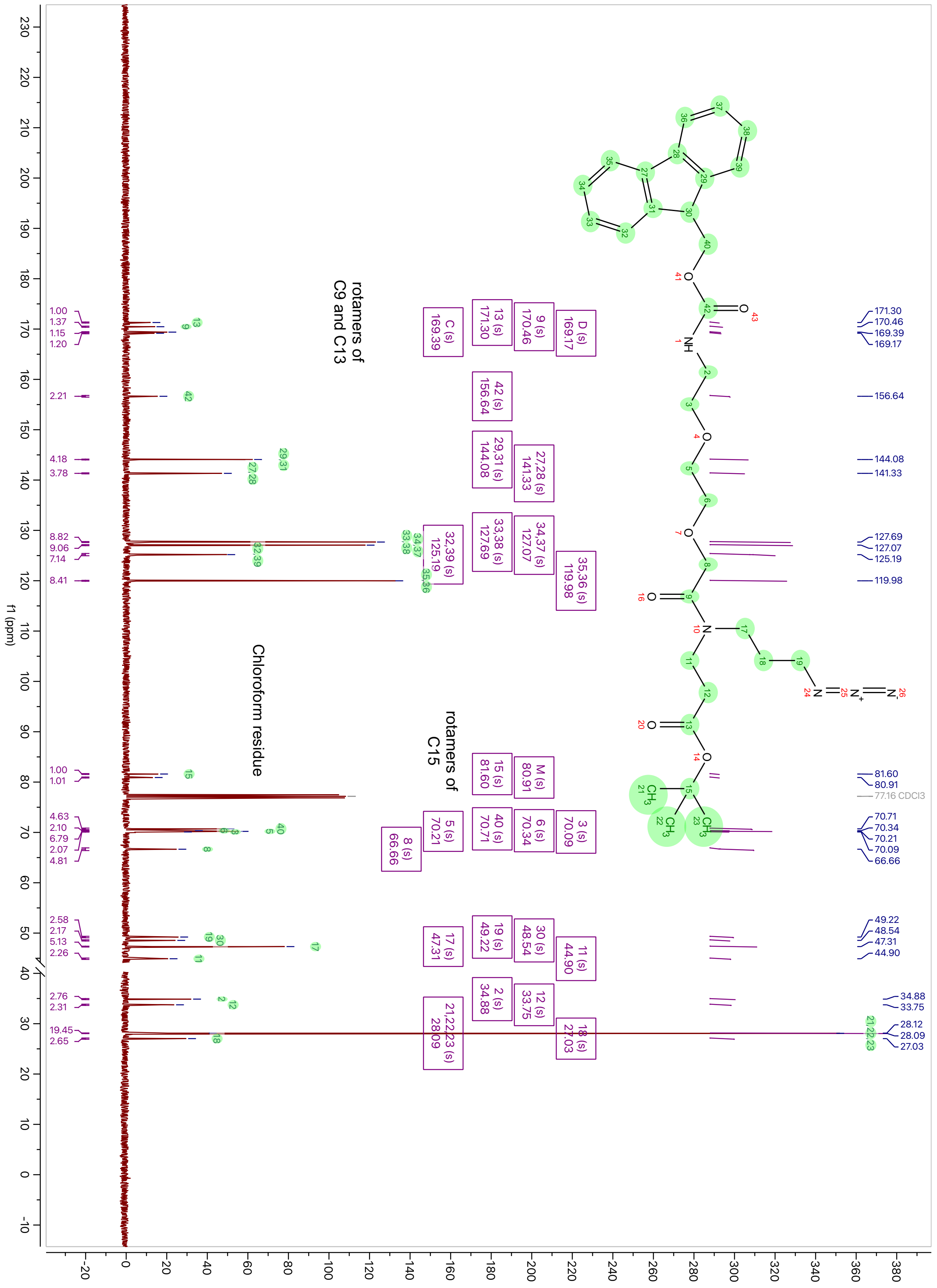

$^{1}$H-NMR spectrum of **4**

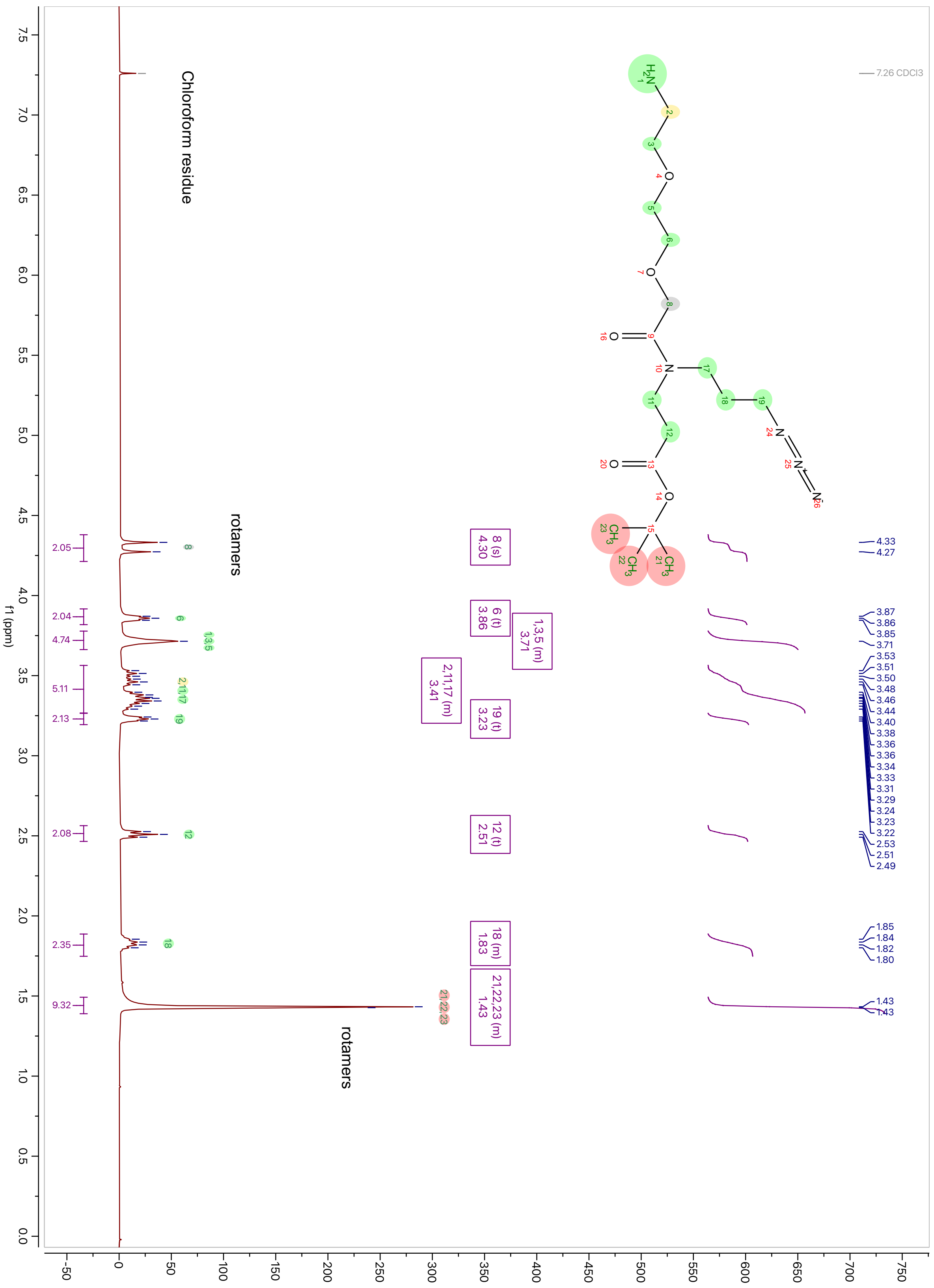

$^{13}$C-NMR spectrum of **4**

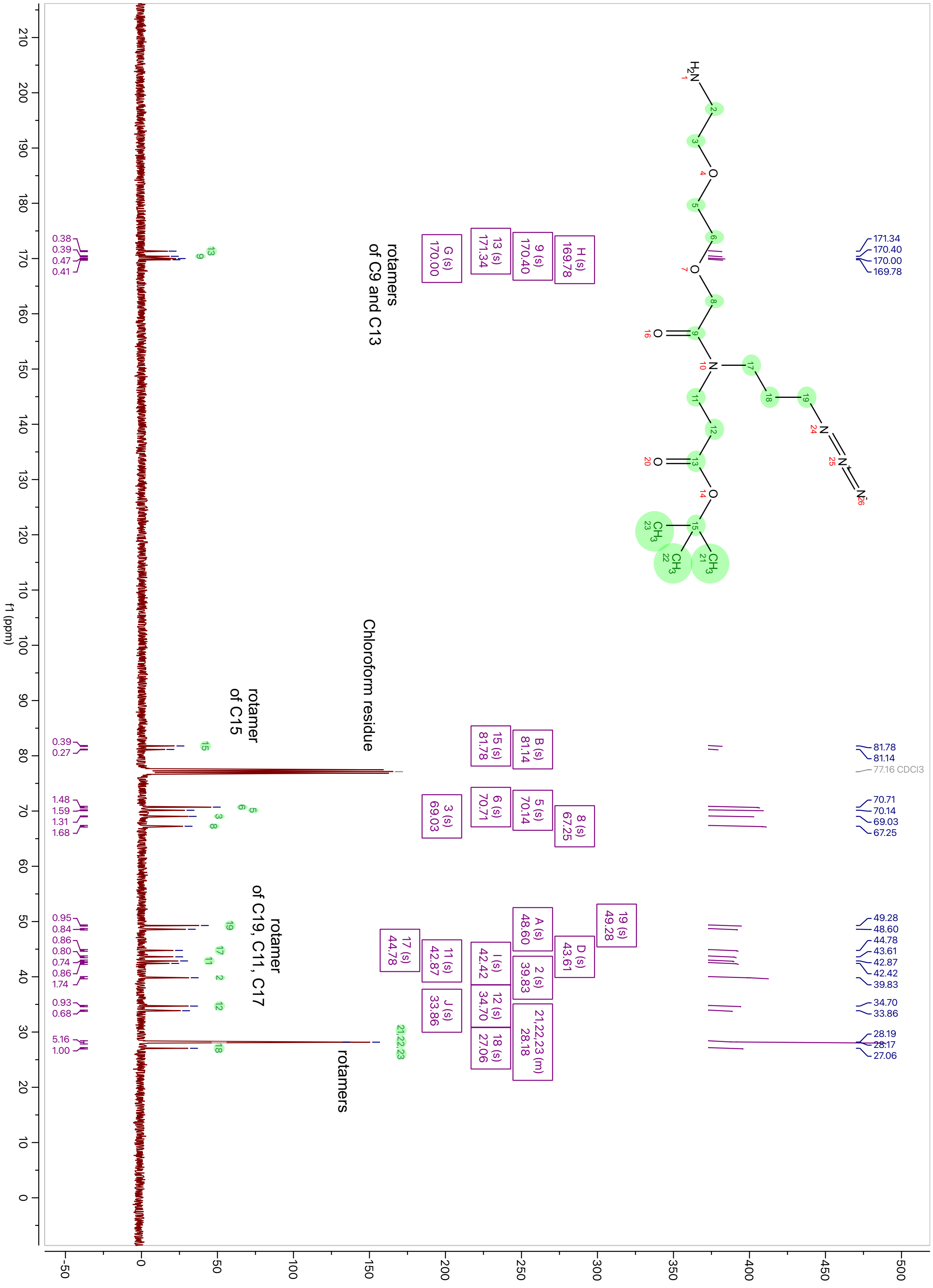

$^{1}$H-NMR spectrum of **5**

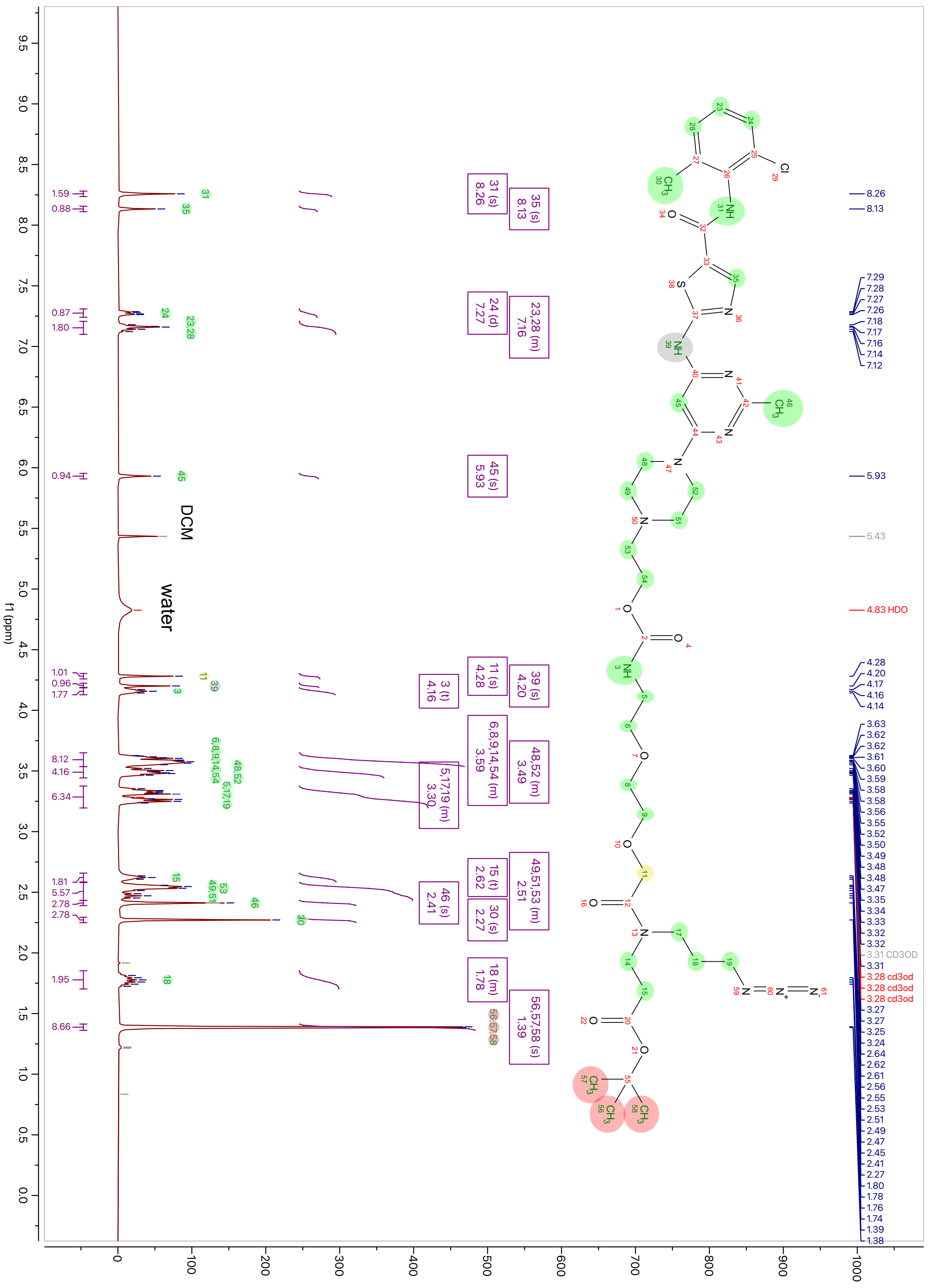

$^{13}$C-NMR spectrum of **5** (Note: The chemical shifts are partially altered by residual trifluoroacetic acid)

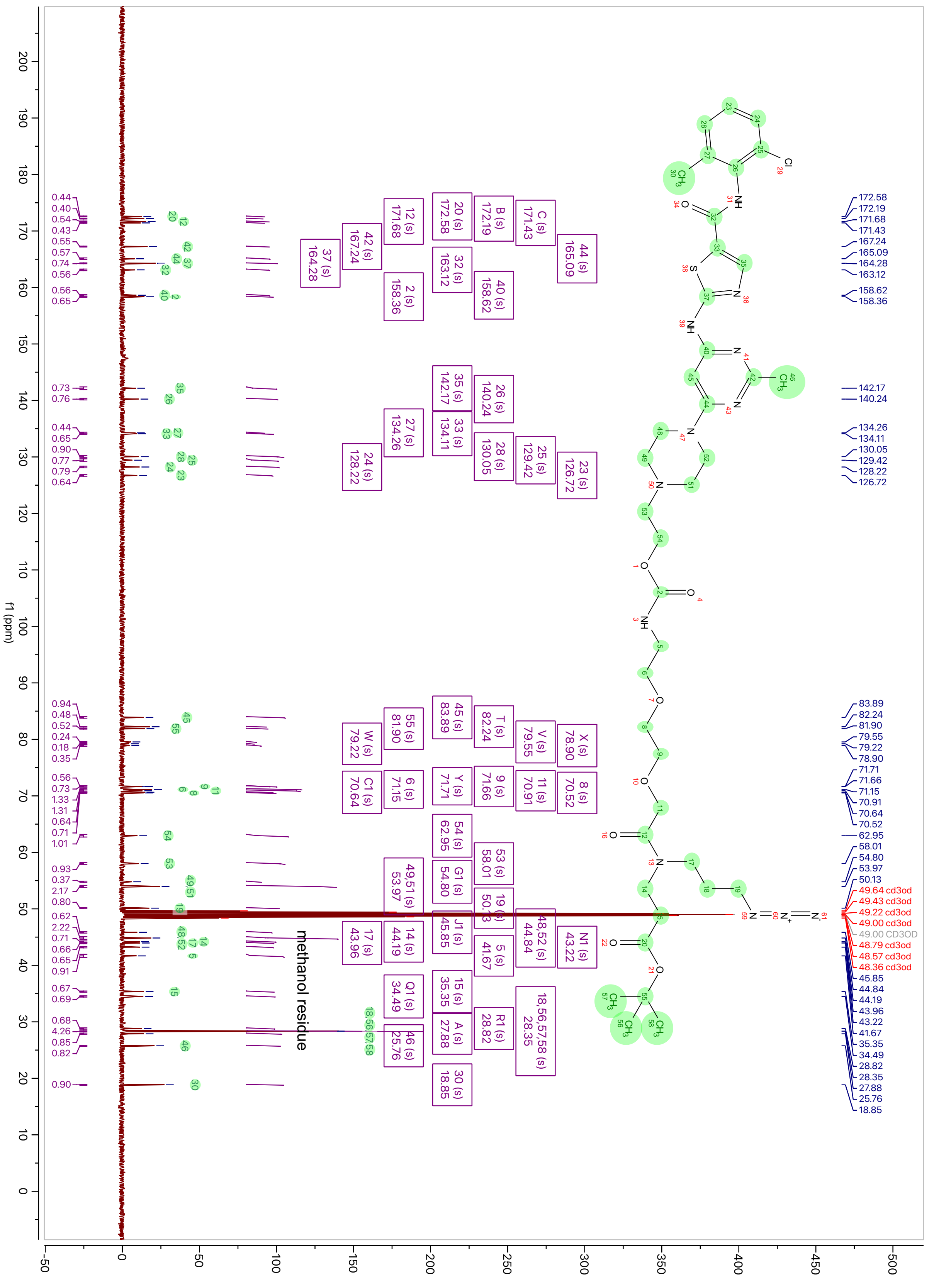

$^1$H-NMR spectrum of **6**

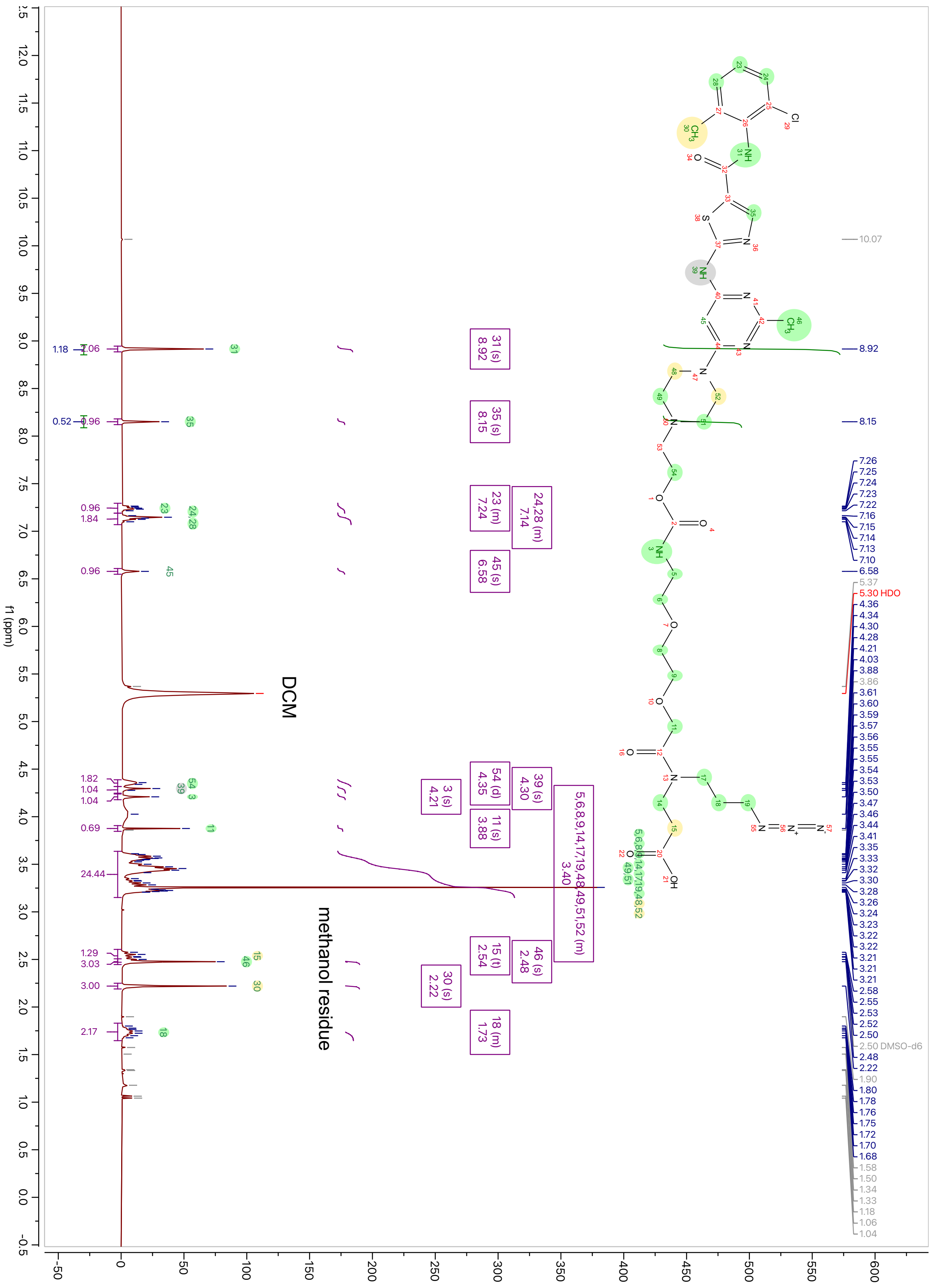

$^{13}$C-NMR spectrum of **6** (Note: The chemical shifts are partially altered by residual trifluoroacetic acid)

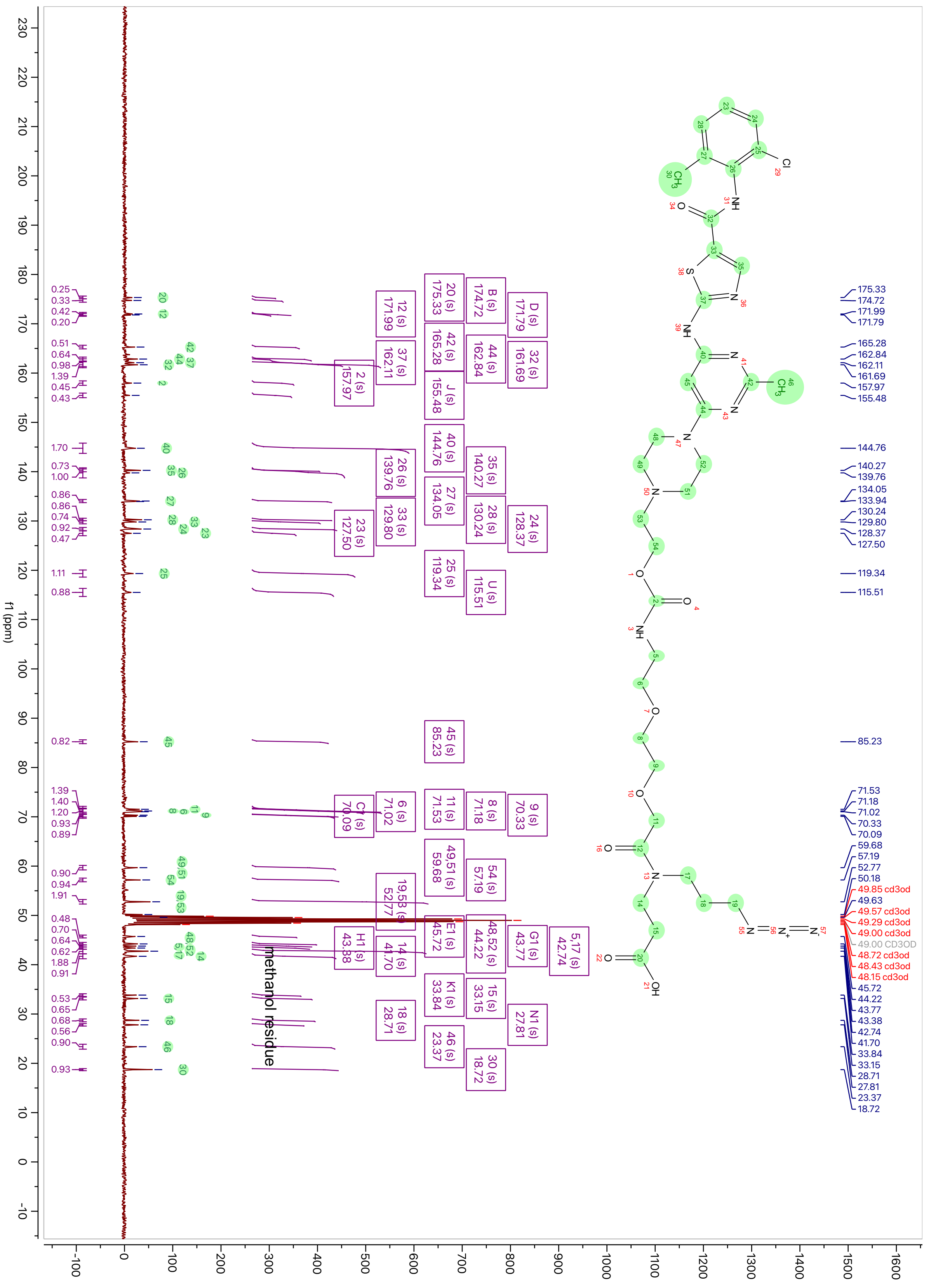

$^{1}$H-NMR spectrum of **Das-AI**

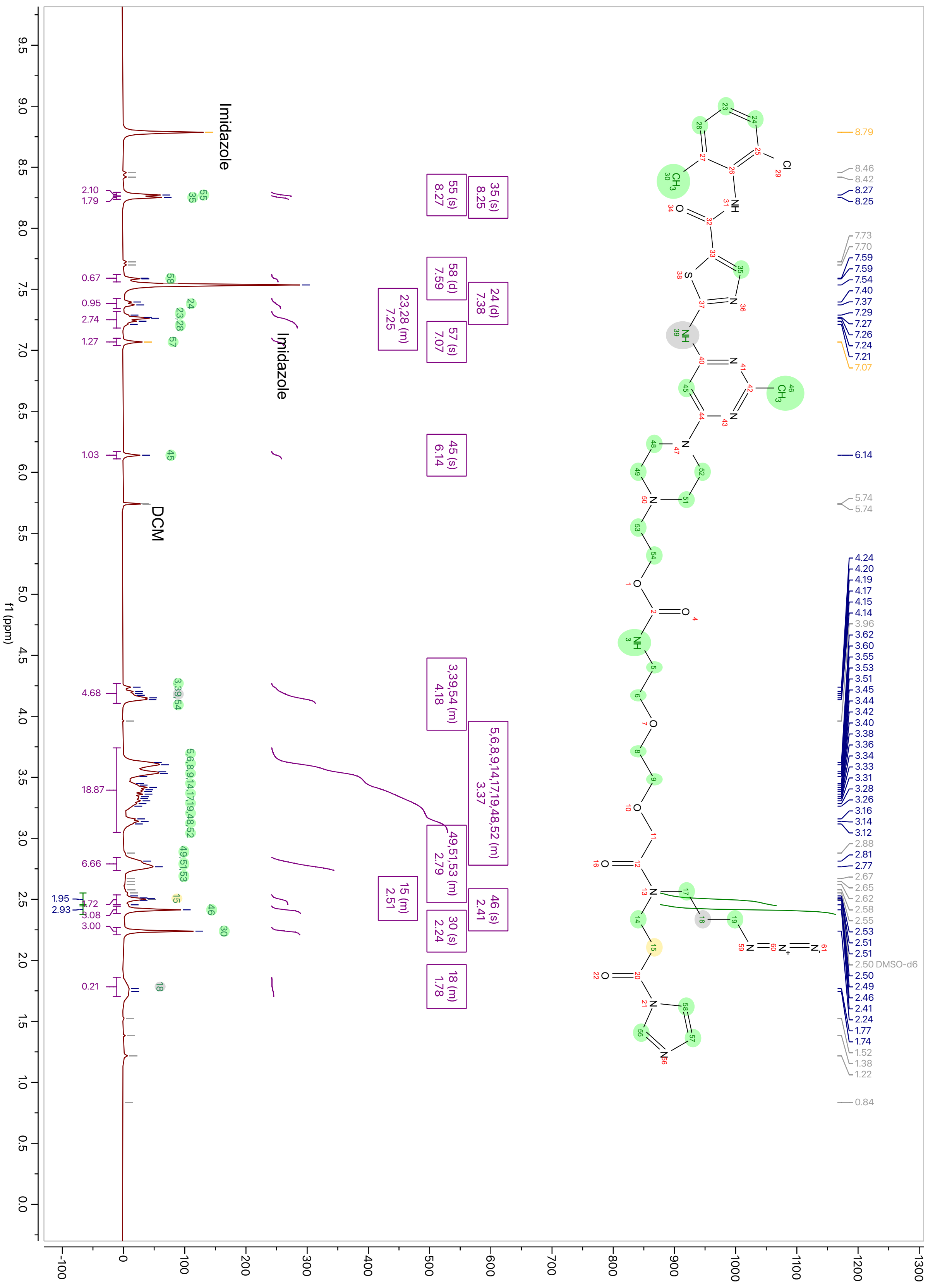

#### $^{13}$C-NMR spectrum of **Das-AI**

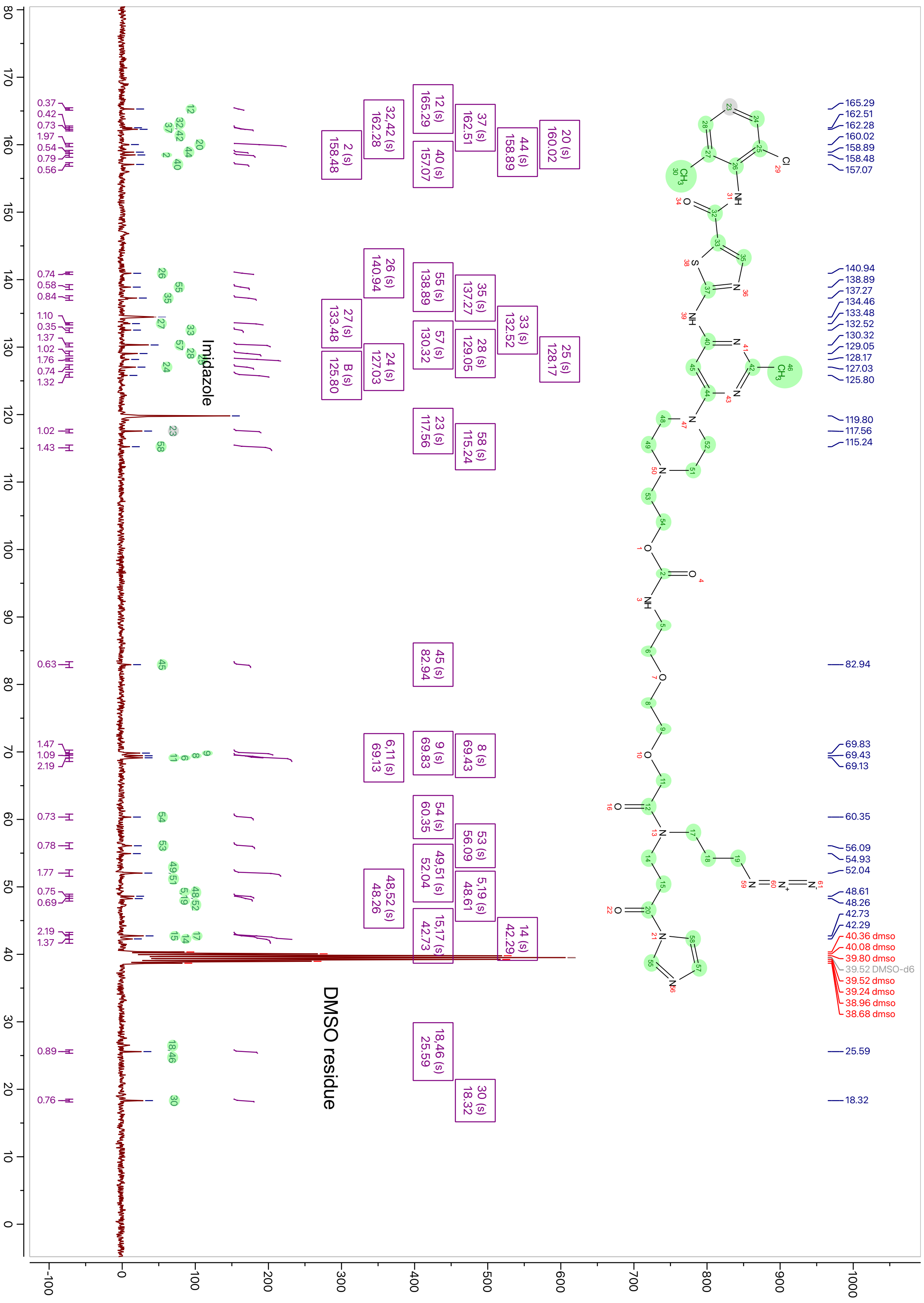

$^{1}$H-NMR spectrum of **7**

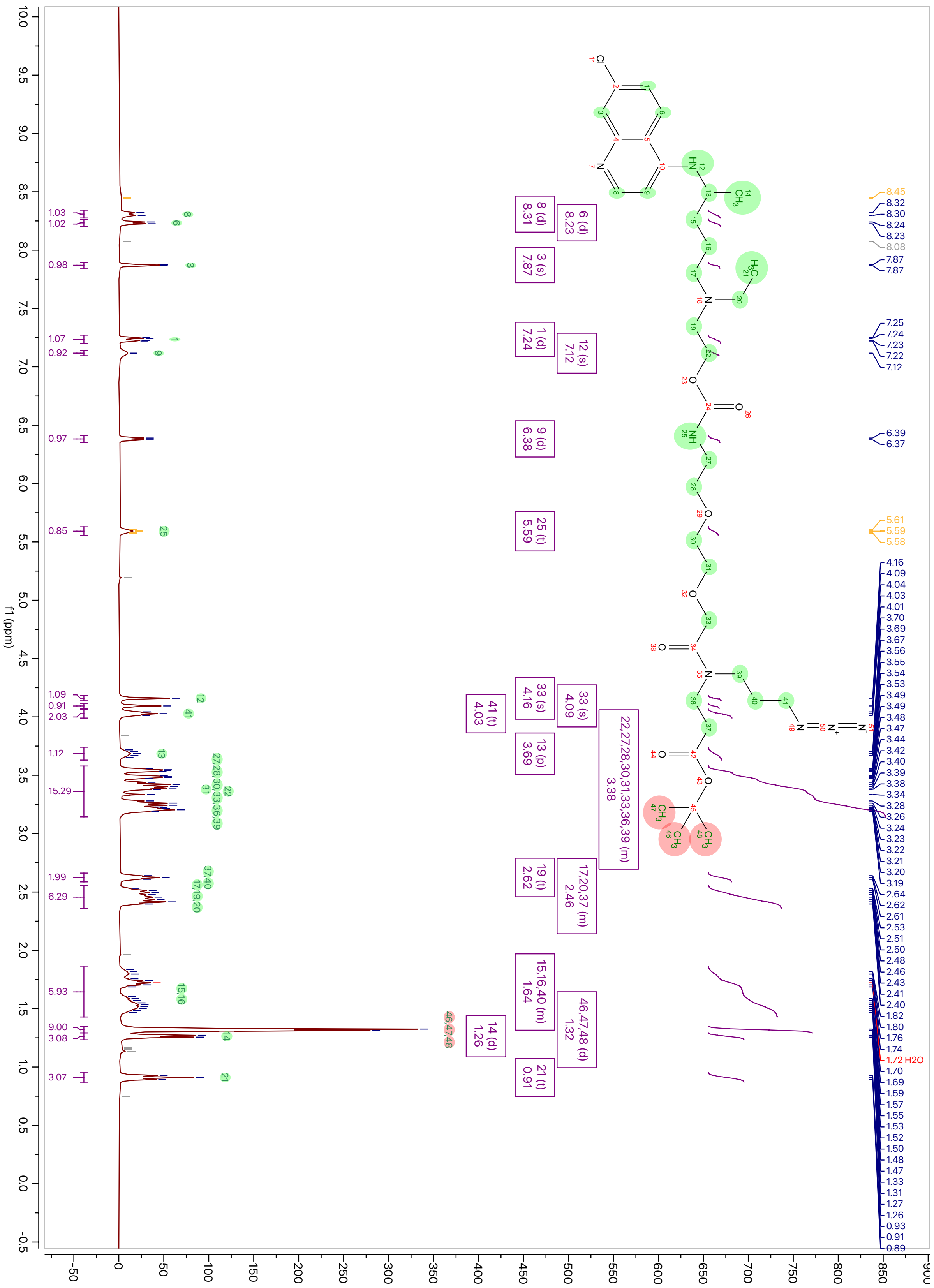

$^{13}$C-NMR spectrum of **7**

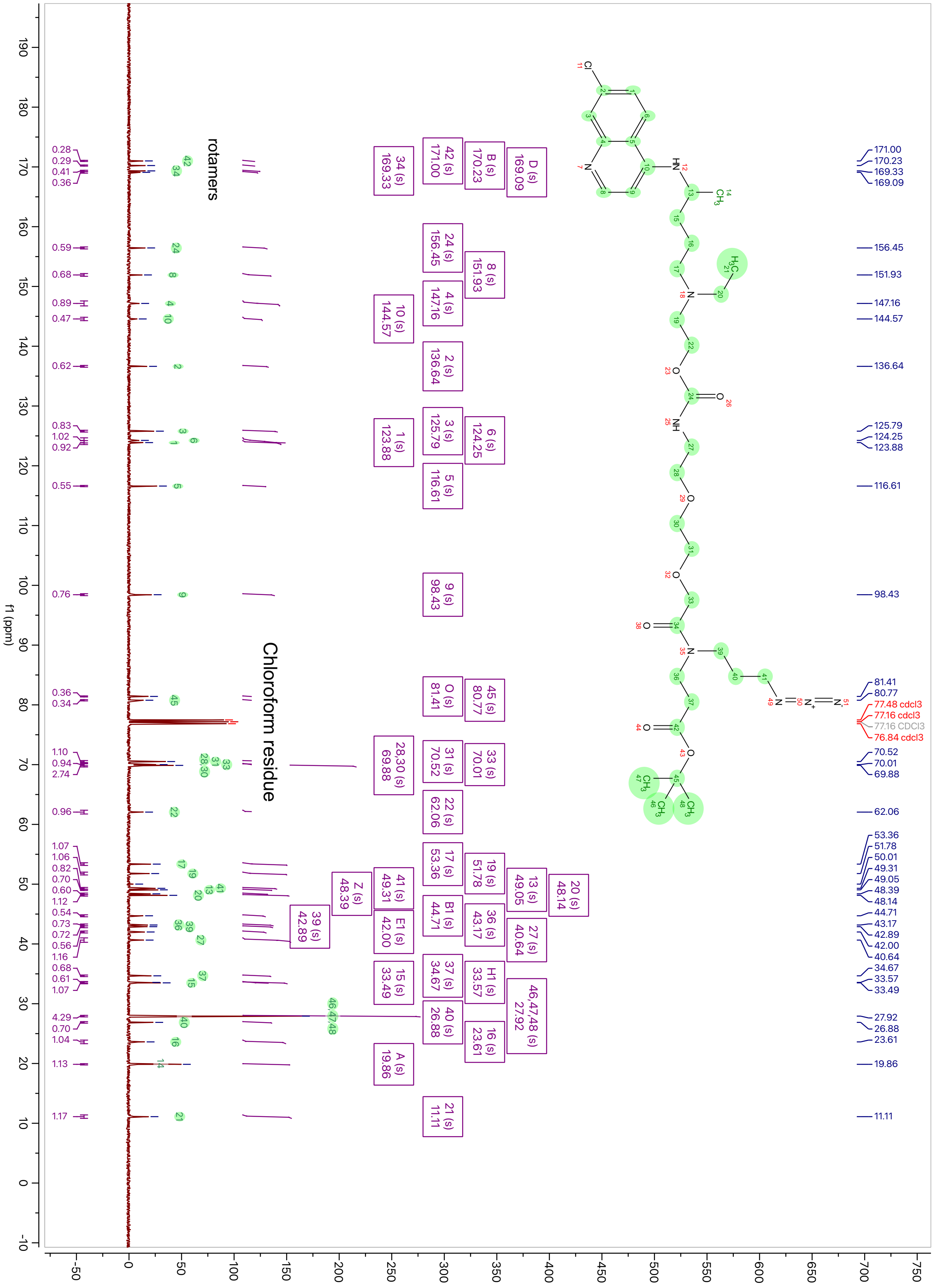

$^{1}$H-NMR spectrum of **8**

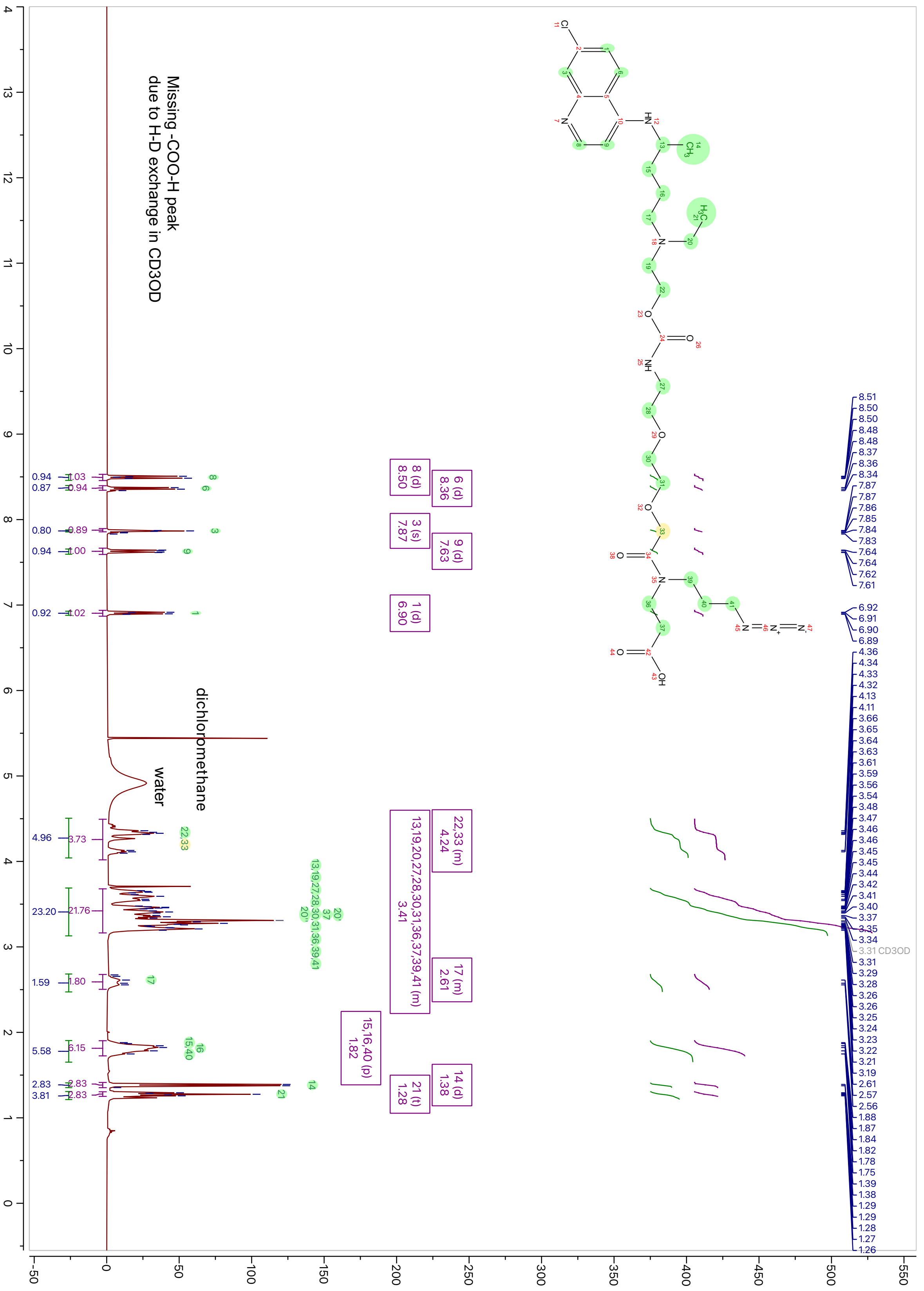

$^{13}$C-NMR spectrum of **8**

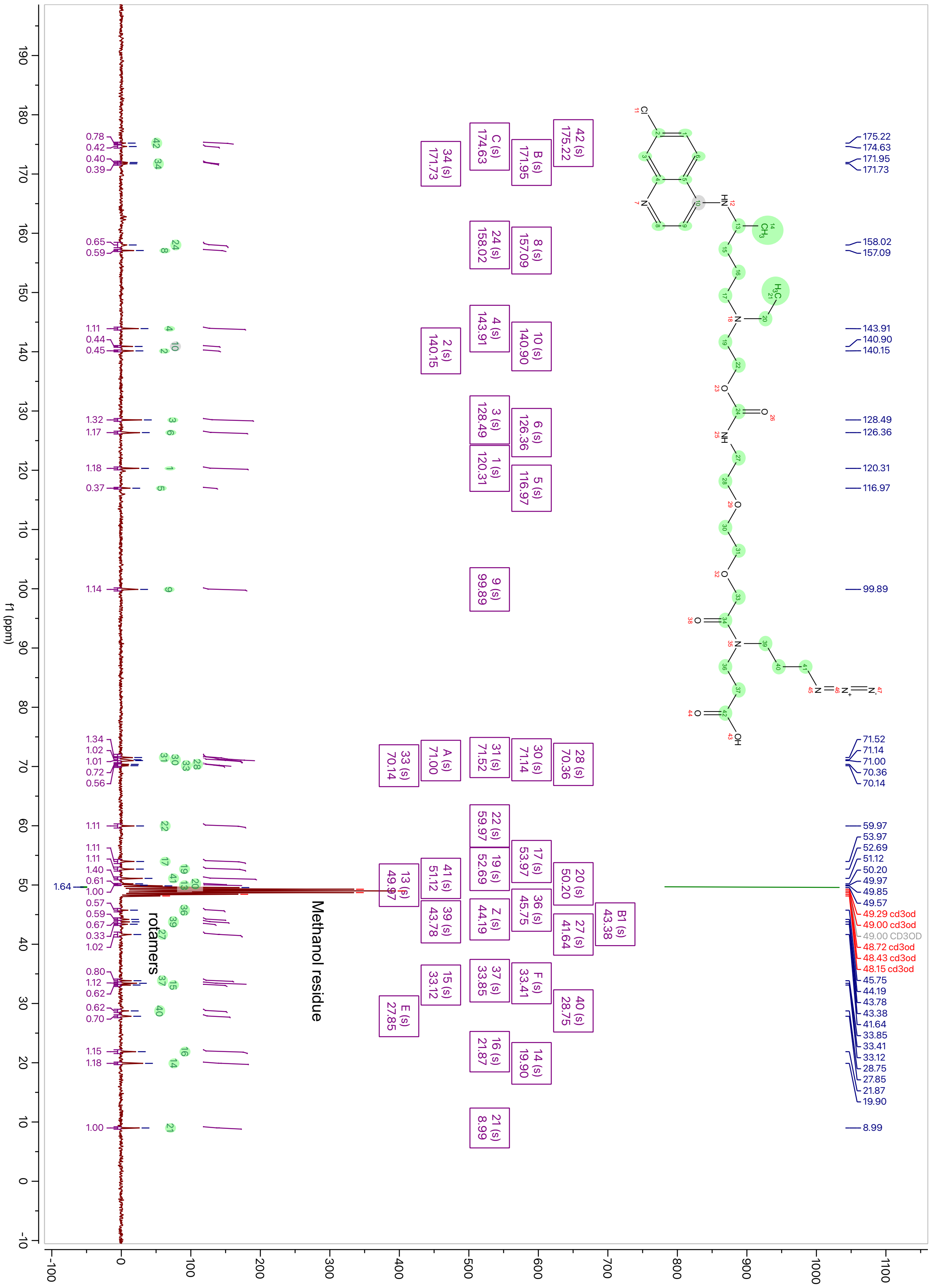

$^{1}$H-NMR spectrum of **HCQ-AI**

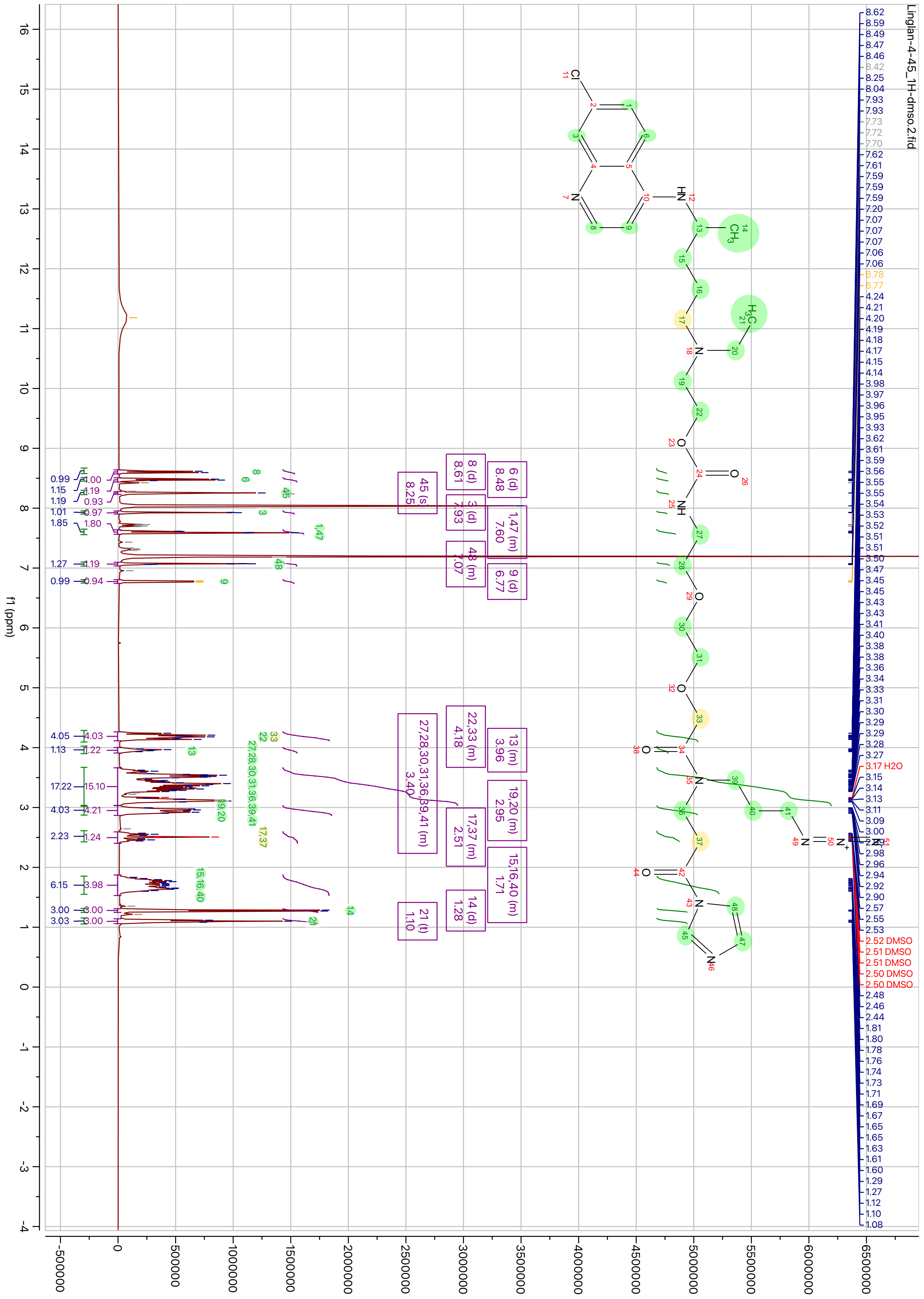

$^{13}$C-NMR spectrum of **HCQ-AI**

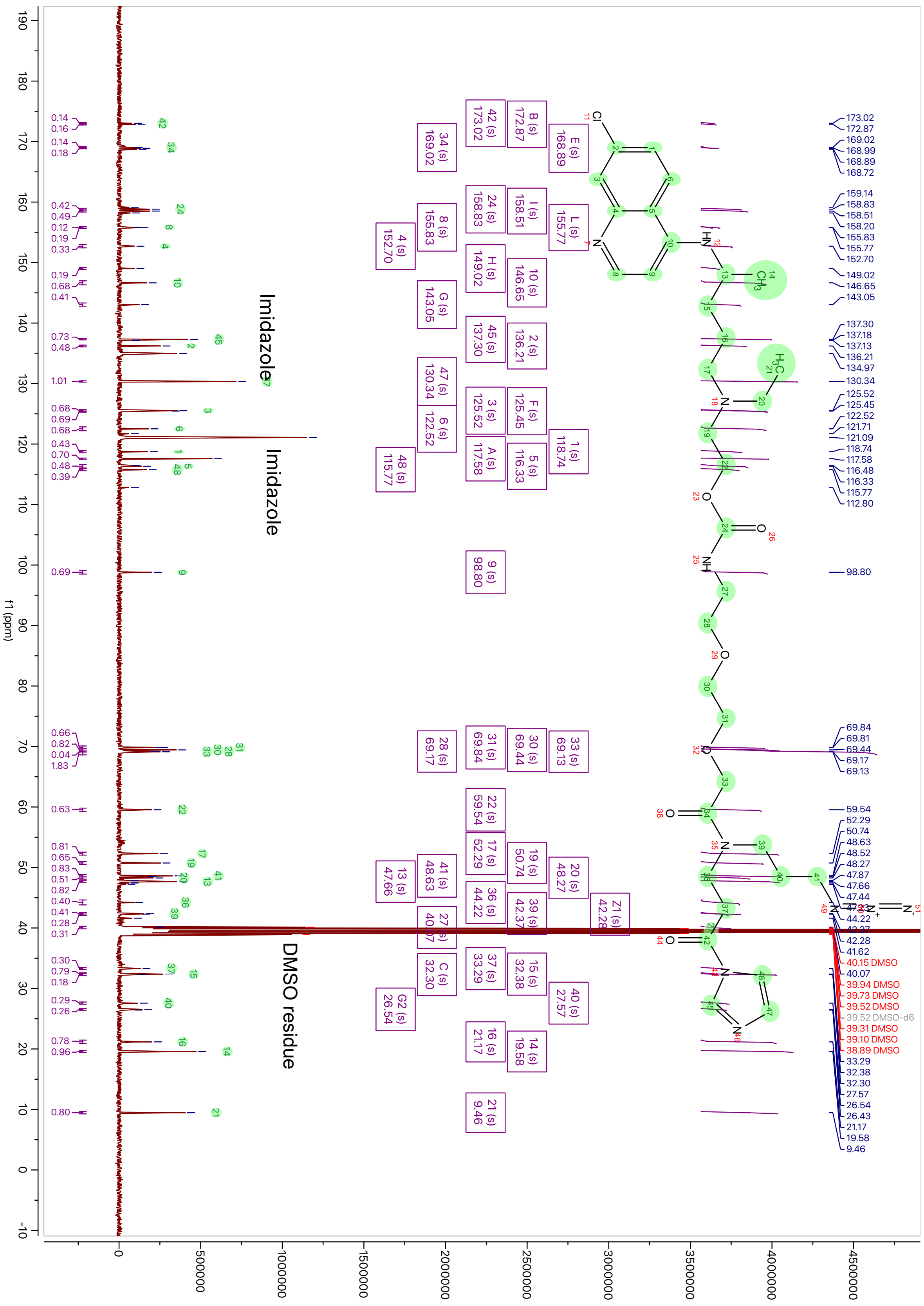

[1]H-NMR spectrum of **9** (This compound is a mixture of isomers)

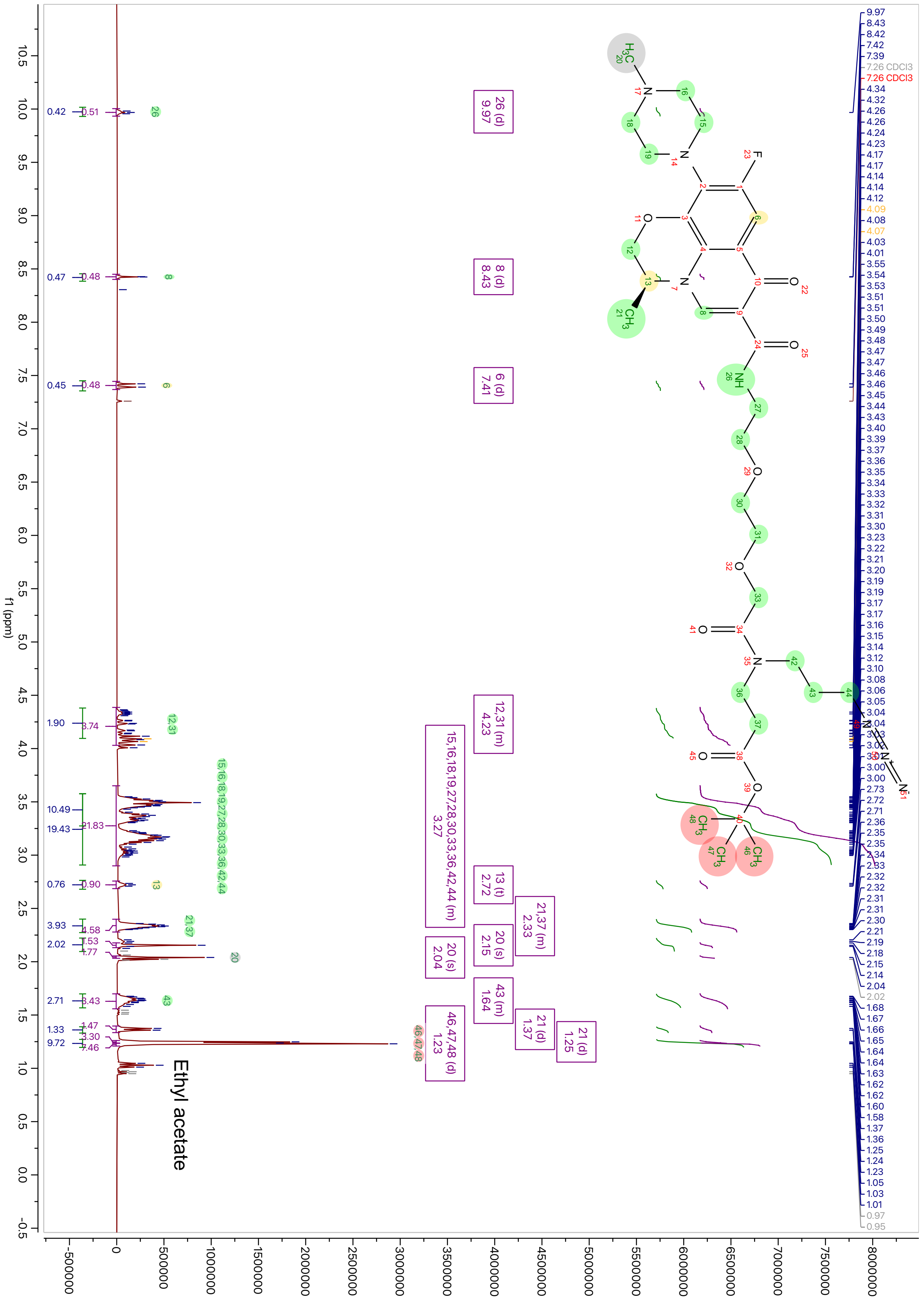

$^{13}$C-NMR spectrum of **9** (This compound is a mixture of isomers)

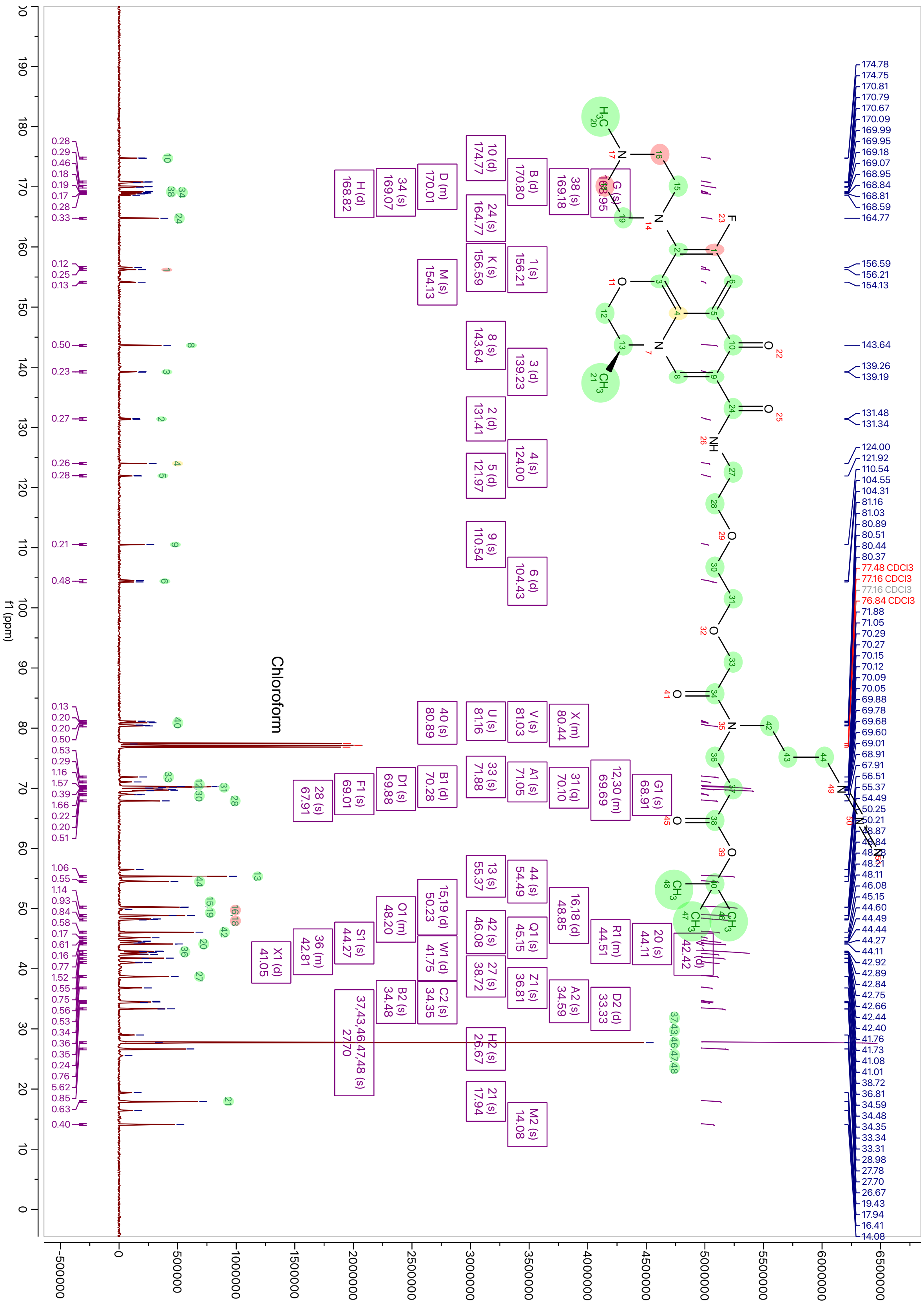

$^{1}$H-NMR spectrum of **10**

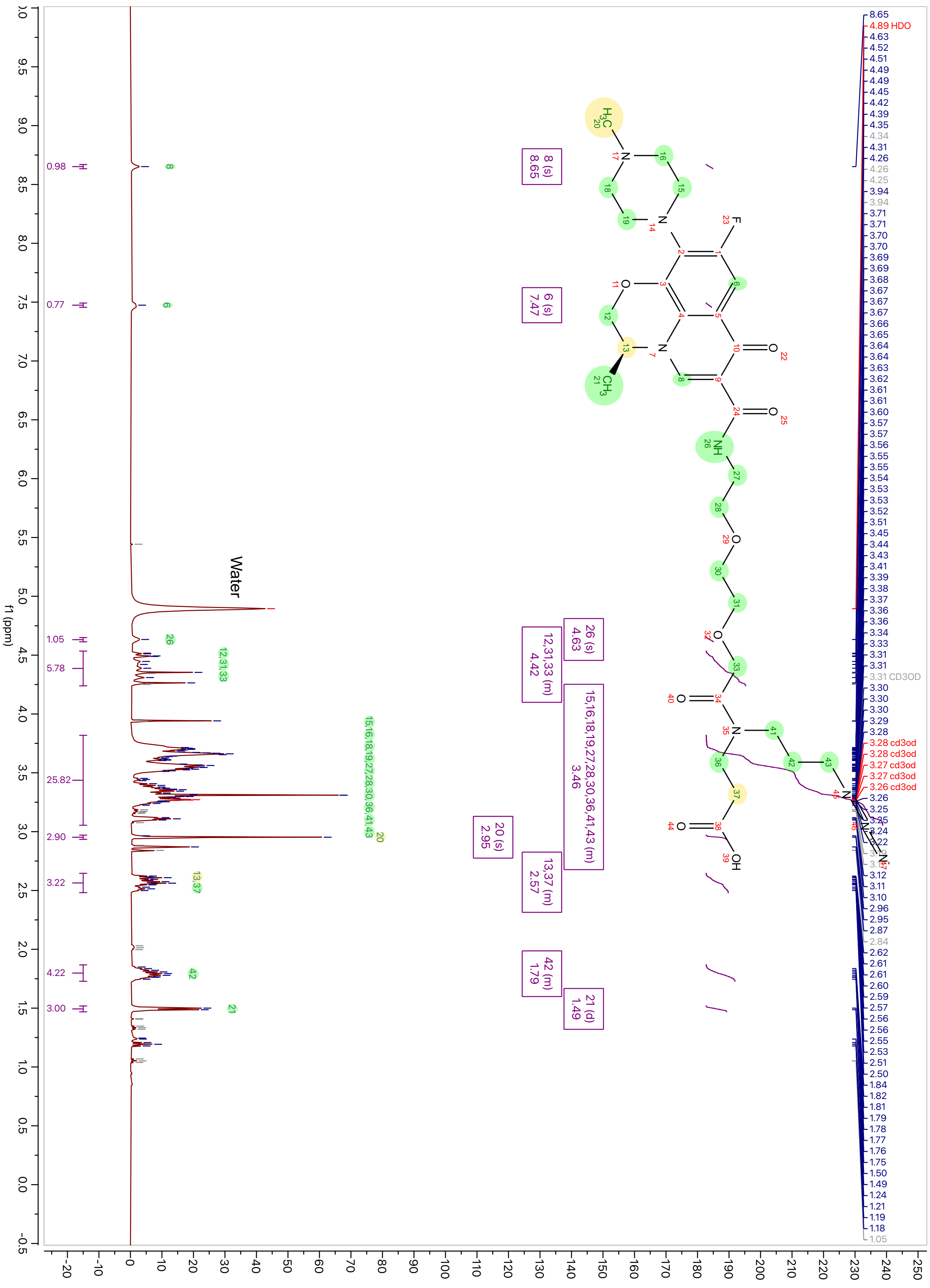

$^{13}$C-NMR spectrum of **10**

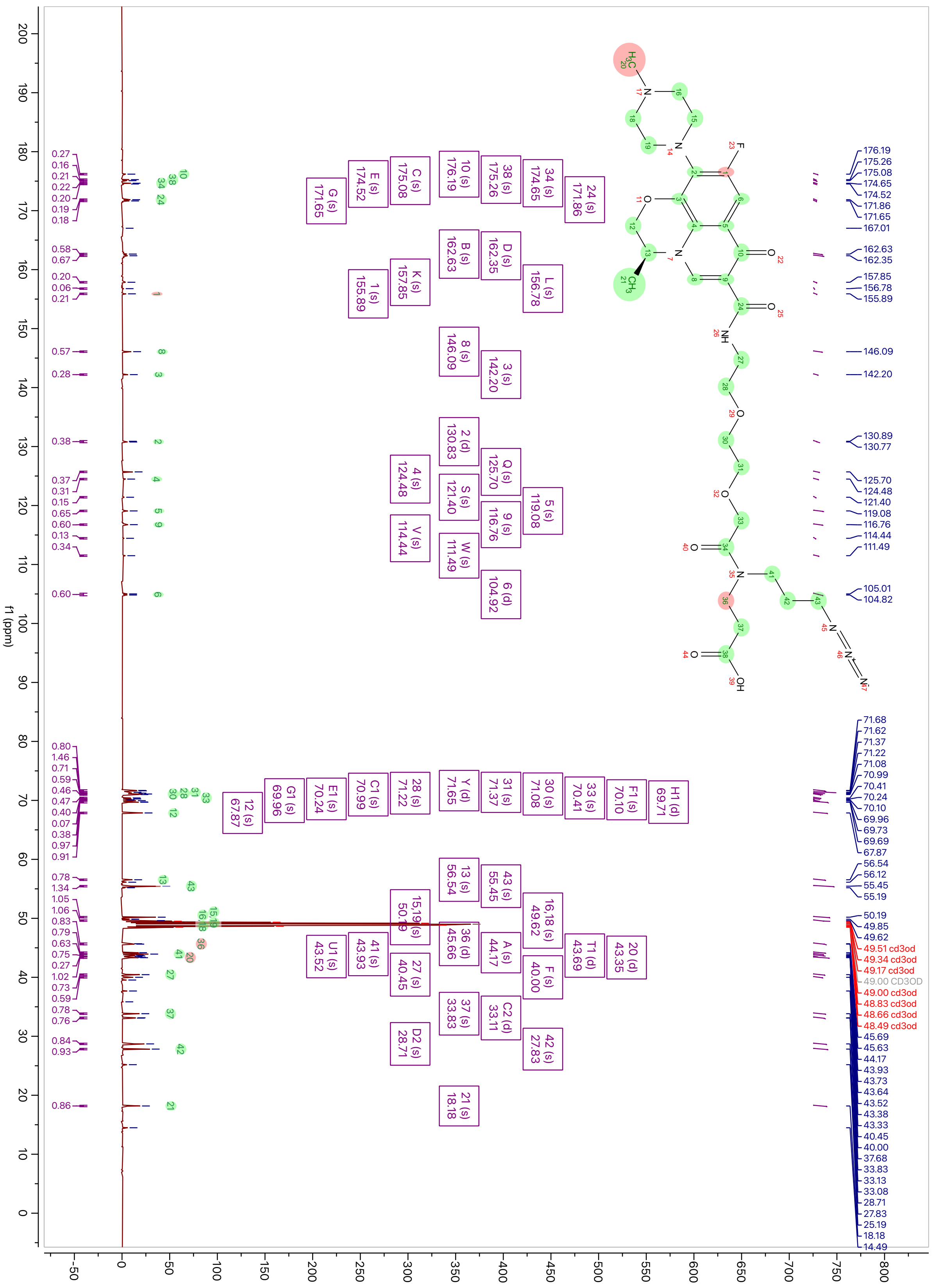

$^{1}$H-NMR spectrum of **Lev-AI**

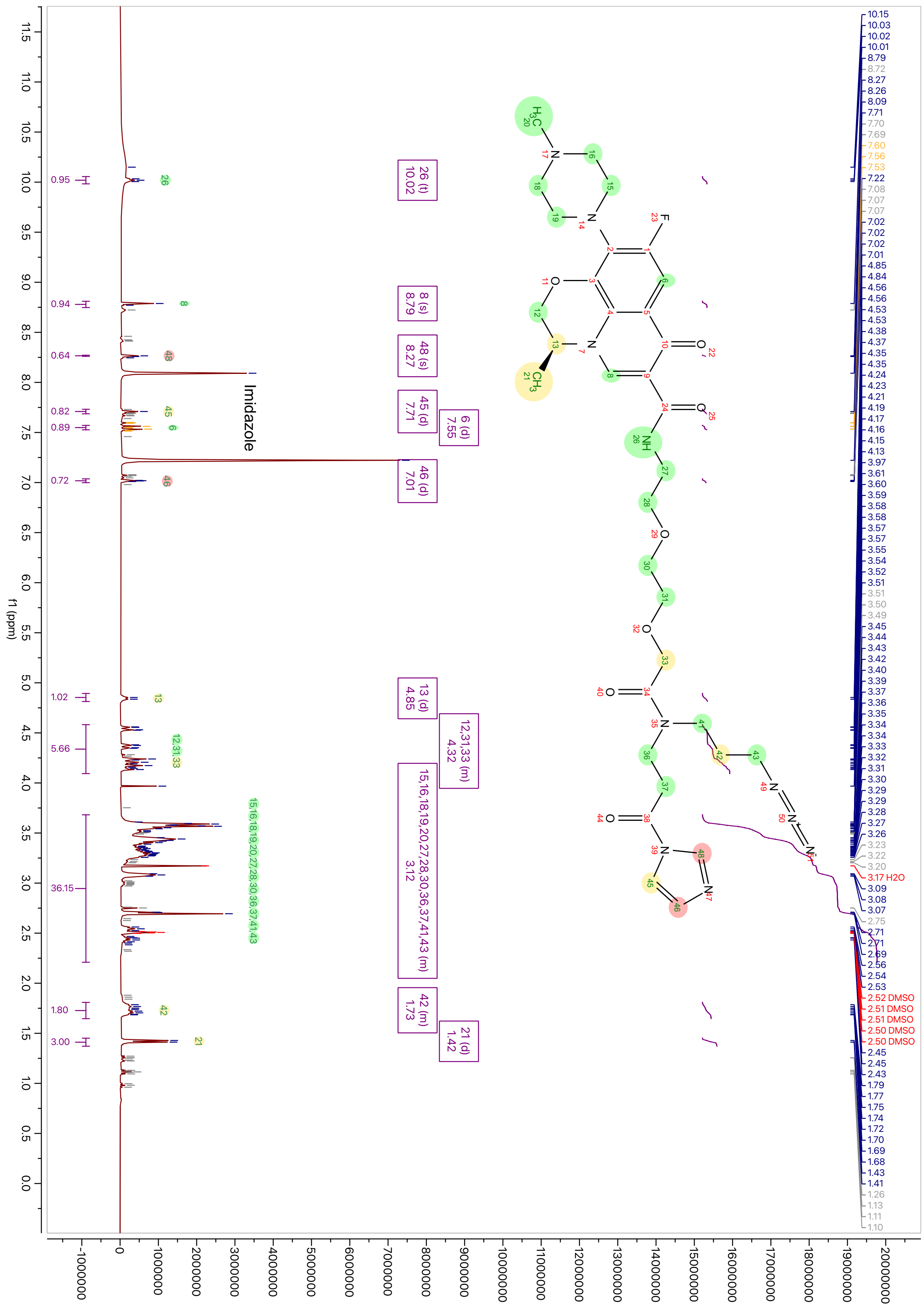

#### $^{13}$C-NMR spectrum of **Lev-AI**

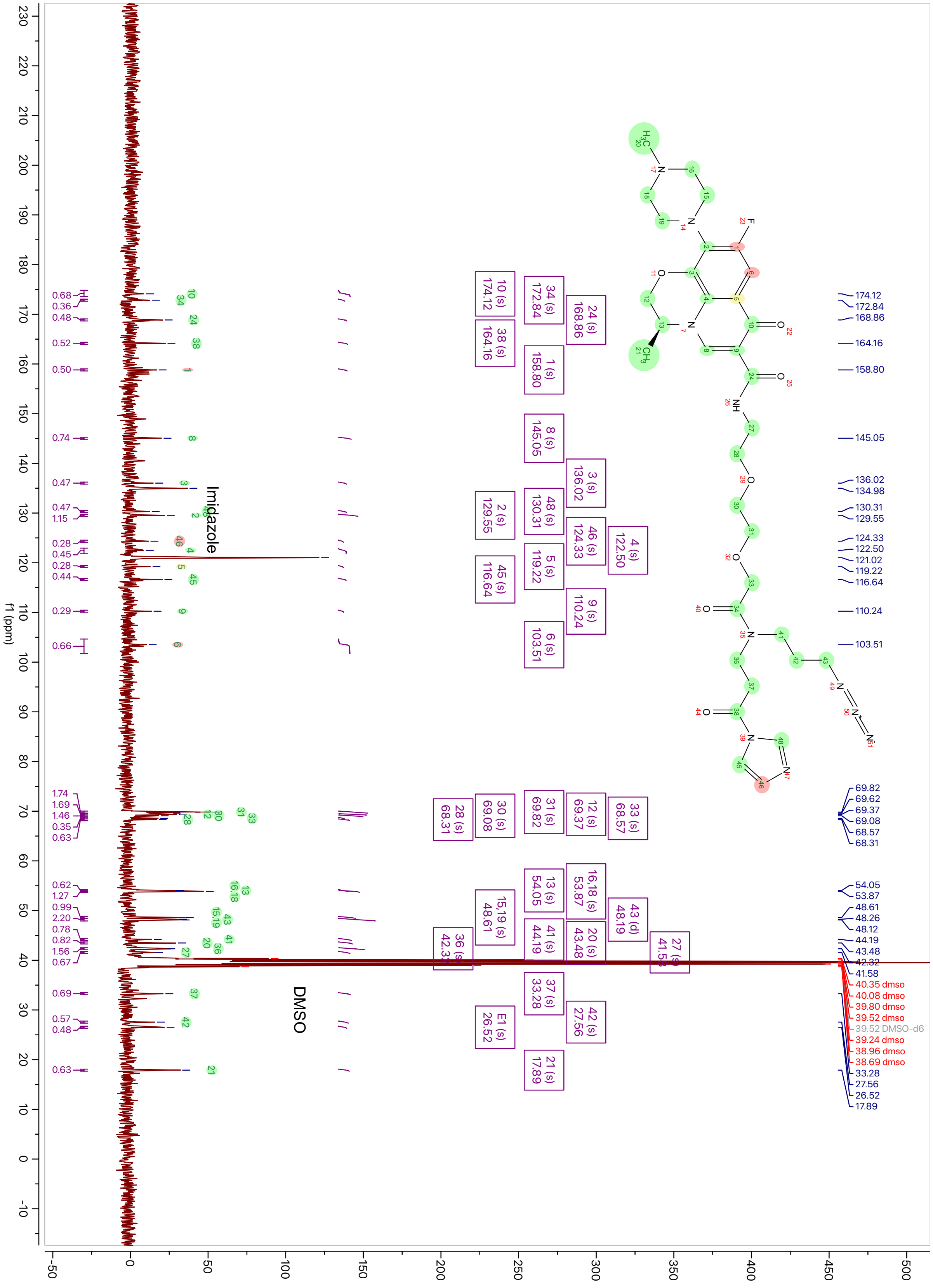

$^{1}$H-NMR spectrum of **Lev-diazirine**

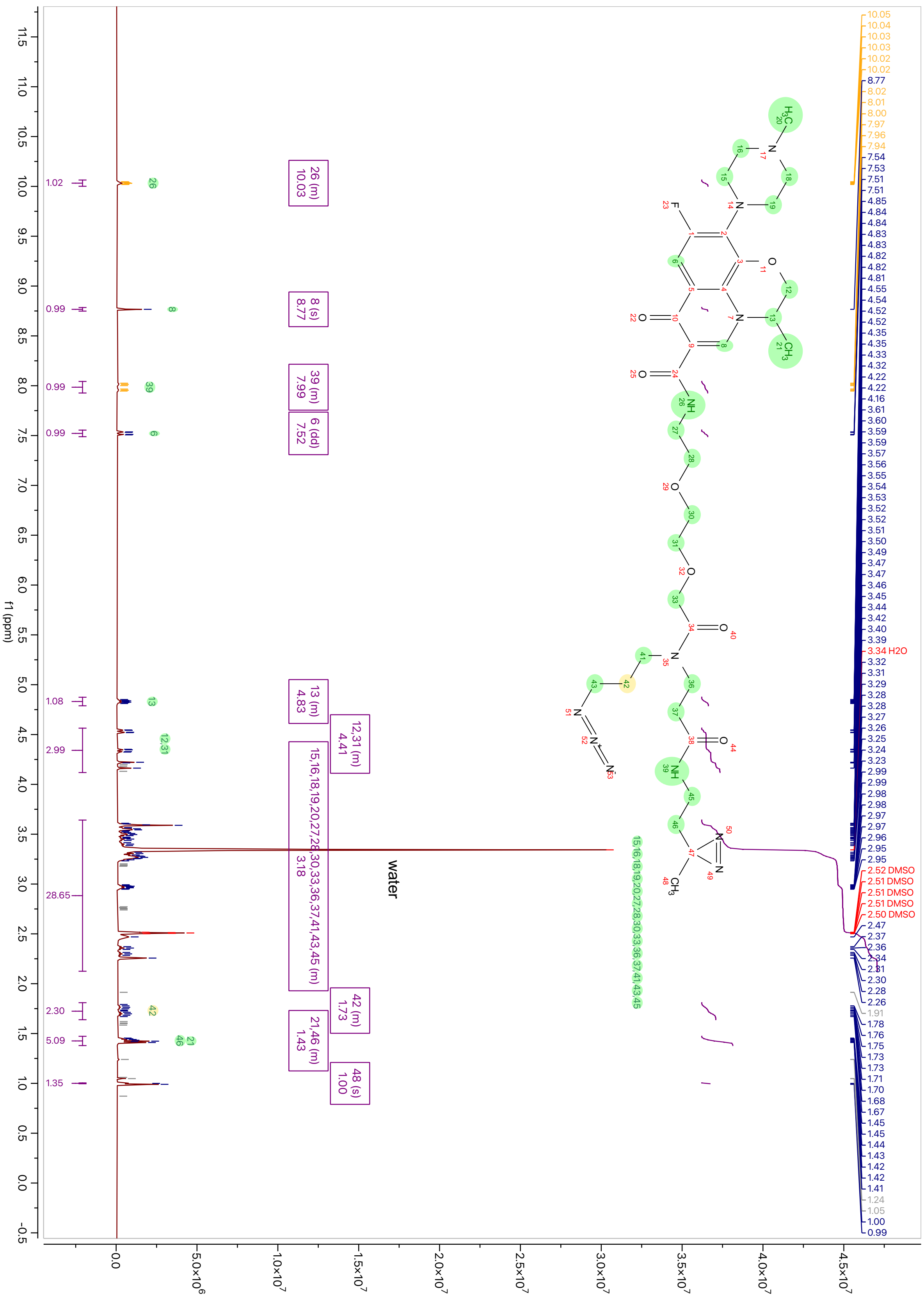

$^{13}$C-NMR spectrum of **Lev-diazirine**

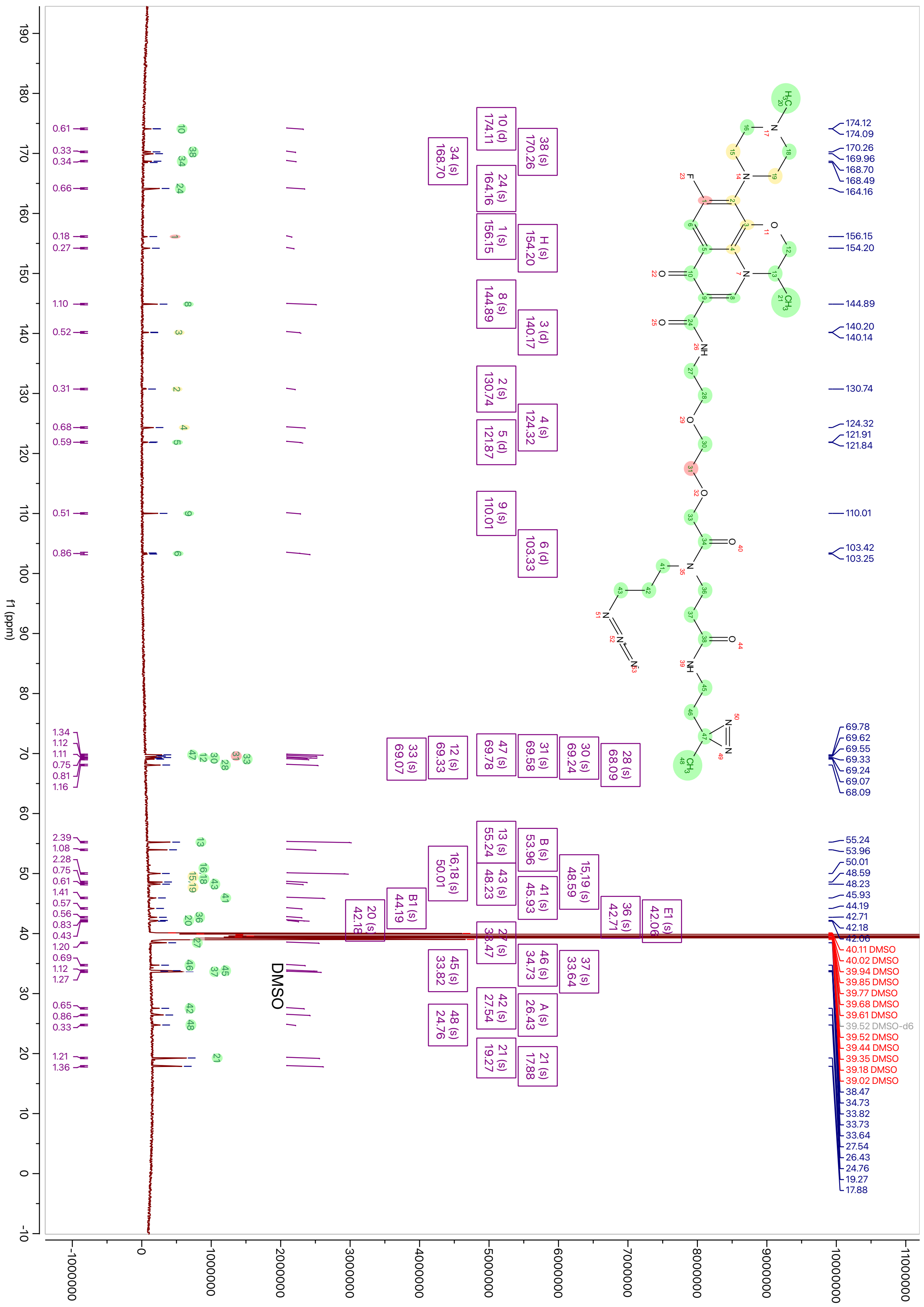

$^1$H-NMR spectrum of **11**

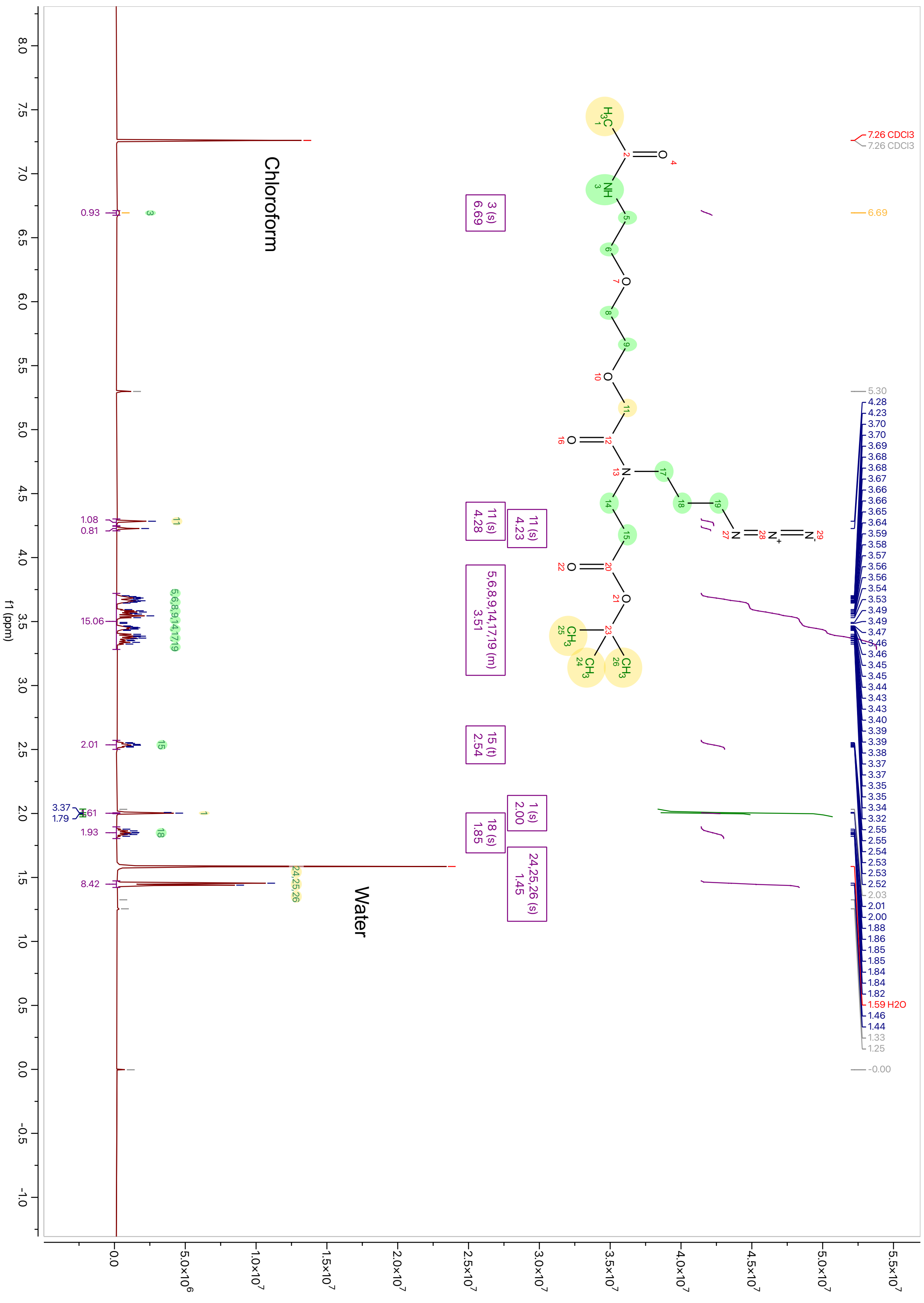

$^{13}$C-NMR spectrum of **11**

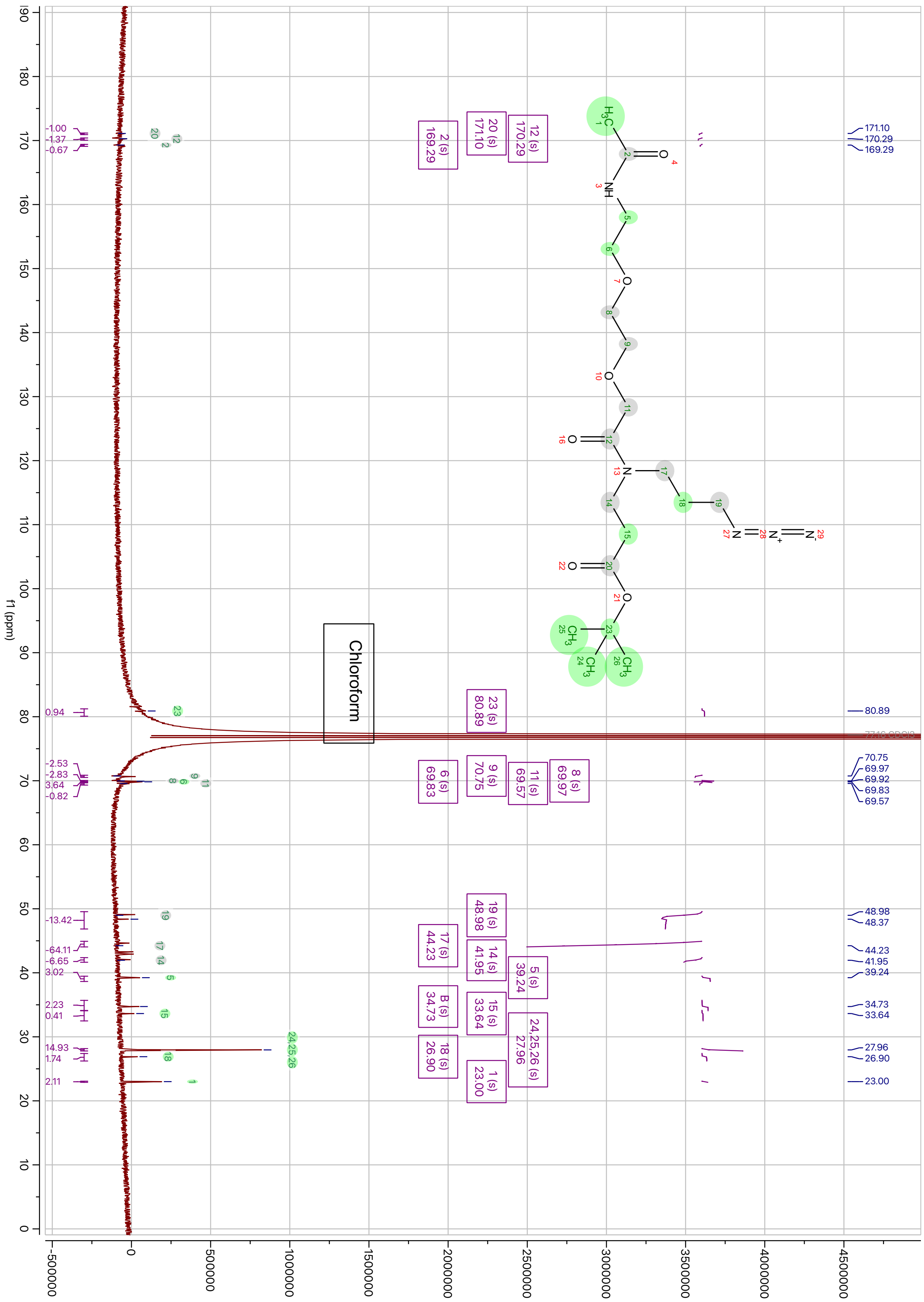

$^{1}$H-NMR spectrum of **12**

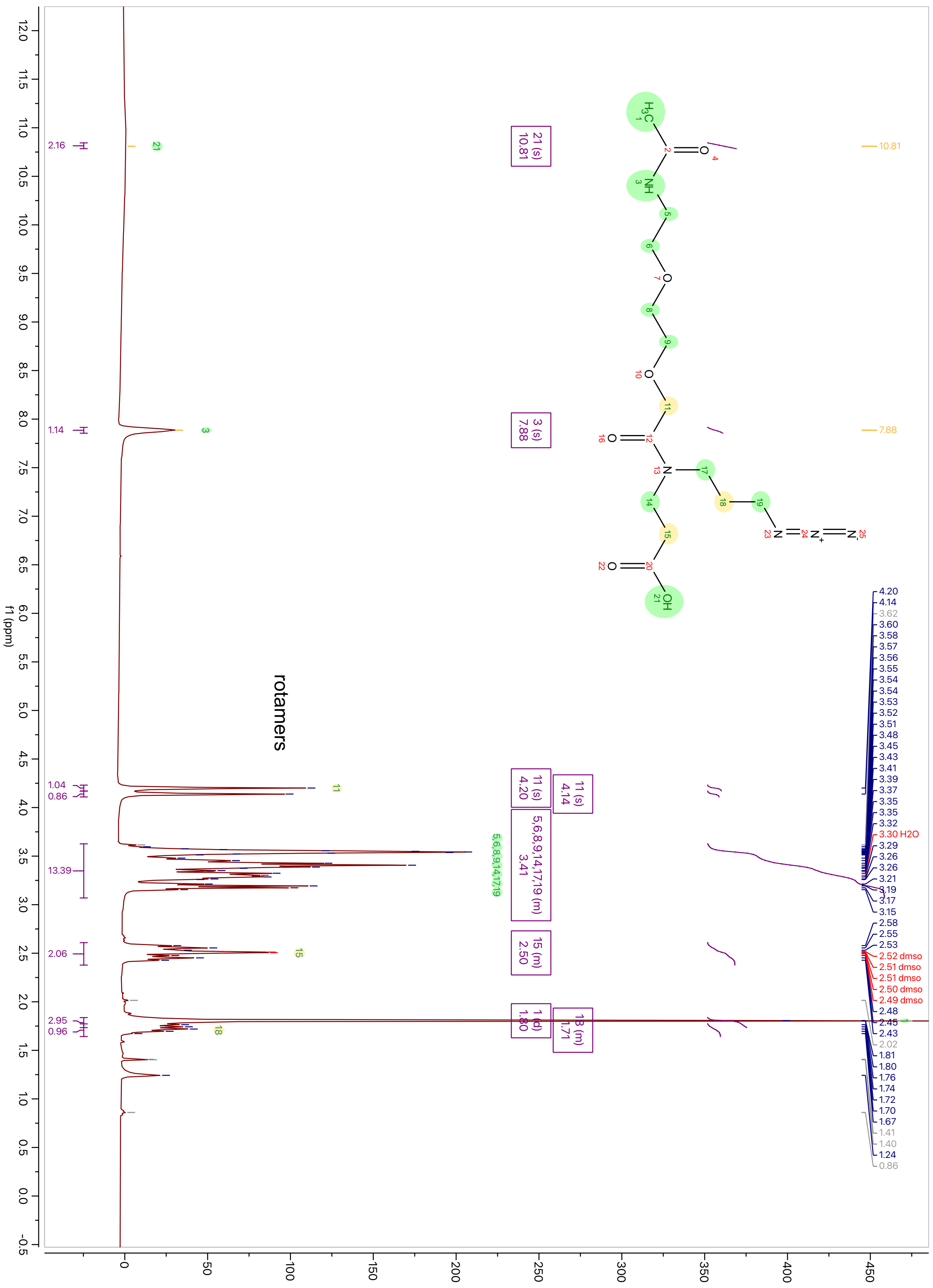

$^{13}$C-NMR spectrum of **12**

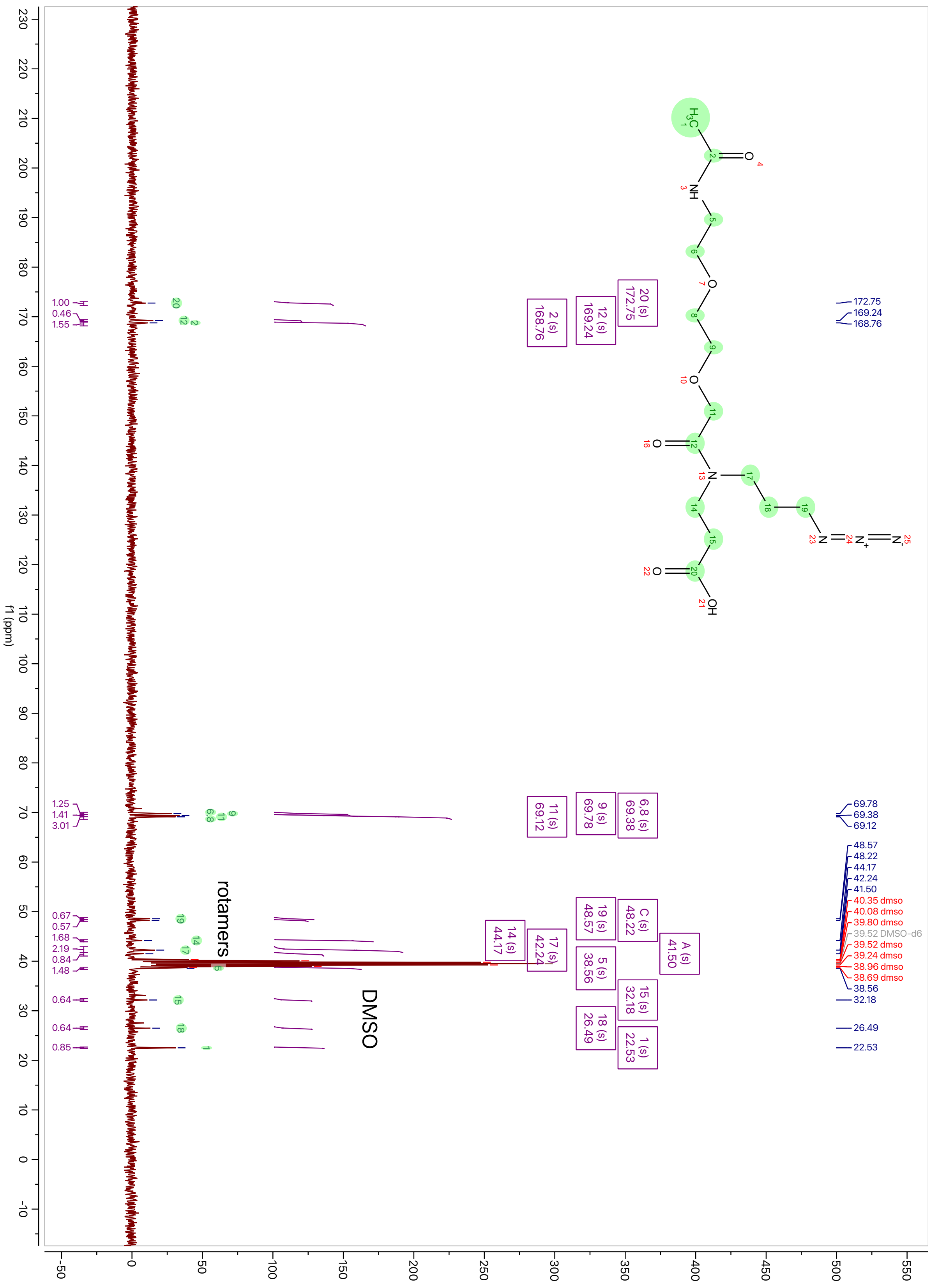

$^{1}$H-NMR spectrum of **Linker-AI**

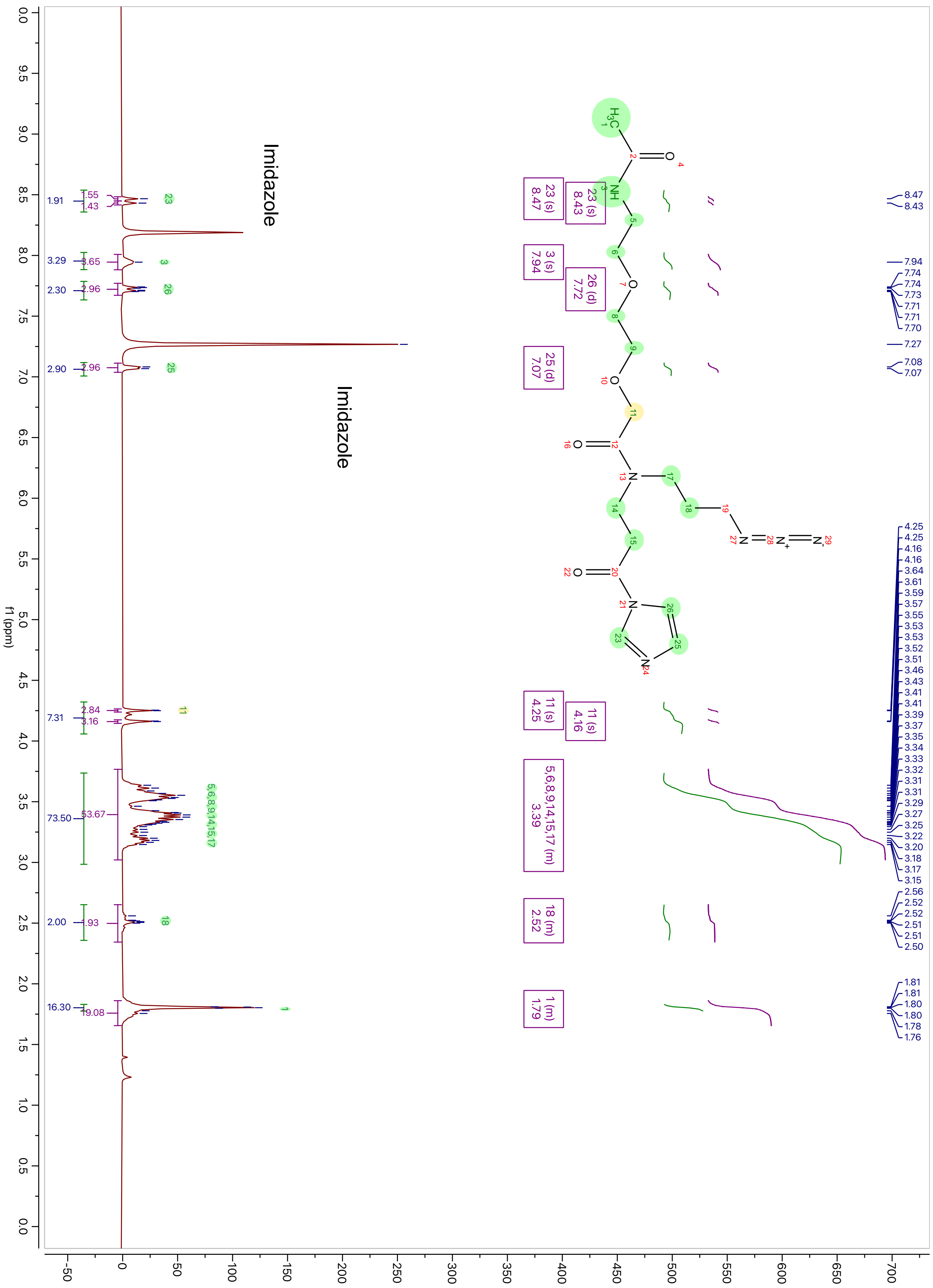

$^{13}$C-NMR spectrum of **Linker-AI**

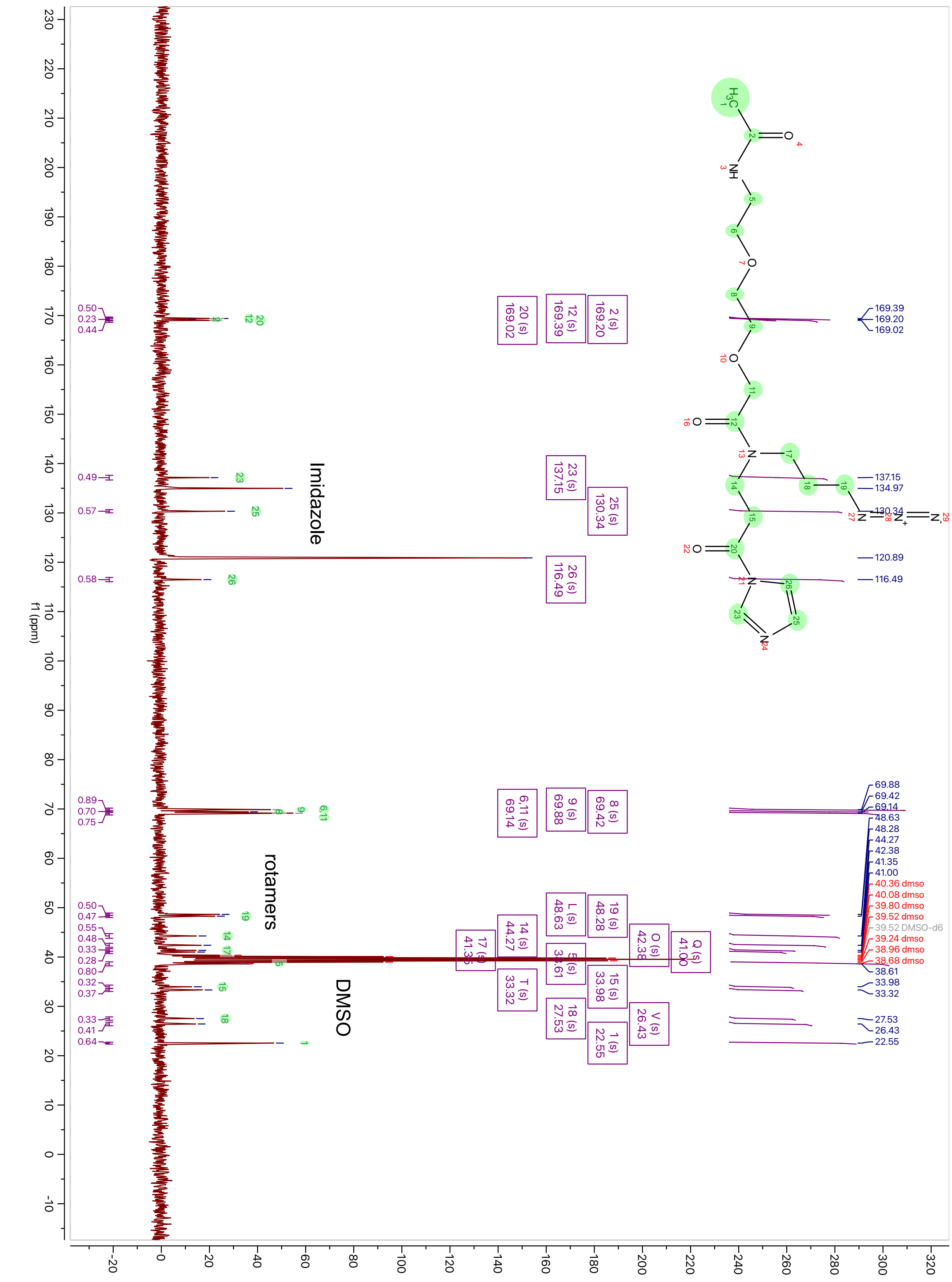

**High-Resolution Mass Spectrometry (HRMS) data:** The sample was analyzed by LC-flow injection ESI/MS on the Waters Acquity H-Class Plus UPLC and Thermo Exploris 240 BioPharma Orbitrap mass spectrometer. Acetonitrile with 0.1% formic acid at a flow rate of 0.2mL/min was used to transport the injected sample to the source directly, without chromatographic separation. The injection volume was 5uL. Spectra were collected in full scan MS mode with polarity switching (collecting scans alternating between positive and negative ionization potentials), Orbitrap resolution 120000, mass range of 100-1000 Da. RunStart Easy IC Internal Mass Calibration was enabled.

*Tert*-butyl 3-((3-azidopropyl)amino)propanoate (**2**):

tmcl_X240_18924.raw, C10H20N4O2 [H]+

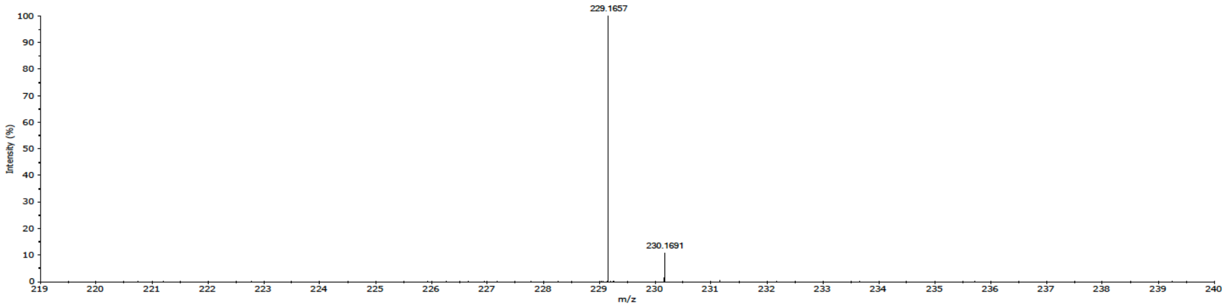


*Tert*-butyl 13-(3-azidopropyl)-1-(9*H*-fluoren-9-yl)-3,12-dioxo-2,7,10-trioxa-4,13-diazahexadecan-16-oate (**3**)

tmcl_X240_18925.raw, C31H41N5O7 [H]+

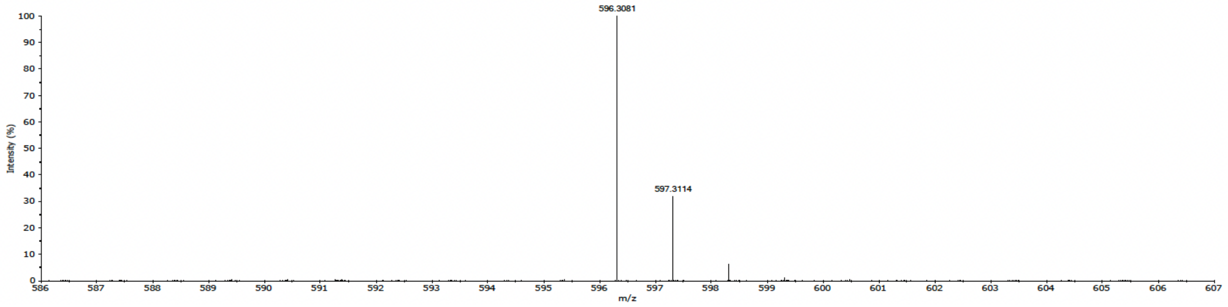


*Tert*-butyl 3-(2-(2-(2-aminoethoxy)ethoxy)-*N*-(3-azidopropyl)acetamido)propanoate (**4**)

tmcl_X240_18926.raw, C16H31N5O5 [H]+

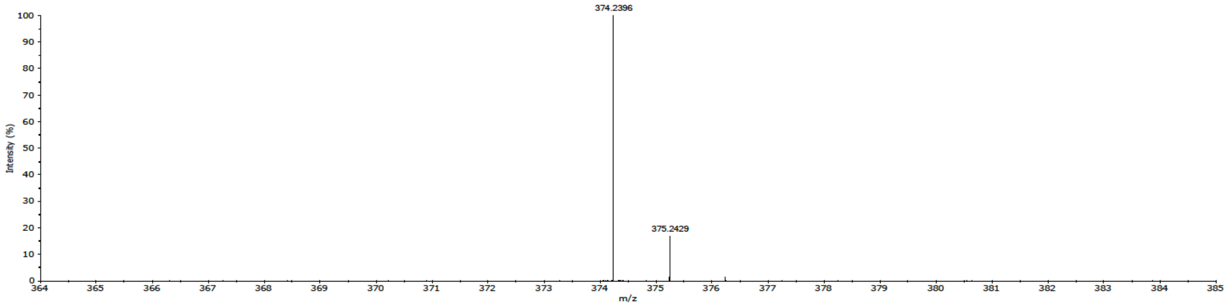

*Tert*-butyl 14-(3-azidopropyl)-1-(4-(6-((5-((2-chloro-6-methylphenyl)carbamoyl)thiazol-2-yl)amino)-2-methylpyrimidin-4-yl)piperazin-1-yl)-4,13-dioxo-3,8,11-trioxa-5,14-diazaheptadecan-17-oate (**5**)

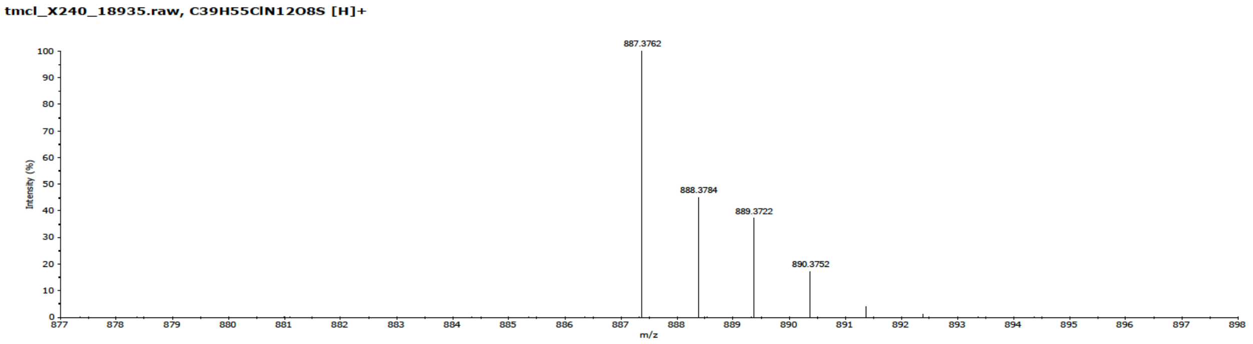


14-(3-Azidopropyl)-1-(4-(6-((5-((2-chloro-6-methylphenyl)carbamoyl)thiazol-2-yl)amino)-2-methylpyrimidin-4-yl)piperazin-1-yl)-4,13-dioxo-3,8,11-trioxa-5,14-diazaheptadecan-17-oic acid (**6**)

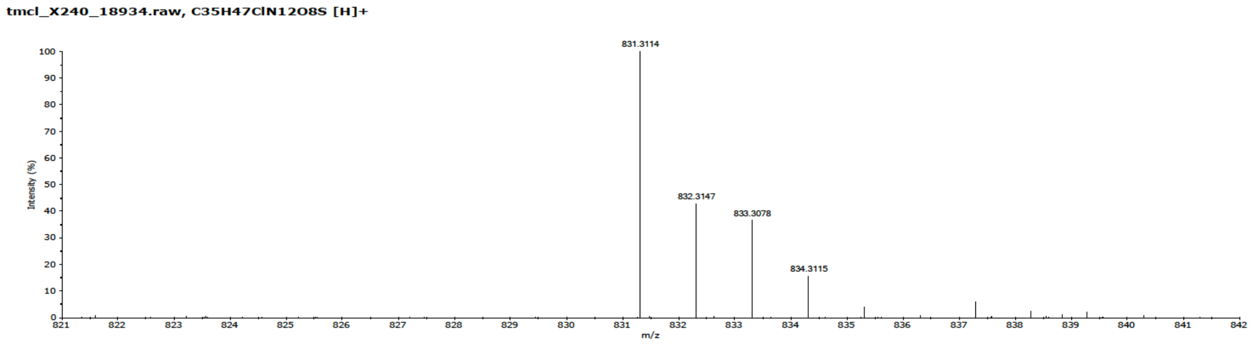


**Das-AI**

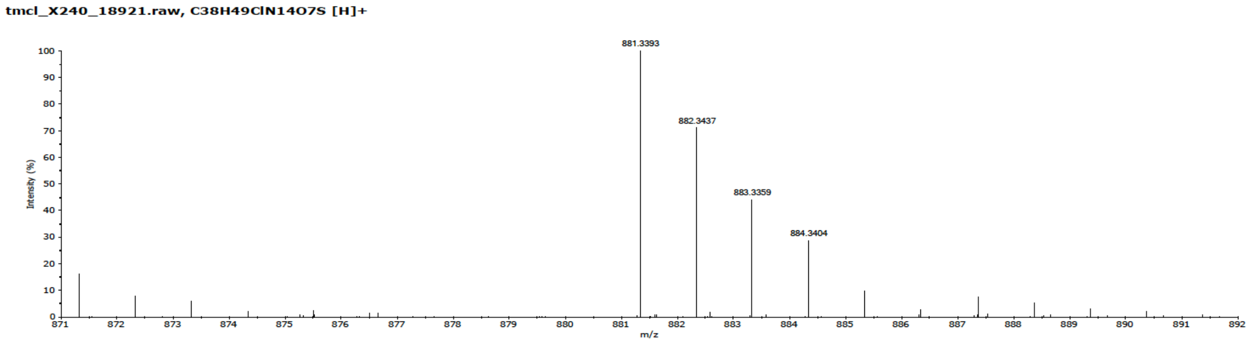

*Tert*-butyl 4-(3-azidopropyl)-22-((7-chloroquinolin-4-yl)amino)-18-ethyl-5,14-dioxo-7,10,15-trioxa-4,13,18-triazatricosanoate (**7**)

tmcl_X240_18931.raw, C35H55ClN8O7 [H]+

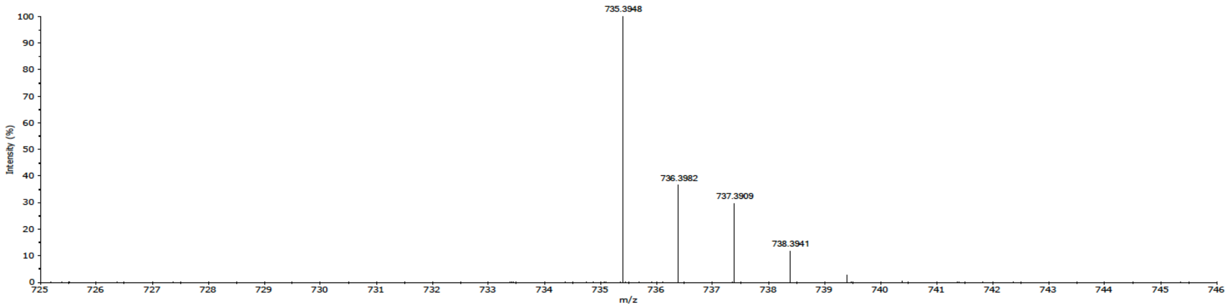


4-(3-Azidopropyl)-22-((7-chloroquinolin-4-yl)amino)-18-ethyl-5,14-dioxo-7,10,15-trioxa-4,13,18-triazatricosanoic acid (**8**)

tmcl_X240_18933.raw, C31H47ClN8O7 [H]+

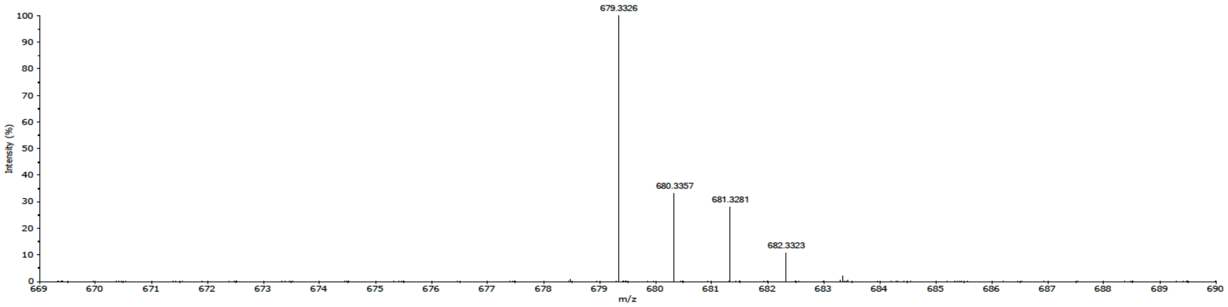


HCQ-AI

tmcl_X240_18919.raw, C34H49ClN10O6 [H]+

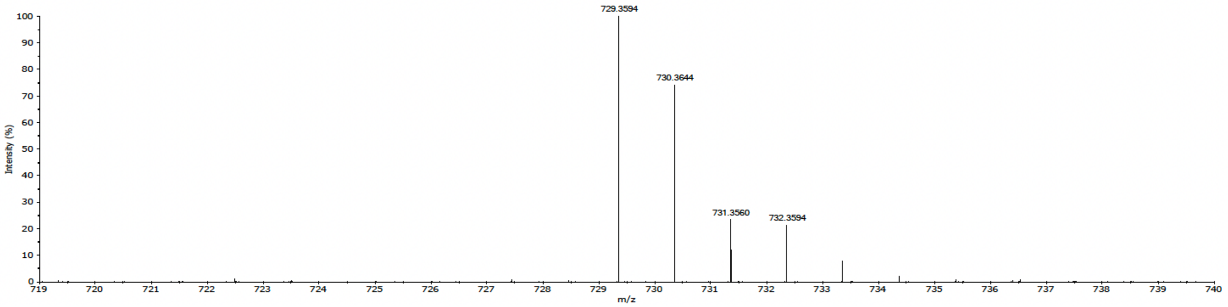

*Tert*-butyl 12-(3-azidopropyl)-1-(9-fluoro-3-methyl-10-(4-methylpiperazin-1-yl)-7-oxo-2,3-dihydro-7*H*-[1,4]oxazino[2,3,4-*ij*]quinolin-6-yl)-1,11-dioxo-5,8-dioxa-2,12-diazapentadecan-15-oate (**9**)

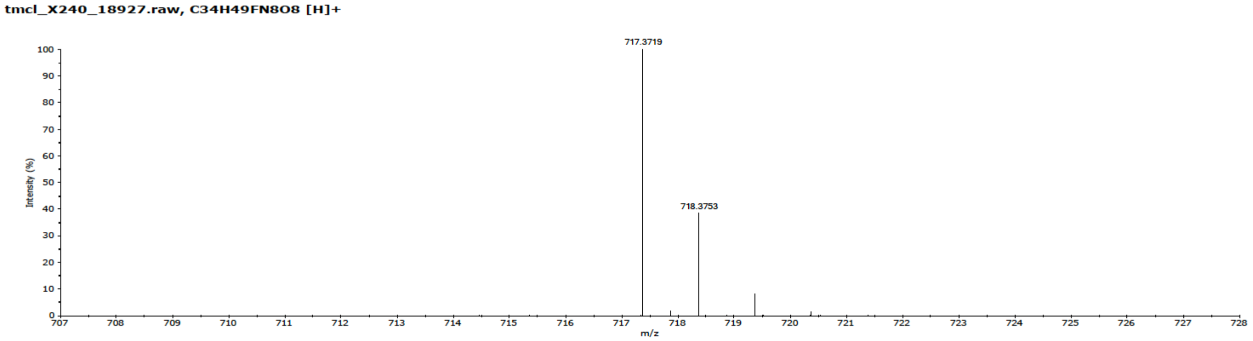


12-(3-Azidopropyl)-1-(9-fluoro-3-methyl-10-(4-methylpiperazin-1-yl)-7-oxo-2,3-dihydro-7*H*-[1,4]oxazino[2,3,4-*ij*]quinolin-6-yl)-1,11-dioxo-5,8-dioxa-2,12-diazapentadecan-15-oic acid (**10**)

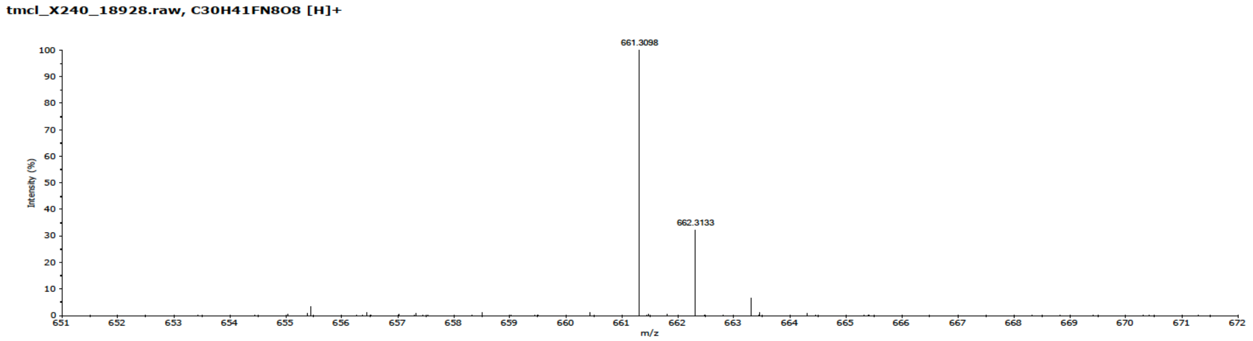


Lev-AI

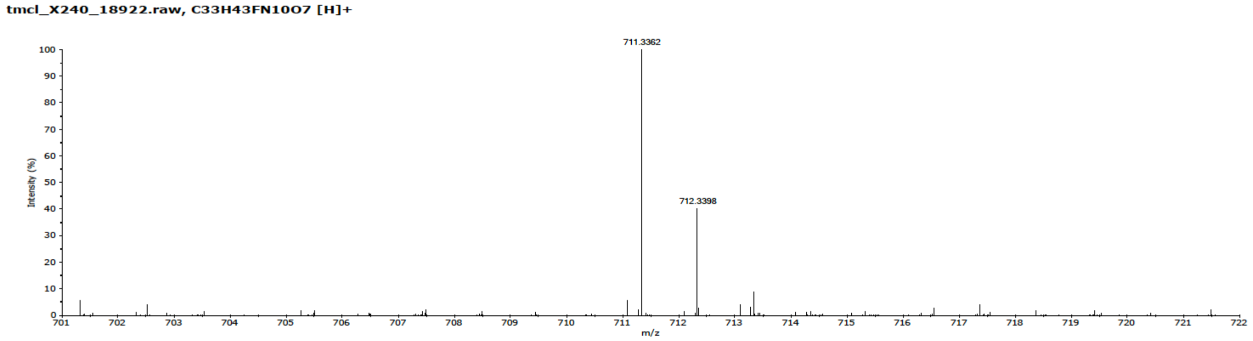

Lev-diazirine

**tmcl_X240_18923.raw, C34H48FN11O7 [H]+**

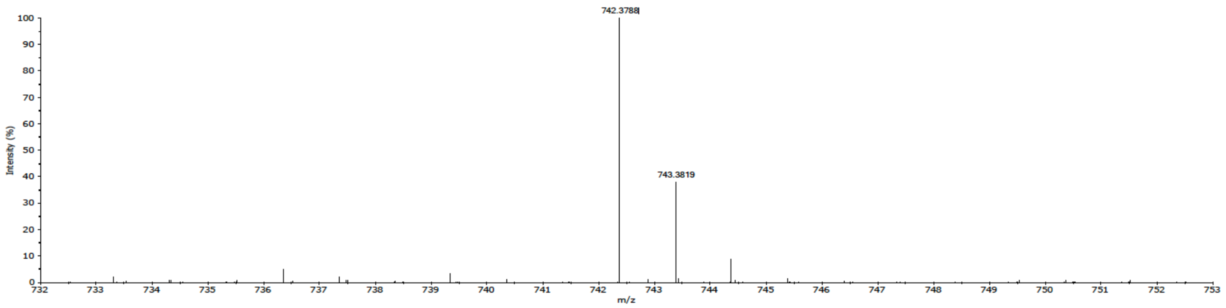


*Tert*-butyl 12-(3-azidopropyl)-2,11-dioxo-6,9-dioxa-3,12-diazapentadecan-15-oate (**11**)

**tmcl_X240_18929.raw, C18H33N5O6 [H]+**

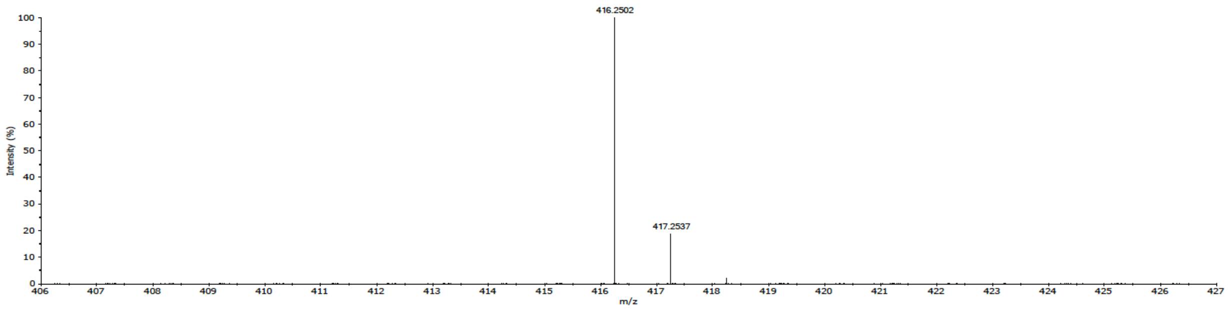


12-(3-azidopropyl)-2,11-dioxo-6,9-dioxa-3,12-diazapentadecan-15-oic acid (**12**)

**tmcl_X240_18930.raw, C14H25N5O6 [H]+**

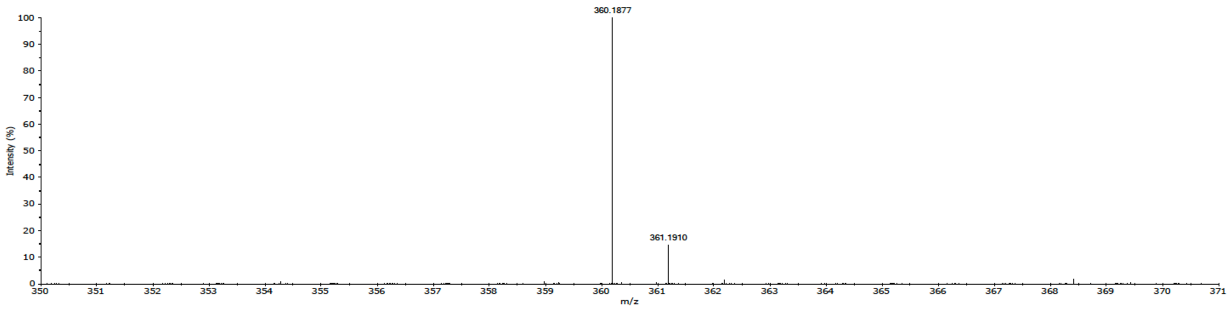


Linker-AI

**tmcl_X240_18920.raw, C17H27N7O5 [H]+**

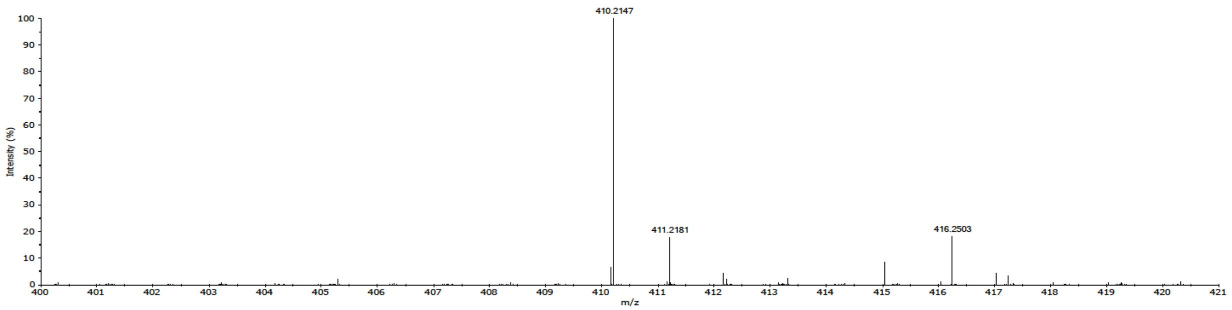
